# Structural Elucidation and Revised Biosynthetic Pathway of the Membrane Vesicle–Associated Antifungal Compound AFC

**DOI:** 10.1101/2025.10.28.685075

**Authors:** Inês Tavares, Yi-Chi Chen, Kirsty Agnoli, Michael Berger, Simon Sieber, Karl Gademann, Leo Eberl

## Abstract

Bacterial membrane vesicles (MVs) serve as delivery vehicles for hydrophobic and membrane-associated secondary metabolites, enhancing their solubility, stability, and bioactivity. Here, we show that the antifungal compound AFC-BC11 (AFC), produced by *Burkholderia cenocepacia* K56-2, is selectively packaged into and released via MVs. Using HR-MS/MS, NMR, and stable isotope feeding experiments, we determined the chemical structure of AFC and analyzed its biosynthesis. Our results confirm that the structure largely matches the recent report by Zhong et al., with a key difference: the double bond in the fatty acid moiety is positioned between C11 and C12. We provide compelling evidence that this constitution reflects the direct incorporation of *cis*-vaccenic acid, the most abundant fatty acid in *B. cenocepacia*, rather than a tailoring modification. Comparative analysis of afcU, afcF, and afcS mutants suggests a biosynthetic pathway involving ω-modification of *cis*-vaccenic acid, revising previous proposals of citric acid conjugation to myristic acid and opening avenues for acyl chain engineering. Together, these findings establish AFC as an MV-associated antifungal metabolite, provide a refined structural and biosynthetic model, and highlight the role of MVs in the dispersal of hydrophobic bioactive compounds.

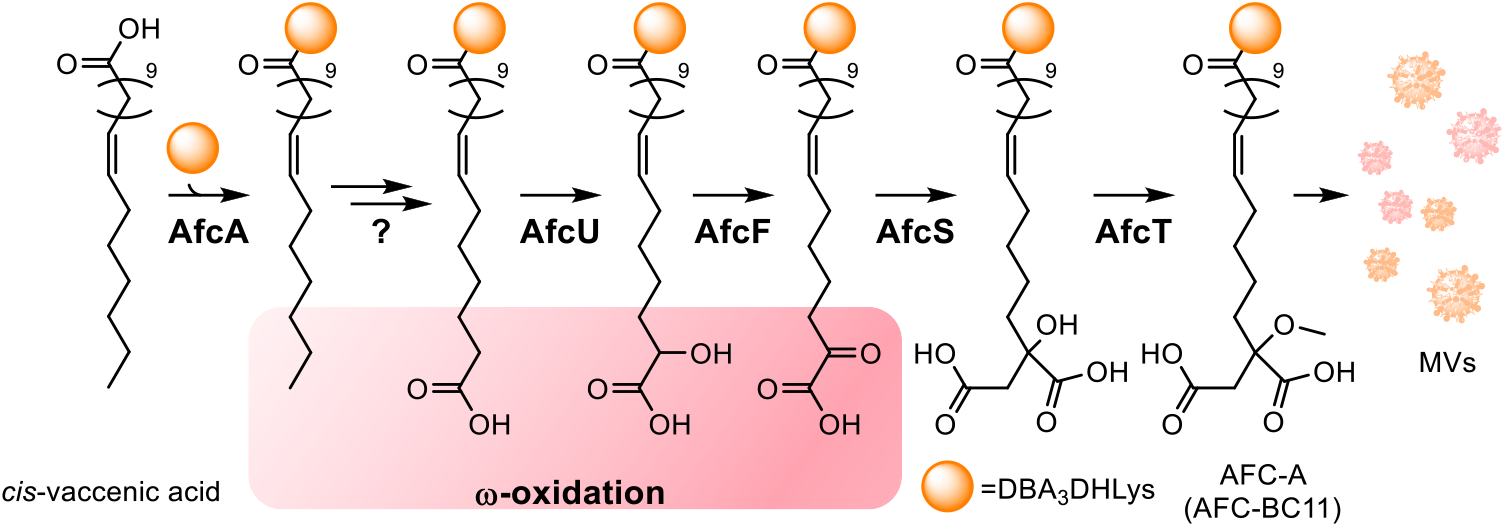

## Introduction

Bacteria employ diverse mechanisms to deliver secondary metabolites to their targets, including protein secretion systems^1^, membrane transporters^2^, and membrane vesicles (MVs)^3^. MVs are small, spherical particles (40-400 nm in diameter) derived from the cell envelope that can carry specific cargo^3^. They arise via different biogenesis routes, such as outer membrane blebbing in Gram-negative bacteria, passage through peptidoglycan pores in Gram-positive bacteria, or cell lysis^3^. MVs have been proposed to function as carriers for membrane-associated or hydrophobic molecules^3^. Indeed, several bacterial secondary metabolites have been shown to be MV-associated (**Fig. 1A**). For example, the *Pseudomonas aeruginosa* quinolone signal (PQS) was shown to intercalate into the outer membrane, inducing curvature and driving MV formation via the “bilayer-couple” mechanism, thereby ensuring their own packaging and transport^4^. Other hydrophobic signaling molecules, such as C16-HSL in *Paracoccus denitrificans*^5^ and CAI-1 in *Vibrio harveyi*^6^, follow a similar principle. Beyond signaling, a wide range of bioactive compounds have been shown to be selectively enriched in MVs, including rhamnolipids and hydroxyalkylquinolines in *Burkholderia thailandensis*^7^, violacein in *Chromobacterium violaceum*^8^, linearmycins in *Streptomyces* sp. Mg165^9^, and the bacteriocin micrococcin P1 in *Staphylococcus hominis*^10^. Strikingly, MV-association can increase solubility, stability, and activity of these compounds, as demonstrated for the algicidal tambjamine LY2, which is packaged into MVs by *Chitinimonas prasina* and efficiently delivered to target microalgae^11^.

**Figure 1.**
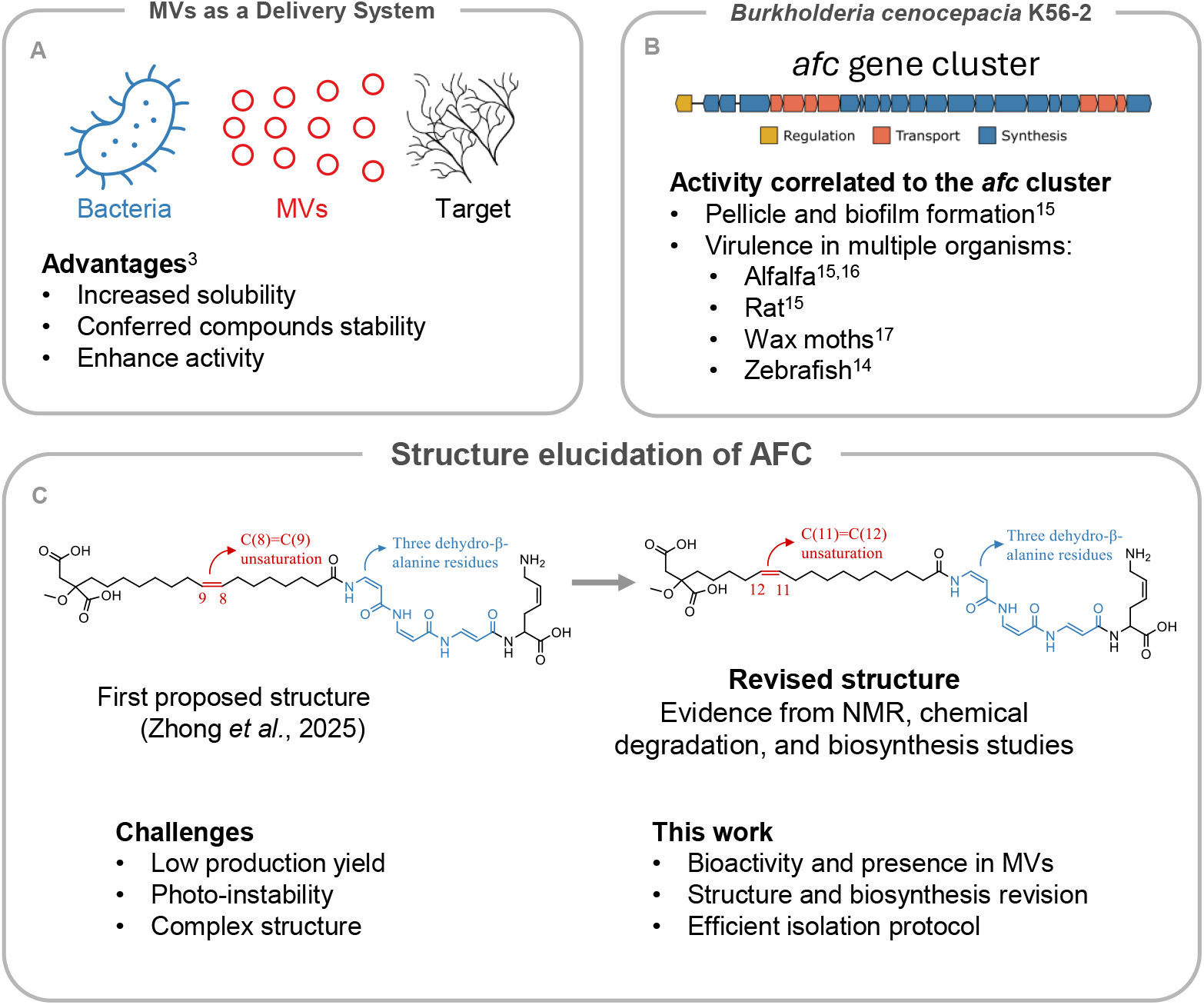
Membrane vesicles, the *afc* gene cluster, and revised AFC structure.

Members of the genus *Burkholderia* are well known for producing a wide range of antifungal compounds^12^. Among them, *Burkholderia cenocepacia* K56-2 produces the antifungal compound AFC-BC11 (AFC-A, **1**), which exhibits activity against diverse plant-pathogenic fungi^13^. Although the chemical structure of AFC was initially unresolved, Kang and colleagues described it as a novel, membrane-associated lipopeptide with limited water solubility in 1998^13^. The corresponding *afc* biosynthetic gene cluster (**Fig. 1B**) has since been linked to pellicle formation, biofilm development, and virulence in multiple model organisms, including alfalfa, rats, wax moths, and zebrafish^14–17^. Notably, in zebrafish, AFC is essential for the transition from intracellular persistence to acute pro-inflammatory infection and its production correlates with increased virulence among closely-related strains^14^.

AFC is produced by several members of the *Burkholderia sensu lato*, a group of closely-related bacteria encompassing seven genera^18^ that inhabit soil, freshwater, industrial environments and a variety of hosts, including plants, fungi, animals and humans^19^. Homologous *afc*-like gene clusters are also found in bacteria outside the *Burkholderia sensu lato*, including in members of the genera *Collimonas, Pseudomonas*, and *Chromobacterium*. Moreover, the *afc* cluster shares gene homology with the bolagladin biosynthetic pathway^20^. While bolagladins display structural similarities to AFC, their synthesis involves a large, modular non-ribosomal peptide synthase that yields a cyclodepsipeptide scaffold absent in AFC (**Fig. S1**).

During a detailed analysis of the cargo of MVs produced by various *Burkholderia* strains, we found that MVs from *B. cenocepacia* K56-2 cultures exhibited strong antifungal activity. We demonstrate here that this activity is linked to AFC, which is specifically associated with MVs and thereby appear to facilitate its dispersal in aqueous environments. We also determined the chemical structure of AFC produced by *B. cenocepacia* K56-2. However, while this manuscript was in preparation, Zhong and colleagues reported on the structure of the AFC from *B. orbicola* and *B. puraquae*^20^ and provided valuable insights into its biosynthesis. Our findings are broadly consistent with their results but also complement them by revealing notable differences in both the structure and biosynthetic pathway of AFC in *B. cenocepacia*.

## Results

### *Burkholderia* sp. release antifungal compounds via MVs

To investigate the role of MV-mediated secretion of bioactive molecules, MVs were isolated from selected *Burkholderia* strains that inhibited the growth of the plant pathogen *R. solani* in dual culture assays on agar plates. The MVs were then tested for antagonistic activity against *R. solani* and *B. subtilis* (**Fig. 2 and S2**). MVs from *B. thailandensis* E264 and *B. cenocepacia* K56-2 showed activity against both organisms, while MVs from *B. vietnamiensis* ABIP434 displayed antifungal activity only. The antifungal activity of MVs from strain K56-2 was of particular interest, as this strain is known to produce AFC-BC11^13^. Given previous descriptions of AFC-

**Figure 2.**
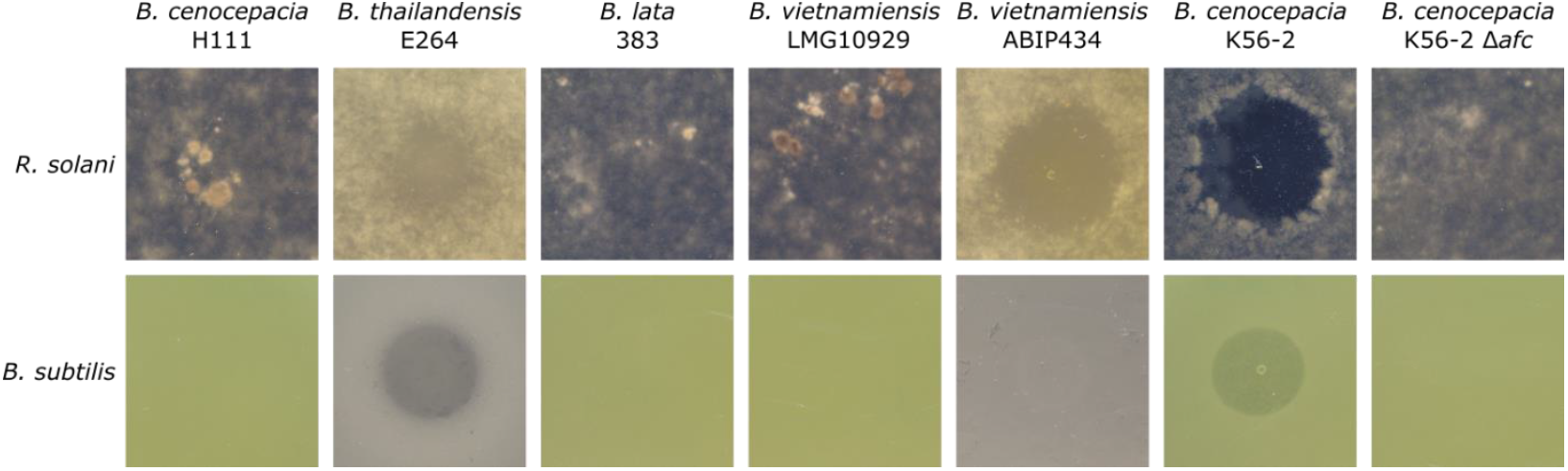
Antagonistic activities of membrane vesicles isolated from selected *Burkholderia* strains against *R. solani* (top) and *B. subtilis* (bottom).

BC11 as a poorly water-soluble molecule with antifungal activity^13^, we hypothesized that the MV-associated activity of K56-2 was attributable to AFC-BC11. To test this, MVs were isolated from both wild-type K56-2 and a mutant strain (Δ*afc*) lacking the entire *afc* biosynthetic cluster and their antifungal activities were compared (**Fig. 2**). The complete loss of antagonistic activity against both organisms in the Δ*afc* mutant suggests that AFC-BC11 was responsible for the activity observed in the MV fraction.

### AFC could be isolated by a novel extraction method

Determining the structure of AFC has been a longstanding challenge, with nearly 25 years elapsing between its initial identification and significant advances in its chemical characterization, despite substantial progress in elucidating its biological activities during this period. The major obstacles in isolating AFC have been its poor solubility in both aqueous and common organic solvents and its instability. To overcome this, we established a novel and straightforward protocol that enabled efficient purification of AFC. In brief, whole K56-2 cultures after 24 h of growth were harvested without centrifugation and directly extracted with chloroform/methanol (1:2). The extract was then centrifuged (4000 × g), and both the aqueous and organic layers were discarded. The resulting precipitate was subsequently extracted with DMF containing 1% formic acid. The yield and purity of AFC using this extraction protocol was compared to the AFC obtained with an alternative protocol described in Zhong *et al*.^20^(**Fig. S3**). Our procedure facilitated the efficient recovery of AFC, and, because both water- and organic-soluble molecules were removed, the crude extract was already of high purity (**Fig. S3**). Final purification was achieved by HPLC for downstream structural and functional analyses.

### Structure elucidation of AFC-A

Purified AFC-A (**1**) was subjected to UHPLC-ESI-HR-timsTOF analysis. A single peak with 734.3965 (Δppm = −0.79) was observed (**Fig. S4A and B**). The HR-MS/MS spectrum displayed major ions corresponding to the fragments in the peptide part of the molecule, generating fragments that are a combination of three dehydro-β-alanine (DBA) and dehydrolysine (DHLys) residues (**Fig. S4C and D**). Interestingly, fragments α, β and γ observed in MS/MS indicated that AFC-A (**1**) might have lost one, two or three DBA moieties, respectively. In addition, the fragmentation of the aliphatic side chain with one DBA subunit (ion b1, 452.2643) was obtained by a pseudo-MS^3^ measurement via Q Exactive orbitrap (**Fig. S5**), showing several fragments with a combination of *O*-methylation, carboxylic acid, or methylcarboxylic acid loss, which confirmed the *O*-methylated malic acid-fatty acid (MMFA) motif.

The structure was further analyzed by a combination of NMR techniques, including 1D and 2D experiments (**Fig. S6-S14, Table S1**). ^1^H – ^1^H TOCSY experiments revealed that the compound was composed of five spin systems, including four modified amino acids and a fatty acid derivative with an unsaturated C=C bond. The connectivity between the different parts of AFC-A (**1**) was assigned using the correlations observed in the ^1^H – ^13^C HMBC and ^1^H – ^1^H NOESY spectra (**Fig. S15**). Further analysis by ^1^H – ^1^H COSY, ^1^H – ^13^C HSQC, ^1^H – ^13^C HMBC experiments led to the identification of a modified lysine residue with a C(4)=C(5) double bond (DHLys) and three DBA units. These amino acids possessed similar chemical shifts to those reported for bolagladins^21,22^. A detailed analysis of the vicinal scalar coupling constants indicated that the double bond of the DHLys residue adopts a cis configuration (^3^*J* = 8.7 Hz), the DBA^1^ fragment is trans configured (^3^*J* = 13.9 Hz), and the DBA^2^ and DBA^3^ units are cis configured (^3^*J* = 8.9 Hz and ^3^*J* = 8.8 Hz, respectively). Structural elucidation of the fatty acid side chain was achieved using a combination of ^1^H – ^1^H COSY, ^1^H – ^1^H TOCSY, ^1^H – ^13^C HSQC, ^1^H – ^13^C HMBC, and ^1^H – ^13^C HSQC-

TOCSY spectroscopy. Notably, the two carbon chemical shifts of C18 and C20 (174.0 and 172.1 ppm) could be identified but not differentiated. To obtain further high quality data, the compound was analyzed on a Bruker AVANCE NEO 1.2 GHz NMR spectrometer. Clear separation between the protons H10 (1.95 ppm) and H13 (1.97 ppm, **Fig. S16-S17**) was observed. In addition, a high-resolution ^1^H – ^13^C HSQC spectrum focusing on the methylene region enabled assignment of C10 and C13 (**Fig. S18**). Finally, correlations observed in the ^1^H – ^13^C HSQC-TOCSY spectra recorded with mixing times of 60 and 100 ms allowed the determination of the proton and carbon atoms at positions 14 and 15 (**Fig. S19-20**). These results were further corroborated by analysis of an AFC-A (**1**) sample obtained from cultures grown in LB medium instead of PDB (see section below). Unlike the PDB-derived samples, the LB samples included an inseparable co-eluting impurity, which facilitated the analysis. A full characterization was also performed on this sample (**Fig. S21-S29, Table S2**) and a significant shift was observed for the proton at position 16 (now at 1.69 ppm, previously 1.57 ppm). This enabled a clear differentiation of the positions 2, 3, and 16, and revealed a distinct correlation in the ^1^H – ^1^H TOCSY spectrum between the protons at position 16 with those at positions 12 and 13, and a ^1^H – ^13^C HSQC-TOCSY spectrum between the proton at position 13 and 15. These results indicated that the double bond is located between carbons 11 and 12. To further investigate the location of the double bond, oxidative cleavage of AFC-A (**1**) was carried out by ozonolysis, followed by oxidative workup with H2O2. The analysis of the cleavage products by UHPLC-HR-MS/MS resulted in the detection of 1-methoxypentane-1,1,5-tricarboxylic acid, which unambiguously determined the location of the double bond between carbon atoms 11 and 12 (**Fig. S30-S33**). The configuration of this double bond was assigned as *cis* based on a vicinal coupling constant of 11.8 Hz, determined from a ^1^H – ^13^C HSQC spectrum acquired without ^13^C decoupling and by selective ^1^H homodecoupling at 1.96 ppm on the AFC pure sample (**Fig. S34-S35**).

In the initial report on the AFC structure, the double bond was assigned between C8 and C9^20^. As that study employed *B. puraquae* DSM03137 as the producing strain, the discrepancy in double bond position was initially attributed to strain variation. To clarify this, AFC was isolated from *B. puraquae* DSM03137 following the protocol of Zhong *et al*. and analyzed by UHPLC-HR-MS/MS. The compound displayed an identical retention time, confirmed by spiking experiments, and indistinguishable fragmentation patterns (**Fig. S36**). In addition, comparative analysis of the ^1^H-NMR and ^1^H-^13^C-HSQC spectra confirmed the identity of the two samples (**Fig. S37-S38**), indicating that the double bond resides between C11 and C12, thereby necessitating a structural reassignment of AFC from *B. puraquae* DSM103137.

### Environmental conditions shape the generation of diverse AFC side products

To evaluate the role of the growth medium in AFC biosynthesis, we profiled the AFCs extracted from the supernatants of K56-2 cultures grown in liquid LB and PDB media. Major AFC side products were identified by their conserved MS/MS fingerprint to the DBA3-DHLys fragments. These variants differed in terminal modifications (carboxylic acid, α-hydroxy carboxylic acid, α-oxo carboxylic acid, or α-amine carboxylic acid) and acyl chain length (Cn:0 or C(n+2):1) (**Fig. 3 and S39-S43**). Comparison of the peak areas showed that PDB medium predominantly supported the production of AFC-A (**1**, 82%) and AFC-B (**2**, 6%). In contrast, LB cultures displayed a more complex profile, with a marked increase in ω-carboxylic fatty acid (ω-CFA)-AFCs, specifically AFC-C (**3**, 26%) and AFC-D (**4**, 19%) (**Fig. 4**). Additional minor derivatives were also detected, including the α-hydroxy-CFA-AFCs AFC-E (**5**) and AFC-F (**6**), the α-oxo-CFA-AFCs AFC-G (**7**) and AFC-H (**8**), and the α-amino-CFA-AFCs AFC-I (**9**) and AFC-J (**10**), with structures predicted by MS/MS fragmentation (**Fig. S39-S43**).

**Figure 3.**
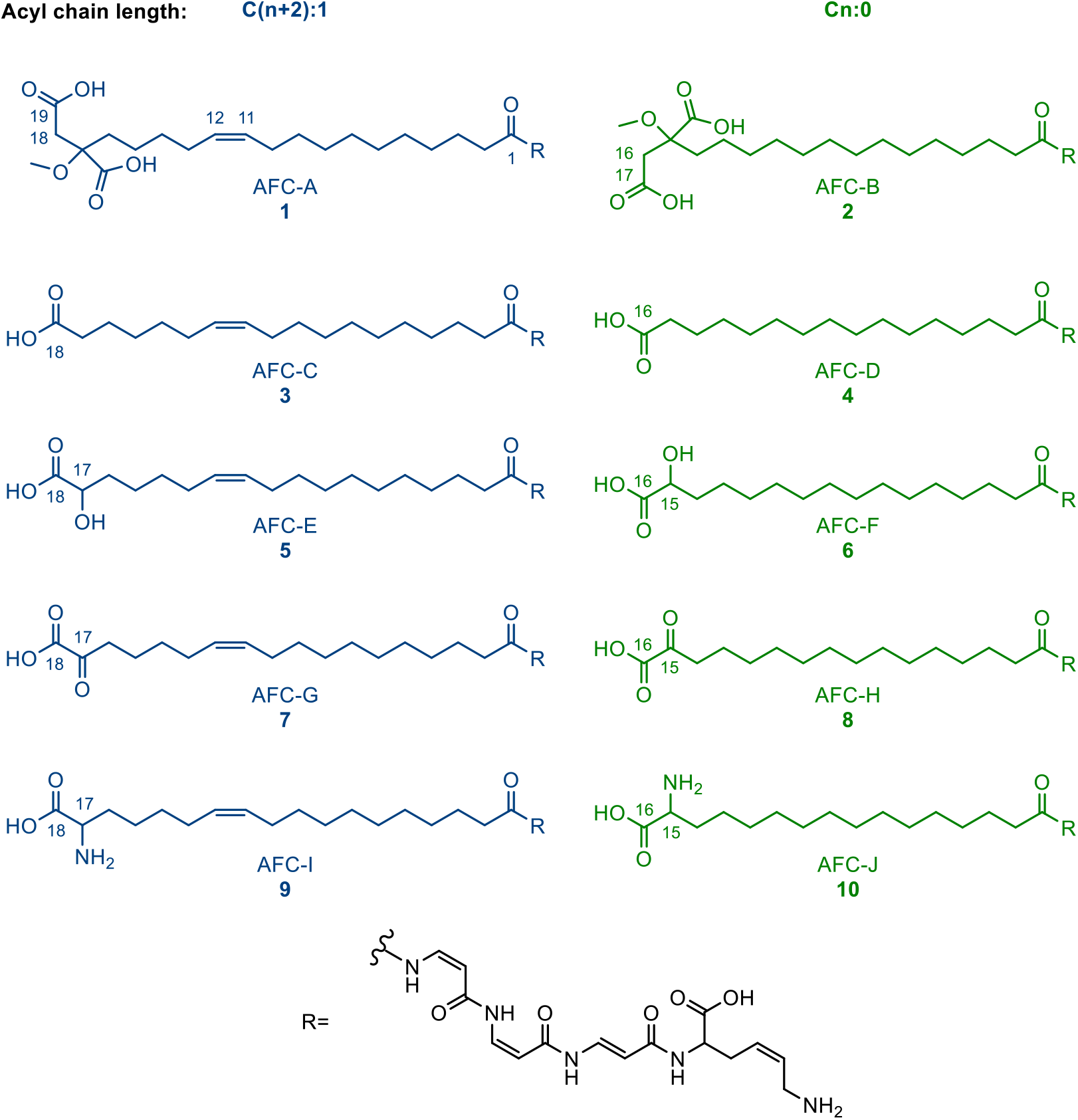
Structure of major AFC products. All products share the same core structure except for the acyl chain composition and the modification on the terminus. AFCs with either acyl chain length C(n+2):1 or Cn:0 are grouped and labeled in blue or green, respectively.

**Figure 4.**
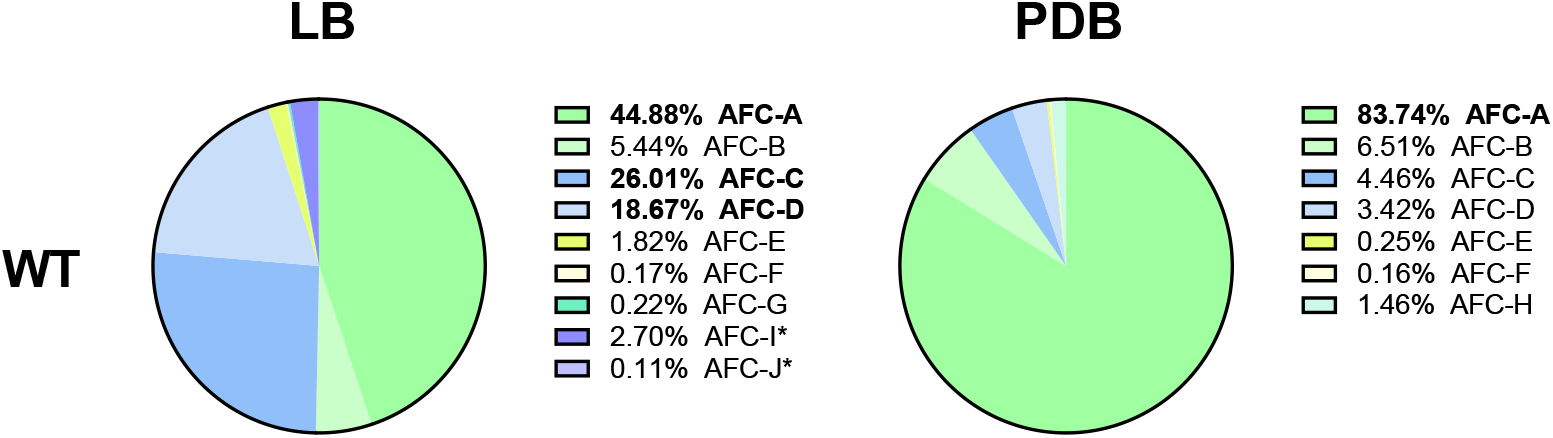
The ratio of ion abundance of major AFC products in LB and PDB cultures. Peak area of ion abundance from AFCs (**1** to **10**) with 1^+^ and 2^+^ charges were integrated. Compounds with more than 10% abundance are shown in bold. *AFC-I and AFC-J have major ions of 2^+^, the exact quantity may be different to the other AFCs.

**Figure 5.**
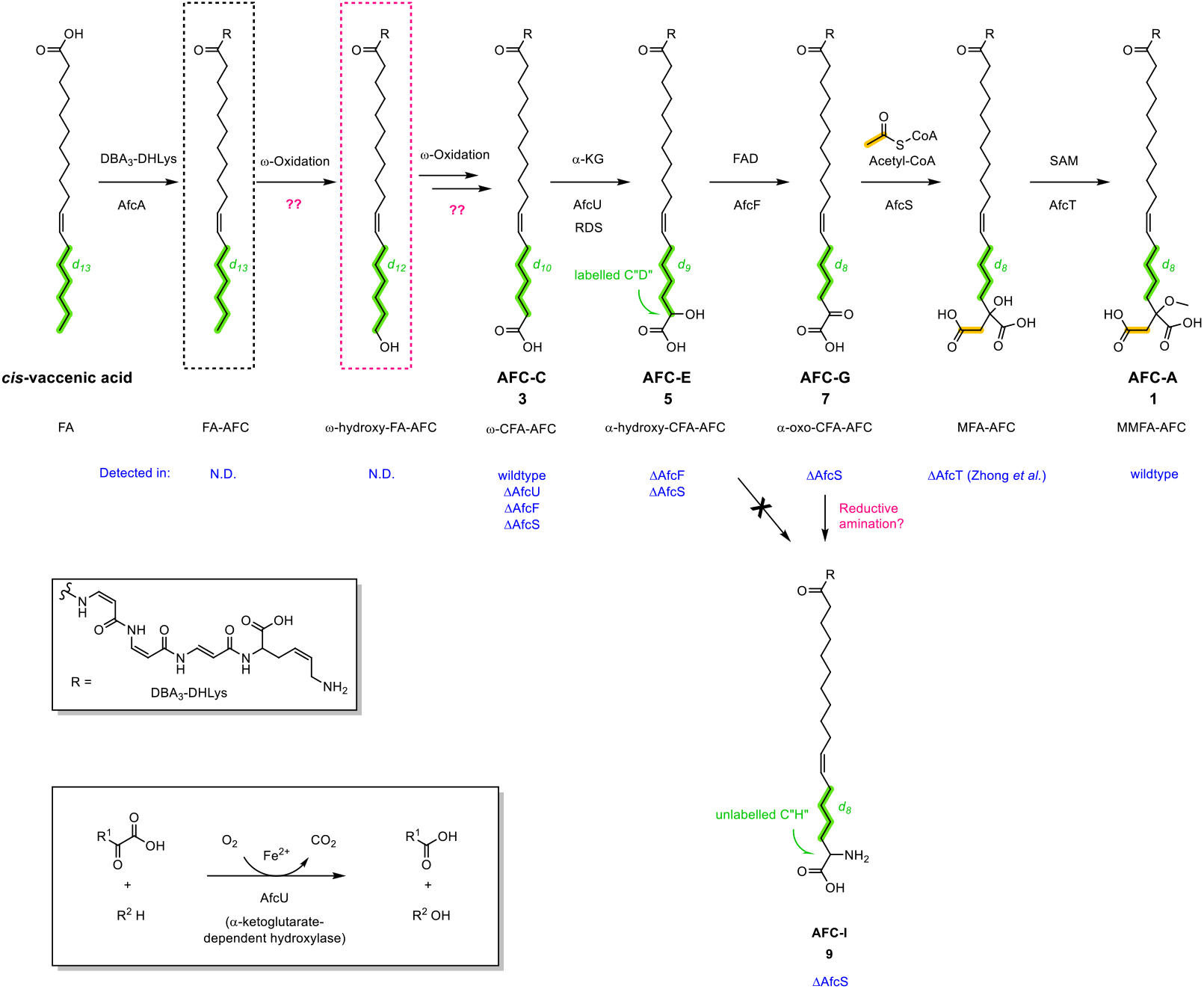
Proposed biosynthetic pathway of AFCs with fatty acid ω-modification. The biosynthesis of AFCs was clarified by using deuterium-labeled (green) *cis*-vaccenic acid. Possible AfcU catalytic mechanism shown in the bottom box is proposed by comparing the reaction with TauD from *E. coli* K12 and *P. putida* KT2442. Intermediates not observed are labeled in dotted lines. Strains that generated the correlated products are labeled in blue. Undetermined reactions and intermediates are labeled in red. Predicted acetyl-CoA addition is labeled in orange. FA, fatty acid; MFA, malic acid-fatty acid.

### Biosynthesis of the AFC fatty acid moiety

It has been proposed that the fatty acid side chain in AFC-A (**1**) is incorporated by initial conjugation to the DBA moiety via an amide bond, followed by terminal modification through citric acid conjugation^20^. In contrast, the biosynthesis of bolagladins is thought to involve elongation of the fatty acid side chain via fatty acid synthase (FAS) activity directly on the *O*-methylated citrate core^21,22^ (**Fig. S44**). To investigate substrate specificity and clarify the biosynthetic route, stable isotope labeling experiments were conducted using deuterium-labeled fatty acid precursors in PDB culture, and their correlated products were analyzed by UHPLC-HR-MS (**Table S3**, compounds **11-16**). While in vitro assays had previously indicated that AfcA shows highest activity toward myristic acid^20^, MS analysis of cultures supplemented with deuterium-labeled myristic acid (myristic acid-*d*_*27*_) revealed no incorporation into AFC-A (**Fig. S45**). Although a peak corresponding to unlabeled AFC-A (**1**) was detected, its labeled counterpart (AFC-A-*d24*) was absent. Instead, ω-CFA-AFC derivatives with ω-oxidized myristic acid (**11)** or ω-oxidized myristic acid-*d24* (**12)** were detected. In addition, trace amounts of shorter MMFA-AFC analogs **13** and **14** were observed (**Fig. S45**). No α-hydroxy carboxy myristic acid-AFCs were detected (**15** and **16**). This suggests that myristic acid is accepted by AfcA but undergoes ω-oxidation without efficient downstream processing into MMFA-AFCs, indicating poor compatibility of myristic acid with the canonical AFC biosynthetic machinery.

In contrast, supplementation with palmitic acid (C16:0) and *cis*-vaccenic acid (C18:1ω7) resulted in the successful production of MMFA-AFCs (**Fig. S46-S47**). Deuterium-labeled palmitic acid (palmitic acid-*d31*) led to the detection of an intermediate ω-CFA-AFC (AFC-D-*d28*, **18**) and the final product AFC-B-*d26* (**17**), which eluted at the same retention time as unlabeled AFC-B (**2**) (**Fig. S46**). The abundance of AFC-B-*d26* (**17**) increased with higher concentrations of labeled substrate, supporting conversion from ω-CFA-AFC to MMFA-AFC. Similarly, treatment with *cis*-vaccenic acid-*d13* resulted in the incorporation of deuterium into AFC-A-*d8* (**19**) (**Fig. S47**), consistent with the AfcA-mediated direct incorporation model proposed by Zhong *et al*.^20^ Importantly, *cis*-vaccenic acid appears to be the most efficiently incorporated substrate, which aligns with the identity of AFC-A (**1**) as the major product featuring a *cis*-vaccenic acid side chain with a double bond at C(11)=C(12). When the medium was supplemented with stearic acid (C18:0, unlabeled), *O*-methylated malic acid-stearic acid-AFC (**21**) could be identified by MS/MS without a significant increase (**Fig. S48**). This result demonstrates that stearic acid can be incorporated into the final AFC product. However, its uptake from the culture medium is inefficient due to low solubility. In summary, *cis*-vaccenic acid appears to be the most efficiently incorporated substrate, consistent with the identification of AFC-A (**1**) as the major product featuring a *cis*-vaccenic acid side chain with a double bond at C(11)=C(12).

### AfcS, AfcF, and AfcU are involved in fatty acid moiety modification

To further investigate the biosynthesis of the fatty acid moiety, we generated deletion mutants of several genes in the *afc* cluster and analyzed their metabolic products by UHPLC-HR-MS, along with their antifungal activity (**Table S4**). Quantification of AFC derivatives (**Figures S49-S53**) revealed that the main metabolites detected in the Δ*afcS* mutant varied in the length of their acyl side chain, being either based on C16:0 or C18:1 and lacked the citric acid moiety. Instead, the aliphatic side chain terminated either with a carboxylic acid (AFC-C (**3**) and AFC-D (**4**)), a α-hydroxy carboxylic acid (AFC-E (**5**) and AFC-F (**6**)), a α-oxo carboxylic acid (AFC-G (**7**) and AFC-H (**8**)), or a α-amino carboxylic acid (AFC-I (**9**) and AFC-J (**10**)). These results are consistent with a recent study by Zhong *et al*., which showed that AFC-C (**3**) and AFC-E (**5**) were produced both by the wildtype and the Δ*afcS* mutant^20^. Similar findings were reported for bolagladin biosynthesis^21,22^: mutation of *bolR*, the homolog of *afcS*, likewise resulted in truncated AFC derivatives possessing either a carboxylic acid or a α-hydroxy carboxylic acid group at the terminal position of the acyl side chain. In the Δ*afcS* mutant, α-oxo-CFA-AFCs (**7** and **8**) and α-amino-CFA-AFCs (**9** and **10**) were also detected, suggesting that disruption of acetylation in this mutant leads to the accumulation of α-oxo-CFA-AFCs (**Fig. S50**), which may subsequently be converted into α-amino-CFA-AFCs, possibly by reductive amination.

Metabolic profiling of the *afcF* mutant revealed it produced derivatives terminating in either a carboxylic acid (**3** and **4**) or a α-hydroxy carboxylic acid (**5** and **6**) at the end of the acyl chain. These two compounds were likewise observed in the Δ*afcF* mutant of *B. pyrrocinia* DSM 10685^20^. Since the α-oxo-CFA-AFCs (**7** and **8**) were observed only in Δ*afcS* and not in Δ*afcF*, we hypothesize that AfcF catalyzes the oxidation of the α-hydroxy-CFA-AFCs into the α-oxo-CFA-AFCs. The Δ*afcU* mutant produced only ω-CFA-AFCs (**3** and **4**), strongly suggesting its role in the initial modification step of the ω-CFA (**Fig. S50**). Based on these findings, we propose that biosynthesis of the AFC fatty acid moiety proceeds via ω-modification of the fatty acid, a pathway distinct from the previously suggested routes reported for AFC and bolaglandins (**Fig. S44 and Fig. 4**)^20–22^.

To test this model, we supplied deuterium-labeled and unlabeled *cis*-vaccenic acids and tracked the intermediates in the wildtype and the Δ*afcU*, Δ*afcF*, and Δ*afcS* mutants (**Fig. S54**). Several deuterium-labeled adducts, including AFC-A-*d8* (**19**), AFC-C-*d10* (**20**), AFC-E-*d9* (**22**), AFC-G-*d8* (**23**), and AFC-I-*d8* (**24**), were detected (**Fig. S54-S55, Table S3**). A stepwise decrease in deuterium labeling was observed from the substrate *cis*-vaccenic acid-*d13* to the final product AFC-A-*d8* (**19**). After the incorporation of the DBA3-DHLys domain via AfcA, the fatty acid terminus was oxidized to form AFC-C-*d10* (**20**), presumably by a FAD-dependent dehydrogenase as suggested by Zhong *et al*.^20^, (**Fig. 4**). No intermediates of the ω-oxidation carrying an OH or CHO group were detected, likely due to rapid turnover or instability. Subsequently, AfcU was demonstrated to introduce a α-hydroxy group, evidenced by the loss of one deuterium atom from AFC-C-*d10* (**20**) to AFC-E-*d9* (**22**) (**Fig. 4 and S44**). Further oxidation by AfcF converted the OH group into the corresponding carbonyl group, with an additional deuterium loss to yield AFC-G-*d8* (**23**) (**Fig. 4 and S44**). The aminated product AFC-I-*d8* (**24**) was observed in the *afcS* mutant but not in *afcF*, indicating that the α-amino group is derived from the α-oxo group rather than the α-hydroxy group, likely via reductive amination catalyzed by an unknown enzyme (**Fig. 4 and S44**). Finally, since α-oxo-CFA-AFC is a structural mimic of oxaloacetate, the carboxymethyl group was likely incorporated from acetyl-CoA via AfcS (**Fig. 4**). These findings support our hypothesis that AFC biosynthesis proceeds via ω-modification. As further evidence, supplementation of the Δ*afcS* mutant with malic acid mimics (mercaptosuccinic acid and bromosuccinic acid) failed to yield corresponding labeled AFC products (**Fig. S56-S57**).

## Discussion

While the major steps in the assembly of the peptide components of both AFC and bolagladins have been well resolved, the exact chemistry underlying the addition and modification of the fatty acid remains enigmatic^20–22^. Apart from the additional 2,3-double bond, the fatty acid side chains of AFC and bolagladins are structurally identical, and the enzymes involved in their synthesis display a high degree of homology (53-84%). Nevertheless, two fundamentally different biosynthetic pathways have been proposed for the formation of the fatty acid moieties in AFC and bolagladins.

In the case of bolagladins, it has been suggested that the fatty acid residue originates from an unusual citryl-CoA starter unit, which is elongated by a malonyl group attached to the primary metabolic fatty acid synthase (FAS) acyl carrier protein (ACP). Subsequent processing of the β-ketothioester, resulting from the elongation of citryl-CoA (or *O*-methyl-citryl-CoA) with malonyl-ACP by the primary metabolic FAS, would yield the saturated *O*-methyl-citrate-primed fatty acyl-ACP thioester. Finally, the 11,12- and 2,3-double bonds are introduced into this thioester by the putative fatty acid and acyl-ACP desaturases BolF and BolL, respectively^21,22^ (**Fig. S44**). In the case of AFC, it has been proposed that myristic acid is activated by AfcA and subsequently transferred to the terminal β-amino group of DBA3-DHLys^20^ (**Fig. S44**). The fatty acid is then conjugated to citric acid by undetermined enzymes to generate the final MMFA-AFC product. However, the mechanism responsible for introducing the double bond in the fatty acid has remained unresolved.

Our data supports the idea that AfcA transfers a fatty acid to DBA3-DHLys, but we propose that *cis*-vaccenic acid, rather than myristic acid, serves as the precursor. This conclusion is supported by both our feeding experiments and also by the fact that *cis*-vaccenic acid is the most abundant fatty acid in *B. cenocepacia*^23,24^. Notably, this would also explain the C(11)=C(12) double bond in the fatty acid residue without requiring an additional enzyme to introduce it. The position of this double bond is identical to that in bolagladins, suggesting that *cis*-vaccenic acid may likewise serve as a precursor for their biosynthesis. Substrate assays with purified AfcA showed the highest enzymatic activity with myristic acid^20^. However, *cis*-vaccenic acid was not included among the tested substrates. Since myristic acid is present only in trace amounts in *Burkholderia sensu lato*^25^, our data suggest that this fatty acid is taken up by the cell and rapidly converted into ω-carboxylic myristic acid-AFC. Yet, this precursor appears to be suboptimal for further modification by downstream tailoring enzymes, resulting in the accumulation of this intermediate. The α-ketoglutarate (αKG)-dependent hydroxylase AfcU appears to represent the rate-determining step with high selectivity for ω-CFA-AFC substrates. This is supported both by the absence of α-hydroxy carboxylic myristic acid-AFCs in myristic acid-treated cultures and by the increased levels of ω-CFA-AFCs observed in LB medium. In contrast, supplementation with palmitic or *cis*-vaccenic acid did not lead to ω-CFA accumulation, suggesting that these fatty acids provide the optimal chain length and structural features required for efficient downstream processing in AFC biosynthesis.

Notably, *cis*-vaccenic acid (C18:1ω7) is not only the most abundant fatty acid in *B. cenocepacia*^26^ but also in most *Burkholderia sensu lato* strains, where it is considered a diagnostic marker^25^. The variety of AFC derivatives observed in LB but not in PDB medium may reflect changes in the cellular fatty acid profile, thereby providing a broader spectrum of precursors for AFC biosynthesis. Interestingly, previous work has linked the *afc* biosynthetic cluster to lipid metabolic pathways^26^. Specifically, mutations in *afcE* and *afcF* were shown to alter the overall lipid composition, leading to an increased abundance of *cis*-vaccenic acid in the cell^26^, possibly indicating an accumulation of this fatty acid when it is no longer utilized for AFC biosynthesis. Taken together, our findings support the notion that AFC biosynthesis directly incorporates fatty acids from primary metabolism, which appears to be influenced by growth conditions.

The greatest remaining mystery in the biosynthesis of both AFC and bolagladins concerns the introduction of the citrate group into the fatty acyl chain. For bolagladin biosynthesis, it has been proposed that citryl-CoA serves as the starter unit for assembly of the unusual fatty acid residue incorporated into the molecule^21,22^. In contrast, for AFC, it has been suggested that a citryl-CoA precursor couples to the terminal methyl group of a myristoyl intermediate^20^. Based on detailed analysis of AFC derivatives and various feeding experiments, we propose an alternative biosynthetic pathway for AFC production. In this model, biosynthesis is initiated by ω-oxidation of *cis*-vaccenic acid-AFC, followed by another oxidation step catalyzed by AfcU (**Fig. 4**). Subsequent oxidation by AfcF, acetylation by AfcS, and methylation by AfcT would then complete the maturation of the terminus of the fatty acid moiety. This pathway would neither require the biosynthesis of an unusual citryl-CoA starter unit nor the ill-defined coupling of citryl-CoA to the terminal alkyl group of a myristoyl intermediate, nor an additional desaturase for the introduction of the double bond.

## ASSOCIATED CONTENT

### Supporting Information

Detailed Material and Methods; supplementary tables including compound list, antifungal activity test, NMR chemical shift summary, bacterial strains and primers used; supplementary figures including antifungal activity test, gene alignment, chromatogram and MS pattern, MS/MS fragment assignment, NMR spectra, chemical shift assignment, ozonolysis, possible biosynthetic pathway, quantification, molecular networking (PDF)

### Author Contributions

^‡^ Inês Tavares and Yi-Chi Chen contributed equally to this work. The manuscript was prepared with contributions from all authors. All authors have read and approved the final version of the manuscript.

### Funding Sources

Swiss National Science Foundation (310030_192800, 212603)

## ACKNOWLEDGMENT

The authors thank the Swiss National Science Foundation (310030_192800212603) for financial support. We thank Simon Jurt (UZH) for strong support with NMR spectroscopy. We would like to thank Luca Bürgi and Prof. Dr. Laurent Bigler from the Department of Chemistry, UZH, for their support with the use of the Bruker timsTOF and Thermo QExactive instruments.

## Notes

### Competing Interest Statement

The authors have declared no competing interest.

