## Supporting Information for "Structural Elucidation and Revised Biosynthetic Pathway of the Membrane Vesicle–Associated Antifungal Compound AFC"

### Materials and Methods

#### Bacterial strains, plasmids and materials

Bacterial strains are listed in **Table S5** and plasmids in **Table S6**. Unless otherwise stated, strains were routinely cultured in Lysogeny Broth (Difco™ LB Broth, Lennox) or Potato Dextrose Broth (PDB, Difco™) at 37 °C with shaking. *B. thailandensis* was grown in LB without salt (LBNS) at 30 °C and *B. vietnamiensis* was grown in LBNS at 37 °C. Antifungal assays were done with Potato Dextrose Agar (PDA, Difco™) and Malt Extract Agar (MEA, Difco™).

*Pseudomonas* Isolation Agar (PIA, Difco™) and LB were used for selection of *Burkholderia* mutants or transconjugants. When necessary, antibiotics were used at the following concentrations: for *E. coli* 10 µg/mL tetracycline (Tc), 50 µg/mL trimethoprim (Tp), 25 µg/mL kanamycin (Km); for *B. cenocepacia* 100 µg/mL Tc, 150 µg/mL Tp, 50 µg/mL gentamycin (Gm).

#### Mutant construction

Mutants were constructed as described by Flannagan and colleagues[1] using the SceI mutagenesis protocol. Briefly, two homologous regions upstream and downstream of the region to delete were amplified and cloned into pGPI-SceI. The pGPI constructs were conjugated into *B. cenocepacia* K56-2 by triparental mating using the pRK2013 helper plasmid, and the plasmid was integrated into the genome after a single crossover event. Strains with successful incorporation of pGPI were selected on PIA Tp. After this step, the plasmid pDAI-SceI was conjugated into the new strain, causing a double stranded break in its genome. Successful recombination to repair the DNA damage and excision of pGPI resulted in resistant clones on LB Tc Gm. Primers used for mutant construction have been listed in **Table S7**.

### *afc* cluster deletion

*B. cenocepacia* strain K56-2Δ*afcA* was constructed as follows: pSHAFT2-FRT-afcKOUP was constructed by amplifying an 1,334-bp fragment of *B. cenocepacia* pC3 using primers upXhoF and upBglIIR, digested with XhoI and BglII, and inserted into these sites within pSHAFT2-FRT. This plasmid was transferred into K56-2 by conjugation, and single-crossover insertion mutants were selected on PIA supplemented with chloramphenicol and verified by sequencing across the joins between pC3 and the pSHAFT2-FRT-afcUP. pEX18Tp-FRT-afcKODOWN was constructed by amplification of a 1,256-bp fragment of pC3 using primers afcKOdownPstF and afcKOdownBamR followed by digestion of this fragment with BamHI and PstI, and insertion into pEX18Tp-FRT digested with the same enzymes. This vector was transferred to K56-2 bearing the single-crossover insertion of pSHAFT2-FRT-afcKOUP, and insertion mutants were selected on PIA containing trimethoprim. Correct insertion was verified by PCR across the pC3-pEX18Tp-FRT-afcKODOWN joins. Excision of the *afc* cluster was achieved by the introduction of pBBR5::FLP, as described in [2]. The *shvR* gene was not deleted.

### Construction of Δ*rep* and Δ*afcF* mutants

Since the *afc* cluster in K56-2 contains a duplication of an internal region, resulting in two copies of genes *afcE* and *afcS* and partial copies of genes *afcR* and *afcT*, this region was deleted in order to obtain the K56-2 Δ*rep* strain without duplications and with a functional *afc* cluster. This strain was used to construct mutants Δ*afcR* and Δ*afcS.*

Gene *afcF* was deleted in the Δ*afcR* strain background, with *afcR* supplied *in trans* using pBBR-afcR.

**MV isolation and quantification**

Cell cultures were pelleted at 22,000 × *g* for 10 min and the supernatant was filtered with a 0.45 μm PVDF membrane (Durapore Membrane Filter, Merck). The supernatant was ultracentrifuged at 150,000 × *g* at 4 °C for 1.5 h and the MV pellet was washed with Milli-Q $H_2O$ and reharvested using the same conditions. MVs were resuspended in appropriate volumes of Milli-Q $H_2O$, depending on pellet size. Protein content was quantified with the Pierce™ BCA Protein Assay Kit. Lipid content was measured with the FM1-43 dye (Life Technologies, USA).

**Bacterial culture and substrate supplementation for AFC production**

All fatty acids, including myristic acid (Sigma), palmitic acid (Sigma), stearic acid (Sigma), *cis*-vaccenic acid (Sigma), myristic acid-$d_{27}$ (Sigma), palmitic acid-$d_{31}$ (Sigma), *cis*-vaccenic acid-$d_{13}$ (Cayman Chemical) were dissolved in DMSO. For malic acid mimicking substrates, mercaptosuccinic acid (Sigma) and bromosuccinic acid (Sigma) solutions were prepared in $H_2O$ and the pH was adjusted to 7. Fatty acids or malic acid mimics were supplemented with final concentration of 1 mM or 5 mM, respectively. K56-2 wildtype or mutant strains were inoculated in 2-5 mL PDB with OD 0.01 with or without supplements for 24 h in 37 °C. Cells were extracted by adding $CHCl_3$:MeOH=1:2 with the same amount as the culture volume. The mixture was extracted vigorously and subsequently centrifuged (10 min at 2,500 × *g*) to separate the phases. The bottom chloroform layer and the top aqueous layer were removed, and the middle precipitate layer was resuspended in DMF acidified with 1% formic acid with vortexing and sonication (Branson 2210 sonicator) for 10 min. After centrifugation (10 min at 21,300 × *g*), the supernatants were then stored at 4 °C for further analysis. For comparison with the method of Zhong *et al,* 2025[3], both cultures were grown in PDB for 96 h at 37°C. All samples for quantification and method comparison were prepared with 3 replicates.

#### Antifungal and antibacterial activity assays

Dual culture assays were performed against *R. solani* (obtained from the culture collection of the phytopathology group of the Institute of Plant Sciences, Federal Institute of Technology, Zurich, Switzerland). These were carried out as detailed by O'Grady and colleagues[4] on PDA or MEA plates. For antagonistic activity against *B. subtilis,* 30 to 50 µL of MVs was spotted on top of a lawn of *B. subtilis.* The plates were incubated overnight at 37 °C. For antagonistic activity against *R. solani*, 30 to 50 µL of MVs were spotted on top of MEA plates. After the MV drops were dry, a suspension of disrupted mycelia from *R. solani* was sprayed on top, as described by Jenul *et. al*[5]. The plates were incubated for 2 days at room temperature in the dark.

#### AFC purification

*B. cenocepacia* K56-2 overnight cultures were used to inoculate PDB subcultures at $OD_{600}$ 0.01, which were grown for 24 h at 37 °C. Cells were extracted by adding $CHCl_3$:MeOH=1:2 with the same amount as the culture volume. The mixture was extracted vigorously and subsequently centrifuged (10 min at 21,300 × *g*) to separate the phases. The bottom chloroform layer and the top aqueous layer were removed, and the middle precipitate layer was resuspended in DMF acidified with 1% formic acid with vortexing and sonication (Branson 2210 sonicator) for 10 min. The suspension was centrifuged (10 min at 21,300 × *g*) and the supernatant was injected into an UltiMate™ 3000 UHPLC (ThermoScientific, USA) equipped with a reverse phase C18, 4.6 mm x 250 mm, 5.0 µm, 130 Å chromatography column (Waters Xselect CSH). The column was eluted with A: $H_2O$ + 0.05% formic acid and B : MeCN + 0.05% formic acid with a flow rate of 1 mL/min at 28°C and with the following gradient: 0-2 min, 37% B; 2-27 min, 37%-39% B; 27-27.1 min, 39%-100% B; 27.1-33 min, 100% B; 33-33.1 min, 100%-37% B; 33.1-35 min, 37% B. AFC elution was tracked by UV absorption at 320 nm and fluorescence ($\lambda_{ext}$ = 330 nm, $\lambda_{em}$ = 410 nm) by RS Diode Array Detector and RS Fluorescence Detector, respectively. Fractions were collected and rotary-evaporated in a 45 °C water bath in the dark. Purified samples were stored at -20 °C prior to further NMR and MS analysis. For *B. puraquae* AFC purification, the whole procedure, including bacterial culture and extraction, was performed via the method previously described[3].

**UHPLC and UHPLC-HR-MS/MS analysis**

The analysis of crude and purified AFC was performed in a UltiMate™ 3000 UHPLC (ThermoScientific, USA) equipped with a reverse phase C18, 4.6 mm x 250 mm, 5.0 µm, 130 Å chromatography column (Waters Xselect CSH). The column was eluted with A: $H_2O$ + 0.05% formic acid and B : MeCN + 0.05% formic acid with a flow rate of 1 mL/min at 45°C and with the following gradient: 0-1 min, 4%-44% B; 1-9 min, 44% B; 9-10 min, 44%-100% B; 10-14 min, 100% B; 14-14.1 min, 100%-4% B; 14.1-16 min, 4% B. AFC elution was tracked by UV absorption at 320 nm and fluorescence ($\lambda_{ext}$ = 330 nm, $\lambda_{em}$ = 410 nm) by RS Diode Array Detector and RS Fluorescence Detector, respectively.

UHPLC-HR-MS/MS analysis was performed by timsTOF Pro hybrid quadrupole-time-of-flight (QTOF) mass spectrometer equipped with trapped ion mobility spectrometry (TIMS) produced by Bruker (Bremen, Germany) or Q Exactive hybrid quadrupole-Orbitrap mass spectrometer by Thermo Fisher Scientific (Waltham, MA, USA) equipped with a heated ESI source at position C and a voltage of ±3.5 kV. A Vanquish UHPLC system and was used with ACQUITY Premier

CSH C18 Column 1.7 µm, 2.1 x 100 mm reverse column (Waters). The column was eluted with A: $H_2O$ + 0.1% formic acid and B: MeCN + 0.1% formic acid with a flow rate of 0.3 mL/min at 28 °C and with the following gradient: 0-2 min, 4% B; 2-8 min, 4%-100% B; 8-12 min, 100% B; 12-12.1 min, 100%-4% B; 12.1-14 min, 4% B. MS data was acquired in positive ESI ionization mode. For timsTOF, the data was recorded without ion mobility and the scan range was set to 80 to 1200 *m/z* at a 12 Hz base acquisition rate. Fragment spectra were acquired using the data-dependent acquisition mode (AutoMS/MS) employing 30 and 70 eV fragmentation energies. Mass calibration was performed using the Agilent low concentration tuning mix (13 compounds in acetonitrile, part number G1969-85020) prior to analysis. For additional mass accuracy, a calibration segment was programmed from 0.05 to 0.15 min at every UHPLC run with the help of a 6-port-valve with a 20 µL loop which contained a solution of 10 mM sodium formate. The Q Exactive instrument was operated in high-resolution full-scan mode. For full MS, a resolution of 70,000 (200 *m/z* at full width at half maximum) was selected with a maximum IT of 200 ms and an AGC target of 3e6. Scan range was set at 80-800 *m/z*. For PRM, ion 452.2643 was selected, and a resolution of 35,000 (200 *m/z* at full width at half maximum) was applied with a maximum IT of 110 ms and an AGC target of 5e5. The instrument was calibrated at a mass accuracy ≤ 2 ppm with a PierceTM LTQ Velos ESI Positive Ion Calibration Solution (Thermo). External calibration was performed before each measurement run.

Data quantification was performed by TASQ (Bruker, v2022b). AFC ions with 1 and 2 positive charges were selected and all the automatic integrations were revised manually. The ratio of ion abundance was made from an average of 3 independent replicates. Note that since the major AFC-I and AFC-J ions have 2 positive charges, the ratio may not reflect the real quantity of these molecules. Molecular Networking analysis was further processed on Δ*afcF*, Δ*afcS* and wildtype strains with the software MZmine (version 4.7.8) using MZwizard[6–8]. Intensity in all the MS spectra are shown in either relative abundance (%) or arbitrary units (a.u.).

**NMR analysis**

$^{1}$H NMR spectra were recorded in DMSO-$d_6$ on the instruments Bruker AV-600 (600 MHz) or on the instrument Bruker AVANCE NEO (1.2 GHz); chemical shifts δ in ppm relative to solvent signals ($\delta$ = 2.50 ppm for DMSO-$d_6$), coupling constants $J$ are given in Hz. $^{13}$C NMR spectra were recorded in DMSO-$d_6$ on the instruments Bruker AV-600 (126 MHz); chemical shift in ppm relative to solvent signals ($\delta$ = 39.53 ppm for DMSO-$d_6$). The NMR spectra were recorded at 313K.

**Ozonolysis of AFC**

The procedure followed reported protocols with modifications[9–11]. AFC (100 µL, DMF) was transferred to a round bottom flask (5 mL), and the solution was diluted with MeOH (900 µL), cooled to -78 °C, and treated with ozone for 2 min. Nitrogen was bubbled into the blue solution for 1 min before $H_2O_2$ (50 µL, 30%) was added, and the solution was stirred at RT for 1 h. The solvent was evaporated, and the solids were dissolved in MeOH/$H_2O$ (100 µL, 20%), centrifuged (5 min at 20,238 × $g$) and analyzed by UHPLC-HR-MS with a UHPLC system (Vanquish Horizon, ThermoFischer) equipped with a Diode Array Detector, Split Sampler HT, Binary Pump H, Column Compartment H connected to an HRMS (Exploris 240, ThermoFischer) in negative and positive modes using a HILIC column (BEH amide from Waters, 2.1 x 100 mm, 1.7 µm).

For the analysis, 2 µL was injected, the flow rate was 400 µL/min, the column oven was set to 40 °C, and the solvent was composed of A: a 5 mM aq. $NH_4HCO_3$ soln. and B: MeCN. The column was equilibrated for 0.5 min with 95% B and, after injection, the isocratic gradient was kept for

0.5 min at 95% B. The running gradient was decreased to 5% B over 3 min and stayed at 5% B for 2 min. The gradient was increased to the initial conditions at 95% B over 0.1 min and kept for 1 min at those conditions. The ion source parameters of the MS in positive and negative mode were set as follows: spray voltage 2.5 kV for negative mode and 3.5 kV for positive mode; ion transfer tube temperature at 325 °C; sheath gas 50 L/min; aux gas 10 L/min; sweep gas 1 L/min; and vaporizer temperature 340 °C. The EASY-ICTM internal calibration system was used at the beginning of each run. A full scan with a resolution of 60,000, a scan range of 70-900 *m/z*, and RF lens of 70% were used in positive mode for one run and negative mode for another run and a $ddMS^2$ was set up with a scan width of 0.4 *m/z*, at a resolution of 30,000, and with an HCD collision energy at 15, 30, and 50%) and using a previously defined list of potential degradation products ($C_{10}H_{15}O_7^-$ $[M-H]^-$, $C_{11}H_{21}O_3N^-$ $[M-H]^-$, $C_2H_3O_3N^-$ $[M-H]^-$, $C_6H_7O_7N^-$ $[M-H]^-$, and $C_{11}H_{23}O_3N^+$ $[M+H]^+$).

**Supplementary tables**

**Table S11**. Summary of chemical shifts and correlations for AFC-A (**1**). The chemical shifts of the carbon atoms were determined by a combination of $^{1}$H-$^{13}$C-HMBC and $^{1}$H-$^{13}$C-HSQC-TOCSY analyses.

| Moiety | Pos. | $\delta_H$ in ppm ($J_{HH}$ in Hz) | $\delta_C$, type | $^{1}$H-$^{13}$C HMBC | $^{1}$H-$^{1}$H NOESY | $^{1}$H-$^{1}$H TOCSY (100ms) | $^{1}$H-$^{13}$C HSQC-TOCSY (60ms) | $^{1}$H-$^{13}$C HSQC-TOCSY (90ms) |
|---|---|---|---|---|---|---|---|---|
| **DHLys** | 1 | | nd | | | | | |
| | 2 | 4.30 (m) | 52.3, CH | | DHLys(3a,3b) | DHLys(3a,3b,4,5,NH) | DHLys(3) | |
| | 3a | 2.51 (m) | 29.5, $CH_2$ | | | DHLys(2,4,5,NH) | | |
| | 3b | 2.51 (m) | | | | DHLys(2,4,5,NH) | | |
| | 4 | 5.72 (m) | nd | DHLys(6) | DHLys(5) | DHLys(2,3a,3b,5,6a,6b) | DHLys(3,6) | |
| | 5 | 5.54 (dt, $J$ = 11.0,7.4) | 123.4, CH | DHLys(3) | | DHLys(2,3a,3b,4,6a,6b) | DHLys(6) | DHLys(6) |
| | 6a | 3.46 (dd, $J$ = 13.7,7.4) | 35.2, $CH_2$ | DHLys(4,5) | | DHLys(3a,3b,4,5) | DHLys(5) | |
| | 6b | 3.34 (m) | | | | DHLys(3a,3b,4,5) | | |
| | NH | 7.91 (br) | | | DBA$^{1}$(NH) | DHLys(2,3a,3b) | | |
| | $NH_2$ | | | | | | | |
| **DBA$^{1}$** | 1 | | 165.1, C | | | | | |
| | 2 | 5.77 (d, $J$ = 13.9) | 104.2, CH | DBA$^{1}$(1, 3) | DHLys(NH), DBA$^{1}$(NH), DBA$^{2}$(2) | DBA$^{1}$(3,NH) | DBA$^{1}$(3) | DBA$^{1}$(3) |
| | 3 | 7.72 (dd, $J$ = 13.9, 11.1) | 133.1, CH | DBA$^{1}$(1), DBA$^{2}$(1) | DBA$^{1}$(2,NH) | DBA$^{1}$(2,NH) | DBA$^{1}$(2) | DBA$^{1}$(2) |
| | NH | 10.56 (d, $J$ = 11.1) | | DBA$^{2}$(1) | DBA$^{1}$(2,3), DBA$^{2}$(2,3) | DBA$^{1}$(2,3) | DBA$^{1}$(2,3) | DBA$^{1}$(2,3) |
| **DBA$^{2}$** | 1 | | 165.7, C | | | | | |
| | 2 | 5.29 (m) | 98.0, CH | DBA$^{2}$(1) | DBA$^{1}$(2,NH), DBA$^{2}$(3) | DBA$^{2}$(3,NH) | | DBA$^{2}$(3) |
| | 3 | 7.46 (dd, $J$ = 11.2,8.9) | 136.0, CH | DBA$^{2}$(1) | DBA$^{1}$(NH), DBA$^{2}$(2) | DBA$^{2}$(2,NH) | | DBA$^{2}$(2) |
| | NH | 10.97 (d, $J$ = 11.2) | | DBA$^{3}$(1) | DBA$^{1}$(3), DBA$^{2}$(2,3), DBA$^{3}$(2) | DBA$^{2}$(2,3) | DBA$^{2}$(2) | DBA$^{2}$(2,3) |
| **DBA$^{3}$** | 1 | | 165.3, C | | | | | |
| | 2 | 5.49, (d, $J$ = 8.8) | 97.0, CH | DBA$^{3}$(1) | DBA$^{2}$(NH), DBA$^{3}$(3) | DBA$^{3}$(3,NH) | | DBA$^{3}$(3) |
| | 3 | 7.44 (dd, $J$ = 11.3,8.8) | 137.6, CH | DBA$^{3}$(1) | DBA$^{2}$(NH), DBA$^{3}$(2), FA(2) | DBA$^{3}$(2,NH) | | DBA$^{3}$(2) |
| | NH | 10.89 (d, $J$ = 11.3) | | | DBA$^{3}$(3), FA(2) | DBA$^{3}$(2,3) | DBA$^{3}$(2) | DBA$^{3}$(3) |
| **FA** | 1 | | 170.9, C | | | | | |
| | 2 | 2.41 (t, $J$ = 7.4) | 35.2, $CH_2$ | FA (1,3,4) | DBA$^{3}$(NH,3), FA(3,4) | FA (3,4) | FA (3,4) | FA (3,4) |
| | 3 | 1.55 (m) | 24.0, $CH_2$ | FA (1,4) | DBA$^{3}$(NH), FA(2,4) | FA (2) | FA (2,4) | FA (2,4) |
| | 4 | 1.27 (m) | 28.0, $CH_2$ | | | | | |
| | 5 | nd | nd | | | | | |
| | 6 | nd | nd | | | | | |
| | 7 | nd | nd | | | | | |
| | 8 | nd | 28.3, $CH_2$ | | | | | |
| | 9 | nd | 28.7, $CH_2$ | | | | | |
| | 10 | 1.97 (q, $J$ = 6.9) | 26.3, CH | FA(8,9,11/12) | FA(11/12) | FA (11/12) | FA (8,9,11/12) | FA (8,9,11/12) |
| | 11 | 5.31 (dt, $J$ = 11.8, 6.9) | 129.2, $CH_2$ | FA(10,13) | FA(10,13) | FA (10,13,16) | FA (9,10,13,14,15) | FA (8,9,10,13,14,15,16) |
| | 12 | 5.31 (dt, $J$ = 11.8, 6.7) | 129.2, $CH_2$ | FA(10,13) | FA(10,13) | FA (10,13,16) | FA (9,10,13,14,15) | FA (8,9,10,13,14,15,16) |
| | 13 | 1.95 (q, $J$ = 6.7) | 26.4, CH | FA(11/12,14,15) | FA(11/12,16) | FA (11/12,16) | FA (11/12,14,15,16) | FA (11/12,14,15,16) |
| | 14 | 1.22(m) | 29.3, $CH_2$ | | | | | |
| | 15 | 1.31(m) | 22.3, $CH_2$ | | | | | |
| | | 1.22(m) | | | | | | |
| | 16 | 1.57 (m) | 35.6, $CH_2$ | FA (15,17) | FA (13,19a,19b,21) | FA (11/12,13) | FA (13,14,15) | FA (13,14,15) |
| | 17 | | 79.1, C | | | | | |
| | 18/20 | | 174.0/172.1, C | | | | | |
| | 19a | 2.75 (d, $J$ = 14.5) | 42.8, $CH_2$ | FA (17,18/20,21) | FA (15,16,19b,21) | FA (19b) | | |
| | 19b | 2.23 (d, $J$ = 14.5) | | | FA (15,16,19a,21) | FA (19a) | | |
| | 21 | 3.08 (s) | 50.6, $CH_3$ | FA (17) | FA (15,16,19a,19b) | | | |

Data highlighted in red were determined from measurements performed on a 1.2 GHz spectrometer.

**Table S2**. Summary of chemical shifts and correlations for AFC (**1**) obtained from LB medium. The chemical shifts of the carbon atoms were determined by a combination of $^{1}$H-$^{13}$C-HMBC and $^{1}$H-$^{13}$C-HSQC-TOCSY analyses.

| Moiety | Pos. | $\delta_H$ in ppm ($J_{HH}$ in Hz) | $\delta_C$, type | $^{1}$H-$^{13}$C HMBC | $^{1}$H-$^{1}$H NOESY | $^{1}$H-$^{1}$H TOCSY | $^{1}$H-$^{13}$C HSQC-TOCSY |
|---|---|---|---|---|---|---|---|
| DHLys | 1 | | 172.6, C | | | | |
| | 2 | 4.39 (td, $J$ =8.2, 5.5) | 51.0, CH | DHLys(1,3,4), DBA$^1$(1) | DHLys(3a,3b,NH) | DHLys(3a,3b,4,5,NH) | DHLys(3) |
| | 3a | 2.46 (m) | 29.0, $CH_2$ | DHLys(1,2,4,5) | | DHLys(2,4,5,6,NH) | DHLys(2,4) |
| | 3b | 2.53 (m) | | DHLys(1,2,4,5) | | DHLys(2,4,5,6,NH) | DHLys(2,4) |
| | 4 | 5.68 (m) | 130.7, CH | DHLys(6) | DHLys(5) | DHLys(2,3a,3b,5,6,NH) | DHLys(3,5,6) |
| | 5 | 5.52 (m) | 123.7, CH | DHLys(3) | DHLys(4,6) | DHLys(2,3a,3b,4,6,NH) | DHLys(4,6) |
| | 6 | 3.47 (m) | 35.4, $CH_2$ | DHLys(4,5) | | DHLys(3a,3b,4,5) | DHLys(4,5) |
| | NH | 8.12 (d, $J$ =8.2) | | DBA$^1$(1) | DHLys(2,3a,3b,); DBA$^1$(2,NH) | DHLys(2,3a,3b,4,5,6) | DHLys(2) |
| | $NH_2$ | | | | | | |
| DBA$^1$ | 1 | | 165.4, C | | | | |
| | 2 | 5.75 (d, $J$ =14.0) | 103.6, CH | DBA$^1$(1, 3) | DHLys(NH), DBA$^1$(3,NH), DBA$^2$(2) | DBA$^1$(3,NH) | DBA$^1$(3) |
| | 3 | 7.74 (dd, $J$ =14.0, 11.1) | 133.3, CH | DBA$^1$(1) | DHLys(NH), DBA$^1$(2,NH) | DBA$^1$(2,NH) | DBA$^1$(2) |
| | NH | 10.59 (d, $J$ =11.1) | | DBA$^2$(1) | DHLys(NH), DBA$^1$(2,3), DBA$^2$(2,3) | DBA$^1$(2,3) | DBA$^1$(2,3) |
| DBA$^2$ | 1 | | 165.6, C | | | | |
| | 2 | 5.29 (m) | 97.8, CH | DBA$^2$(1,3) | DBA$^1$(2,NH), DBA$^2$(3) | DBA$^2$(3,NH) | DBA$^2$(3) |
| | 3 | 7.47 (dd, $J$ =11.2, 8.9) | 136.0, CH | DBA$^2$(1,2) | DBA$^1$(NH), DBA$^2$(2,NH) | DBA$^2$(2,NH) | DBA$^2$(2) |
| | NH | 10.97 (d, $J$ =11.2) | | DBA$^3$(1) | DBA$^2$(3), DBA$^3$(2) | DBA$^2$(2,3) | DBA$^2$(3) |
| DBA$^3$ | 1 | | 165.3, C | | | | |
| | 2 | 5.49 (d, $J$ =8.8) | 96.8, CH | DBA$^3$(1) | DBA$^2$(NH), DBA$^3$(3) | DBA$^3$(3,NH) | DBA$^3$(3) |
| | 3 | 7.44 (dd, $J$ =11.0, 8.8) | 137.7, CH | DBA$^3$(1) | DBA$^3$(2,NH), FA(2) | DBA$^3$(2,NH) | DBA$^3$(2) |
| | NH | 10.87 (d, $J$ =11.0) | | | DBA$^3$(3), FA(2) | DBA$^3$(2,3) | DBA$^3$(3) |
| FA | 1 | | 170.8, C | | | | |
| | 2 | 2.41 (t, $J$ =8.4) | 35.1, $CH_2$ | FA(1,3,4) | DBA$^3$(NH,3), FA(3) | FA(3,4) | FA(3,4) |
| | 3 | 1.56 (m) | 23.9, $CH_2$ | FA(1,2,4,5) | DBA$^3$(NH), FA(2,4) | FA(2,4) | FA(2,4) |
| | 4 | 1.27 (m) | 27.9, $CH_2$ | | | | |
| | 5 | | | | | | |
| | 6 | | | | | | |
| | 7 | | | | | | |
| | 8 | | | | | | |
| | 9 | 1.26 (m) | 28.3, $CH_2$ | | | | |
| | 10/13 | 1.98 (m) | 26.1, $CH_2$ | FA(9,11/12,14) | FA(9,11/12,14,16) | FA(9,11/12,14,16) | FA(9,11/12,14,15) |
| | 11//12 | 5.32 (m) | 129.1, CH | FA(10/13) | FA(10/13) | FA(9,10/13,14,16) | FA(9,10/13,14) |
| | 14 | 1.26 (m) | 29.1, $CH_2$ | | | | |
| | 15 | 1.22 (m) | 22.1, $CH_2$ | | | | |
| | 16 | 1.69 (m) | 33.8, $CH_2$ | | FA(21) | FA(10/13, 11/12) | FA(14,15) |
| | 17 | - | 79.7, C | | | | |
| | 18 | | | | | | |
| | 19 | 2.67 (m) | 39.3, $CH_2$ | FA(17) | FA(21) | | |
| | 20 | | | | | | |
| | 21 | 3.15 (s) | 50.5, $CH_3$ | FA(17) | FA(15,16) | | |

**Table S3.** Expected structures of AFCs after supplementation with unlabeled or deuterium-labeled myristic acid, palmitic acid, or *cis*-vaccenic acid, along with the corresponding $[M+H]^+$ ion structure and molecular formulas. Possible deuterium labeled positions are highlighted in green.

| | | | |
|---|---|---|---|
| **Myristic acids** | 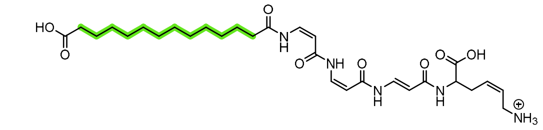 | $C_{29}H_{46}N_5O_8^+$ **(11)** | $C_{29}H_{22}D_{24}N_5O_8^+$ **(12)** |
| | 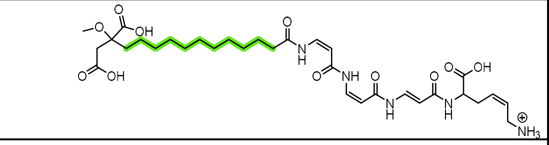 | $C_{32}H_{50}N_5O_{11}^+$ **(13)** | $C_{32}H_{28}D_{22}N_5O_{11}^+$ **(14)** |
| | 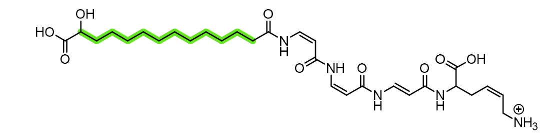 | $C_{29}H_{46}N_5O_9^+$ **(15)** | ($C_{29}H_{23}D_{23}N_5O_9^+$ **16)** |
| **Palmitic acids** | 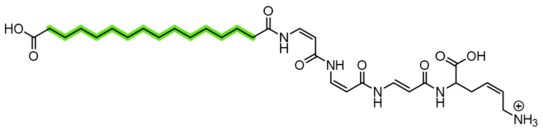 | $C_{31}H_{50}N_5O_8^+$ **AFC-D (4)** | $C_{31}H_{22}D_{28}N_5O_8^+$ **AFC-D-*d*$_{28}$ (18)** |
| | 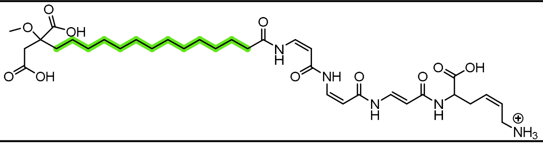 | $C_{34}H_{54}N_5O_{11}^+$ **AFC-B (2)** | $C_{34}H_{28}D_{26}N_5O_{11}^+$ **AFC-B-*d*$_{26}$ (17)** |
| ***cis*-vaccenic acids** | 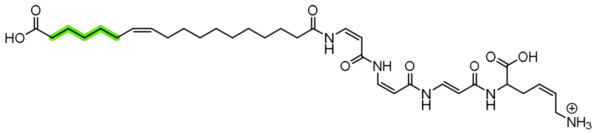 | $C_{33}H_{52}N_5O_8^+$ **AFC-C (3)** | $C_{33}H_{42}D_{10}N_5O_8^+$ **AFC-C-*d*$_{10}$ (20)** |
| | 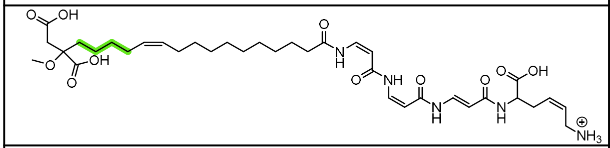 | $C_{36}H_{56}N_5O_{11}^+$ **AFC-A (1)** | $C_{36}H_{48}D_8N_5O_{11}^+$ **AFC-A-*d*$_8$ (19)** |
| | 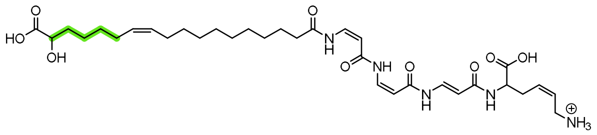 | $C_{33}H_{52}N_5O_9^+$ **AFC-E (5)** | $C_{33}H_{43}D_9N_5O_9^+$ **AFC-E-*d*$_9$ (22)** |
| | 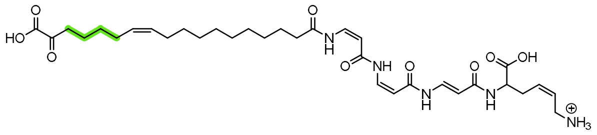 | $C_{33}H_{50}N_5O_9^+$ **AFC-G (7)** | $C_{33}H_{42}D_8N_5O_9^+$ **AFC-G-*d*$_8$ (23)** |
| | 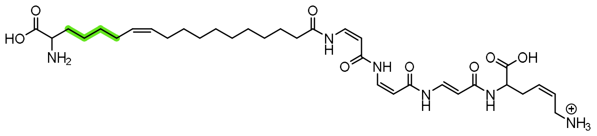 | $C_{33}H_{53}N_6O_8^+$ **AFC-I (9)** | $C_{33}H_{45}D_8N_6O_8^+$ **AFC-I-*d*$_8$ (24)** |

**Table S4.** Colony morphology and antifungal activity against *R. solani* of the *afc* mutants used in this study.

| Strain | Colony morphology | Antifungal Activity | | Reference |
|---|---|---|---|---|
| **AFC-producing** | | | | |
| WT | Rough | Yes | 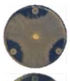 | Subramoni *et al*., 2013 |
| Δ*afcB* | Rough | Yes | 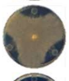 | This study |
| Δ*afcG* | Rough | Yes | 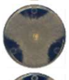 | This study |
| Δ*afcH* | Rough | Yes | 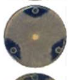 | This study |
| Δ*afcI* | Rough | Yes | 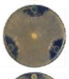 | This study |
| Δ*afcS* | Rough | Reduced | 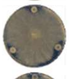 | This study |
| Δ*afcF* | Rough | Reduced | 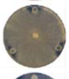 | This study |
| Δ*afcU* | Rough | No | 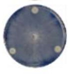 | This study |
| **AFC-non-producing** | | | | |
| Δ*afc* | Shiny | No | 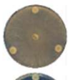 | This study |
| Δ*afcA* | Shiny | No | 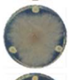 | This study |
| Δ*afcJ* | Shiny | No | 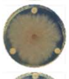 | This study |
| Δ*afcK* | Shiny | No | 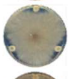 | This study |
| Δ*afcL* | Shiny | No | 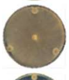 | This study |
| Δ*afcM* | Shiny | No | 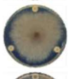 | This study |
| Δ*afcO* | Shiny | No | 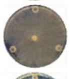 | This study |
| Δ*afcP* | Shiny | No | 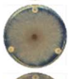 | This study |
| Δ*afcQ* | Shiny | No | 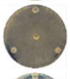 | This study |
| Δ*afcR* | Shiny | No | 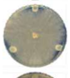 | This study |
| Δ*afcV* | Shiny | No | 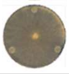 | This study |

**Table S5.** Bacterial strains

| Strain | Genotype or Description | Reference |
|---|---|---|
| ***E. coli* strains** | | |
| SY327λpir | wild type; *araD*, Δ(*lac pro*) *argE*(Am) *recA56* $Rif^R$ *nalA λpir* | 12 |
| DH5α | F- Φ80d*lacZΔM15* Δ(*lacZYA-argF*)U169 *recA1 endA1 hsdR17(rK- mK+) supE44 thi-1 relA1 gyrA96* | 13 |
| MC1061 | $F^-$ *araD139 Δ(ara, leu)7697 Δ(lacIPOZYA)X74 galK16 galE15 rpsL150 mcrA0 mcrB1 hsdR2 relA::IS2 spoT* $e14^-$ $\lambda^-$ ($Sm^R$) | 14 |
| ***Burkholderia* strains** | | |
| *Burkholderia lata* 383 | Isolated from forest soil in Trinidad and Tobago | 15 |
| *Burkholderia vietnamiensis* LMG10929 | Isolated from rice rhizosphere soil | 16 |
| *Burkholderia vietnamiensis* ABIP434 | Rice rhizosphere soil isolate | 17 |
| *Burkholderia thailandensis* E264 | isolated from soil of a rice field | 18 |
| *Burkholderia puraquae* DSM103137 | isolated from haemodialysis water | 19 |
| *Burkholderia cenocepacia* H111 | CF isolate from Germany, R-6282 | 20 |
| ***Burkholderia cenocepacia* K56-2 strains** | | |
| K56-2 | CF sputum isolate (Canada); ET12 lineage | 21 |
| Δ*afc* | K56-2 deletion mutant of the *afc* cluster | This study |
| Δ*afcA* | K56-2 deletion mutant of *afcA* | This study |
| Δ*afcB* | K56-2 deletion mutant of *afcB* | This study |
| Δ*afcG* | K56-2 deletion mutant of *afcG* | This study |
| Δ*afcH* | K56-2 deletion mutant of *afcH* | This study |
| Δ*afcI* | K56-2 deletion mutant of *afcI* | This study |
| Δ*afcJ* | K56-2 deletion mutant of *afcJ* | This study |
| Δ*afcK* | K56-2 deletion mutant of *afcK* | This study |
| Δ*afcL* | K56-2 deletion mutant of *afcL* | This study |
| Δ*afcM* | K56-2 deletion mutant of *afcM* | This study |
| Δ*afcO* | K56-2 deletion mutant of *afcO* | This study |
| Δ*afcP* | K56-2 deletion mutant of *afcP* | This study |
| Δ*afcQ* | K56-2 deletion mutant of *afcQ* | This study |
| Δ*rep* | K56-2 deletion mutant of the duplicated *afcREST* genes | This study |
| Δ*rep*Δ*afcR* | K56-2 Δ*rep* deletion mutant of *afcR* | This study |
| Δ*afcS* | K56-2 Δ*rep* derivative with *afcS* deleted | This study |
| Δ*afcU* | K56-2 deletion mutant of *afcU* | This study |
| Δ*afcV* | K56-2 deletion mutant of *afcV* | This study |
| Δ*afcF* | K56-2 Δ*rep*Δ*afcR* derivative with *afcF* deleted bearing pBBR-afcR | This study |
| **Other strains** | | |
| *Bacillus subtilis* 168 | With plasmid pHY300-zsGreen-term to confer resistance to Tc | 22 |

**Table S6.** Plasmids

| **Plasmids** | **Description** | **Reference** |
|---|---|---|
| pRK2013 | Helper plasmid; RK2 derivative, mob+ tra+ ori ColE1; $Km^R$ | [23] |
| pGPI-SceI | Suicide plasmid with oriR6K, mob+, I-SceI restriction site; $Tp^R$ | Elisabeth Steiner, Laboratory collection |
| pDAI-SceI | pDA17 plasmid carrying the I-SceI nuclease gene; $Tc^R$ | |
| pBBR1MCS-3 | Mobilisable BHR cloning vector; IncP and ColE1 compatible; $Tc^R$) | [24] |
| pBBR-afcR | pBBR1MCS-3 containing the *afcR* gene | This study |

**Table S7.** Primers used in this study.

| Name | Sequence (5´→3´)[a] | Construction |
|---|---|---|
| upXhoF | gcgcctcgagTTCTGATCTTCCTCGTGCTC | Δ*afc* |
| upBglIIR | gcgcagatctGCAACGCAATCAGAACACC | Δ*afc* |
| afcKOdownPstF | gcgcctgcagATTCATCTTGACGGTCGTCG | Δ*afc* |
| afcKOdownBamR | gcgcggatccTTGATCGTACTGGCTGAAGT | Δ*afc* |
| afcA up XbaI | cgcgTCTAGACTGGTACGCATGAAAC | Δ*afcA*, XbaI site |
| afcA SOE R | TCATGCGAACGCTCCCTGAGGATTCTCCCCTTG | Δ*afcA* |
| afcA SOE F | CAAGGGGAGAATCCTCAGGGAGCGTTCGCATGA | Δ*afcA* |
| afcA down EcoRI | gcgcGAATTCAATCAGAAAGGCGGTGC | Δ*afcA*, EcoRI site |
| afcB_up_XbaI_F | gcgcTCTAGACTACCTGAACCCCGACGA | Δ*afcB*, XbaI site |
| afcB_SOE_R | AACGTCGACCAGTTCATCATGCGAACGCTCCC | Δ*afcB* |
| afcB_SOE_F | GGGAGCGTTCGCATGATGAACTGGTCGACGTT | Δ*afcB* |
| afcB_down_SmaI_R | gcgcCCCGGGACCGTGTGATAGTCGAAGAA | Δ*afcB*, SmaI site |
| 220_up_XbaI_F | gcgcTCTAGATGTTCCAGCAATTCAACCTG | Δ*afcG*, XbaI site |
| 220_SOE_R | GACGAATGCGATTCATTACGCCACGCAAACGT | Δ*afcG* |
| 220_SOE_F | ACGTTTGCGTGGCGTAATGAATCGCATTCGTC | Δ*afcG* |
| 220_down_SmaI_R | gcgcCCCGGGAGCTTGATCGTCGGGTAGA | Δ*afcG*, SmaI site |
| 219_up_XbaI_F | gcgcTCTAGAGTATCGCTCTATCACCAGGT | Δ*afcH*, XbaI site |
| 219_SOE_R | CGTACTCACTGGCCATCATGGGGCGAGTCCTT | Δ*afcH* |
| 219_SOE_F | AAGGACTCGCCCCATGATGGCCAGTGAGTACG | Δ*afcH* |
| 219_down_SmaI_R | gcgcCCCGGGCCCATTTCGCTTCGACGA | Δ*afcH*, SmaI site |
| 218_up_XbaI_F | gcgcTCTAGAGAGAACGACGGCCAGAAAT | Δ*afcI*, XbaI site |
| 218_SOE_R | CATGGCAGGGTTCCTGTCACTGGCCATCGGCG | Δ*afcI* |
| 218_SOE_F | CGCCGATGGCCAGTGACAGGAACCCTGCCATG | Δ*afcI* |
| 218_down_SmaI_R | gcgcCCCGGGGCCGTCGCTCATCAGGAA | Δ*afcI,* SmaI site |
| 217 up XbaI | gcgcTCTAGACCGCAACGATCGCCAATATA | Δ*afcJ*, XbaI site |
| 217 SOE R | CGGCCTGTCGCGTGGAGGCAGGGTTCCTGTCA | Δ*afcJ* |
| 217 SOE F | TGACAGGAACCCTGCCTCCACGCGACAGGCCG | Δ*afcJ* |
| 217 down EcoRI | gcgcGAATTCATGTTCAGCGCTTCCGGAT | Δ*afcJ*, EcoRI site |
| 216 up XbaI | gcgcTCTAGACCCACCGTGTTCAACAAATA | Δ*afcK*, XbaI site |
| 216 SOE R | CGCGAGCATGGTCGGAGAGGGGAGCTCCTGAA | Δ*afcK* |
| 216 SOE F | TTCAGGAGCTCCCCTCTCCGACCATGCTCGCG | Δ*afcK* |
| 216 down EcoRI | gcgcGAATTCCGAATCGTTCATCAGCAGCA | Δ*afcK*, EcoRI site |
| 215_up_XbaI_F | gcgcTCTAGAAAGCCGGCAGTTGATCC | Δ*afcL*, XbaI site |
| 215_SOE_R | AACGGGTCGCACCATGGGTCGGATCAGGCCGC | Δ*afcL* |
| 215_SOE_F | GCGGCCTGATCCGACCCATGGTGCGACCCGTT | Δ*afcL* |
| 215_down_EcoRI_R | GCAGCACACGCGGTTAT | Δ*afcL* |
| 214 up XbaI | tataTCTAGAGCGACATCCGCACGTTCT | Δ*afcM*, XbaI site |
| 214 SOE F | GACGGCGAGGCGTGACCATGAATGACACGCTT | Δ*afcM* |
| 214 SOE R | AAGCGTGTCATTCATGGTCACGCCTCGCCGTC | Δ*afcM* |

| | | |
|---|---|---|
| 214 down EcoRI | gcgcGAATTCTTCGACGCACAGATCA | Δ*afcM*, EcoRI site |
| 212_up_XbaI_F | gcgcTCTAGAACCTGTTCCTGCGAATCC | Δ*afcO*, XbaI site |
| 212_SOE_R | GGGCGCTCAGGCGTCATCAGTTGAGCGGCATC | Δ*afcO* |
| 212_SOE_F | GATGCCGCTCAACTGATGACGCCTGAGCGCCC | Δ*afcO* |
| 212_down_EcoRI_R | gcgcGAATTCGTCATCCCGAAGTGGTCGT | Δ*afcO*, EcoRI site |
| 211 up XbaI | gcgcTCTAGAATCATGATGGAGTGGCCGT | Δ*afcP*, XbaI site |
| 211 SOE R | GATCGGCTCCTCGGGTTCAGGCGTCATGGCGG | Δ*afcP* |
| 211 SOE F | CCGCCATGACGCCTGAACCCGAGGAGCCGATC | Δ*afcP* |
| 211 down EcoRI | gcgcGAATTCGTGGATTGACCGGCACGAA | Δ*afcP*, EcoRI site |
| 210_up_XbaI_F | gcgcTCTAGATTCGAACGCTATGCCGCC | Δ*afcQ*, XbaI site |
| 210_SOE_R | AGCGTCGACAGGGTCATCAGGCCGGCCGCACG | Δ*afcQ* |
| 210_SOE_F | CGTGCGGCCGGCCTGATGACCCTGTCGACGCT | Δ*afcQ* |
| 210_down_EcoRI_R | gcgcGAATTCCCGATCCCTTCGTTGAGC | Δ*afcQ*, EcoRI site |
| rep_up_XbaI_F | gcgcTCTAGAGCGATGCGTTGCCCACCTACAT | Δ*rep*, XbaI site |
| rep_SOE_R | GTCGGGCGTCAGCGCGCCCATGCCGCGCGGCC | Δ*rep* |
| rep_SOE_F | GGCCGCGCGGCATGGGCGCGCTGACGCCCGAC | Δ*rep* |
| rep_down_EcoRI_R | gcgcGAATTCAACTGGCGGAAGAACGGGT | Δ*rep*, EcoRI site |
| 209 up XbaI | gcgcTCTAGATCGAGGACGTGCTGTT | Δ*afcR*, XbaI site |
| 209 SOE R | ACCTTGTATGCGCTCATCATGCGGCGGTCTCC | Δ*afcR* |
| 209 SOE F | GGAGACCGCCGCATGATGAGCGCATACAAGGT | Δ*afcR* |
| 209 down EcoRI | gcgcGAATTCGCCCATGAACATCTCG | Δ*afcR*, EcoRI site |
| 207_up_XbaI_F | gcgcTCTAGACCAGTACTACCGCGACGAA | Δ*afcS*, XbaI site |
| 207_SOE_R | CGAAAAGGGTGAGGGTGTCATTCCACCGCGGC | Δ*afcS* |
| 207_SOE_F | GCCGCGGTGGAATGACACCCTCACCCTTTTCG | Δ*afcS* |
| 207_down_EcoRI_R | cgcgGAATTCGGTACATCTCGGTCAGGC | Δ*afcS*, EcoRI site |
| 205 up XbaI | gcgcTCTAGAGAGATGTACCGCGTGCTGG | Δ*afcU*, XbaI site |
| 205 SOE R | CTCATCGTCGGCATCCCGCTCAGGCCAGTGCC | Δ*afcU* |
| 205 SOE F | GGCACTGGCCTGAGCGGGATGCCGACGATGAG | Δ*afcU* |
| 205 down EcoRI | gcgcGAATTCACTCGACCTTCTGCAATT | Δ*afcU*, EcoRI site |
| 204_up_XbaI_F | gcgcTCTAGAGACGCGAACCCGAACCTG | Δ*afcV*, XbaI site |
| 204_SOE_R | GGAACACCTTAGTCATCAGTCGAACAGCATGA | Δ*afcV* |
| 204_SOE_F | TCATGCTGTTCGACTGATGACTAAGGTGTTCC | Δ*afcV* |
| 204_down_SmaI_R | gcgcCCCGGGAGAATGCCGGTCGCGAAGAA | Δ*afcV*, SmaI site |
| 201 up XbaI | gcgcTCTAGACGGCGATCGTGTTCCTGTT | Δ*afcF*, XbaI site |
| 201 SOE R | GCCGTGAAGCGGCGCACGTCAGCTTGCCCGCG | Δ*afcF* |
| 201 SOE F | CGCGGGCAAGCTGACGTGCGCCGCTTCACGGC | Δ*afcF* |
| 201 down EcoRI | gcgcGAATTCTCAATCCTGCCGTCCATCCG | Δ*afcF*, EcoRI site |
| 209 C up XhoI | atatCTCGAGATGACCCTGTCGACGC | pBBR-*afcR*, XhoI site |
| 209 C down XbaI | cgcgTCTAGATCATGCGTCTCCTTCG | pBBR-*afcR*, XbaI site |

a: SOE primers designed according to Horton *et al.*[25] Not underlined nucleotides at the 5' end are the overhangs.

**Supplementary figures**

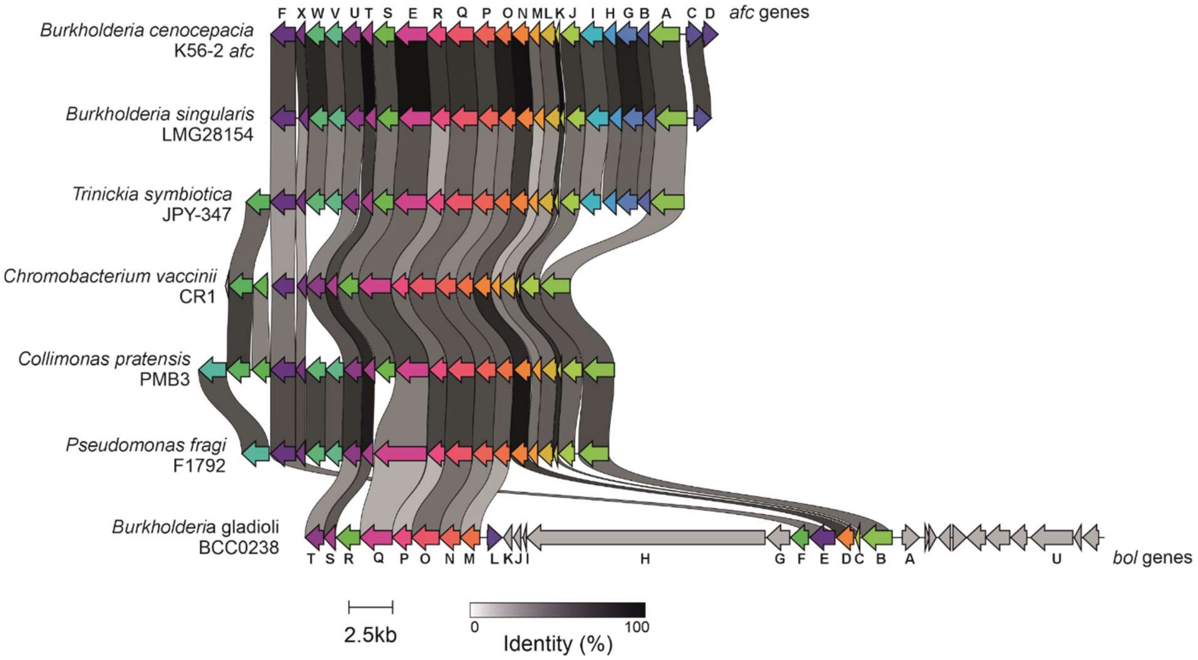


**Figure S1.** Alignment of the *B. cenocepacia* K56-2 *afc* gene cluster (top) and homologous gene clusters from diverse bacteria, including the *bol* cluster from *B. gladioli* BCC0238 (bottom). Gene designations for *afc* and *bol* are shown according to Zhong and Dashti, respectively[3,26]. Image generated using Clinker[27].

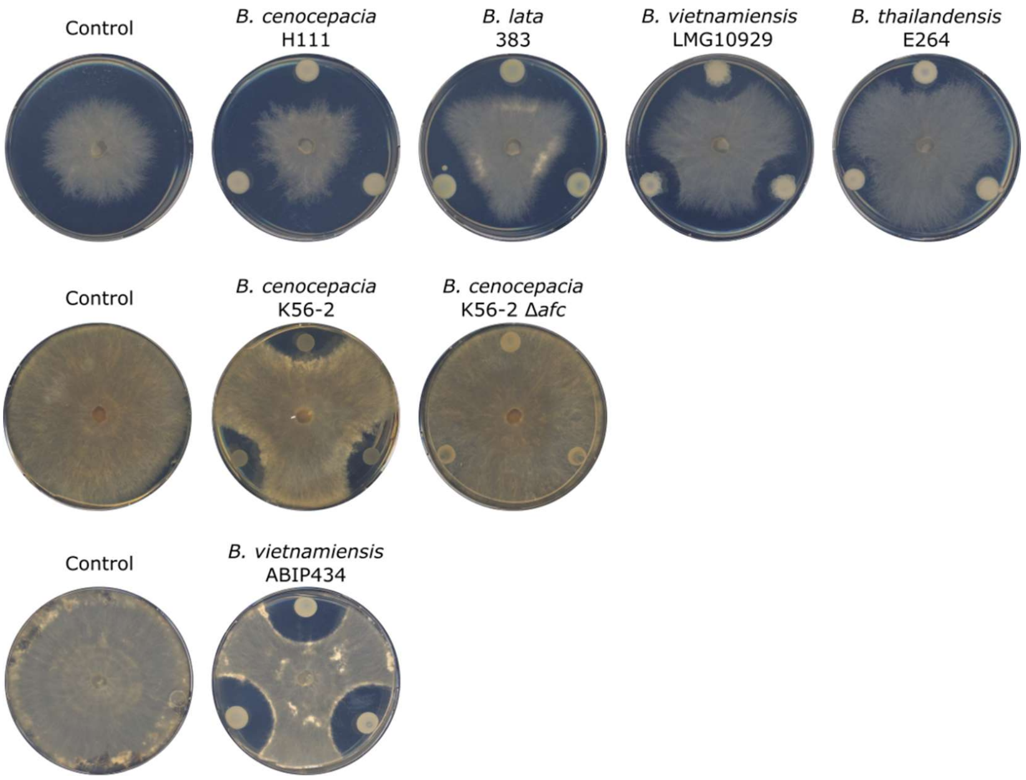


**Figure S2.** Antifungal activity of selected *Burkholderia* strains against *R. solani*.

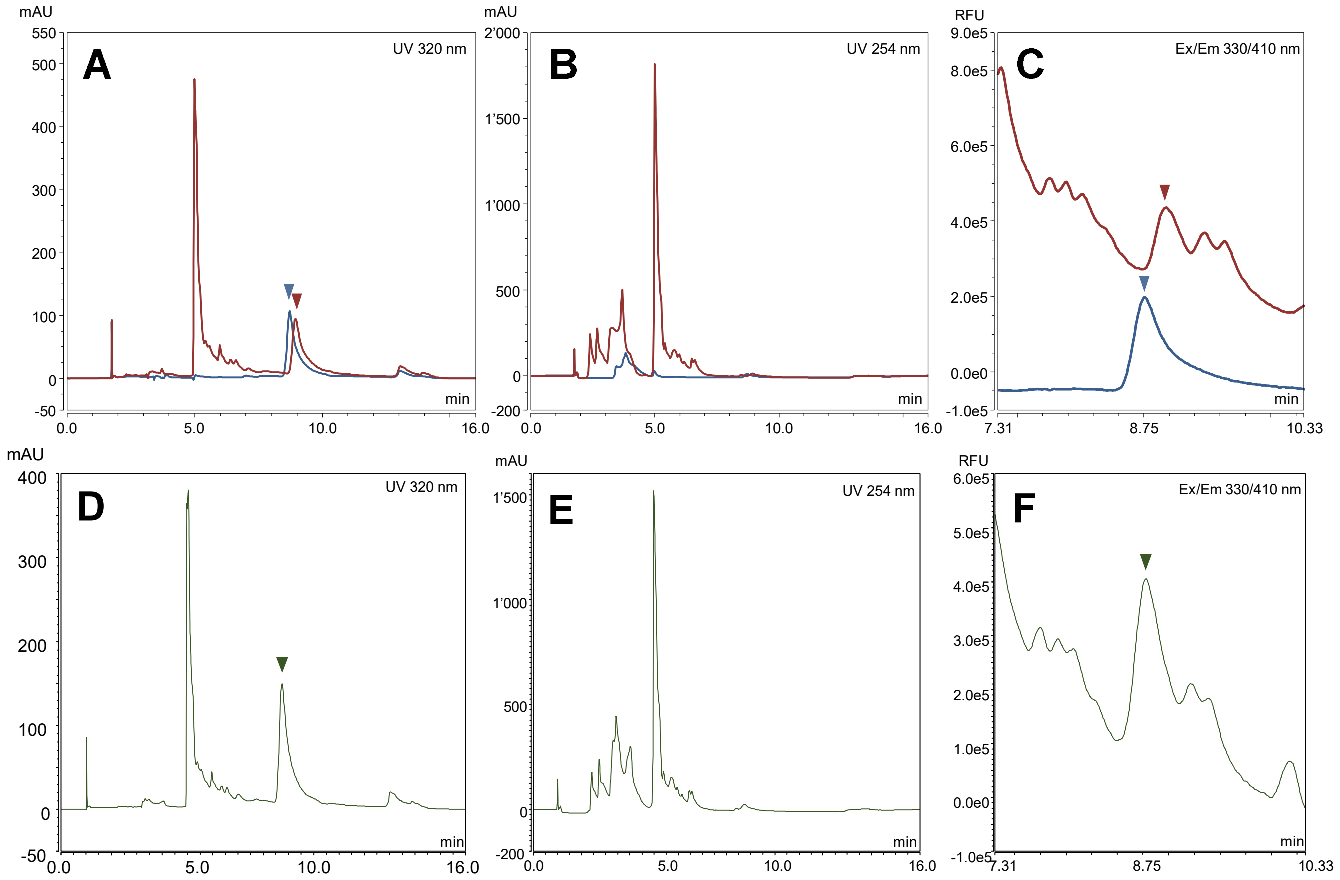


**Figure S3.** Comparison of the extraction methods described in this study (blue), in Zhong *et al.*[3] (brown) and a 1:1 mixture of the two (green) by analytical HPLC. (**A-B**) UV absorption at $\lambda$ = 320 nm (**C-D**) UV absorption at $\lambda$ = 254 nm. (**E-F**) Enlarged emission peak at $\lambda_{em}$ = 410 nm with $\lambda_{ex}$ = 330 nm at the time of elution of AFC. Peaks of AFC-A (**1**) are indicated with arrows.

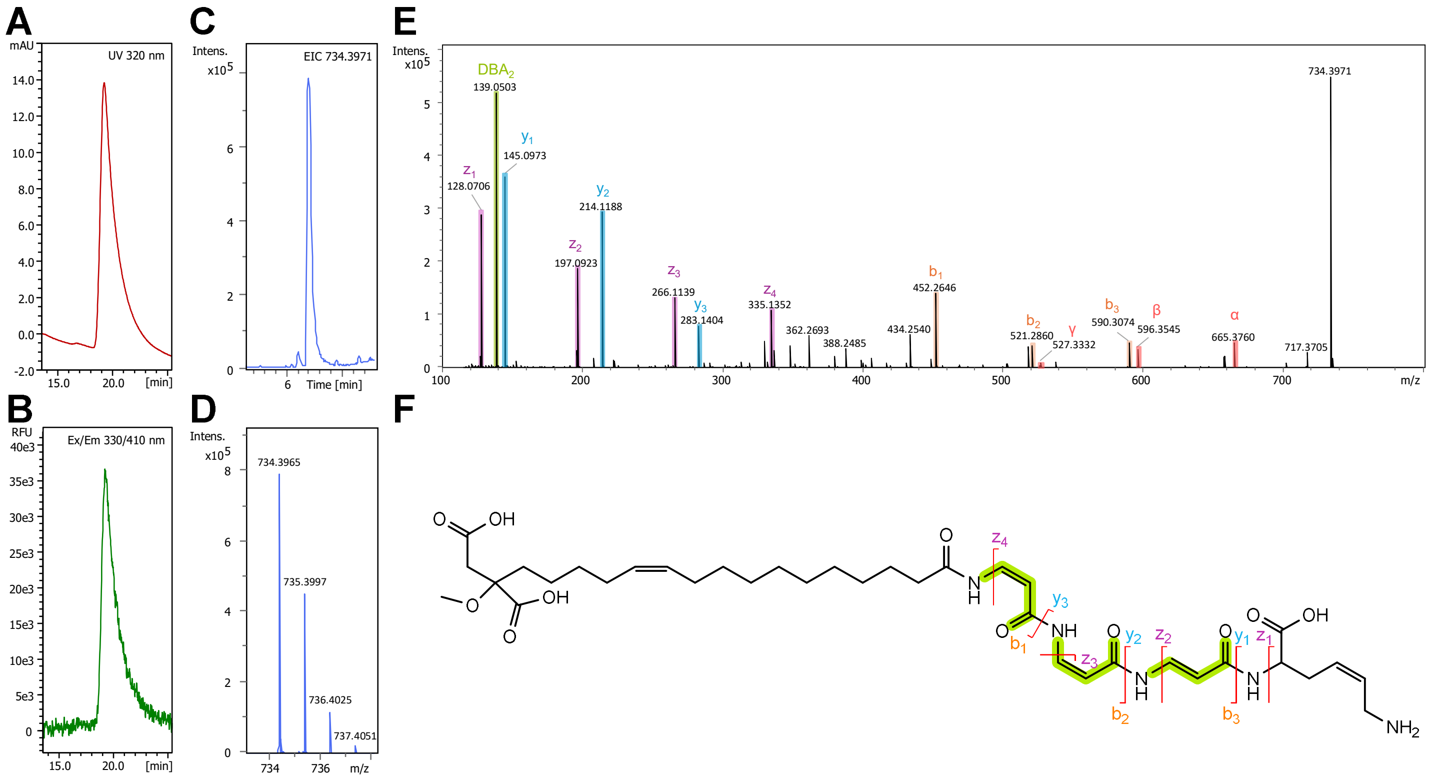


**Figure S4.** Identification of purified AFC-A (**1**) by UHPLC, UHPLC-HR-MS/MS and proposed assignment of selected fragments. Liquid chromatogram of **(A)** UV absorption at 320 nm and **(B)** fluorescence with $\lambda_{ex}$ / $\lambda_{em}$ = 330 nm / 410 nm **(C)** Extracted ion chromatograph of AFC-A (**1**). **(D)** Isotopic pattern of AFC-A $[M+H]^+$ ion. **(E)** HR-MS/MS fragmentation of AFC-A (**1**). Possible fragments involving the loss of one, two or three DBA units are labeled with α, β or γ in red, respectively. **(F)** Assignment of the fragments. DBA subunits are highlighted in light green.

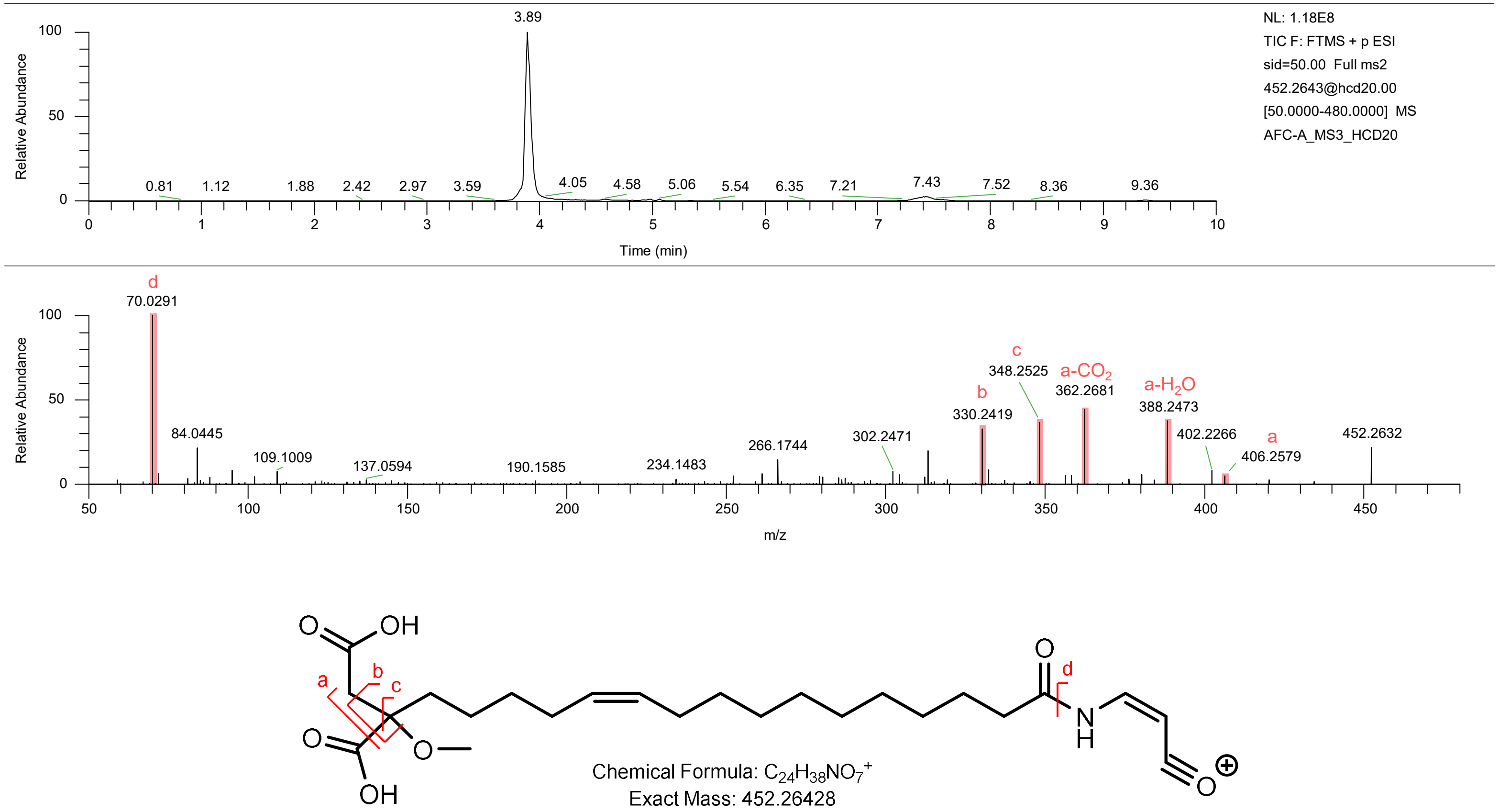


**Figure S5.** Pseudo-MS$^3$ identification of the fatty acid tail fragment of AFC-A (**1**). The identification of the functional group on the fatty acid tail was carried out by Q Exactive UHPLC-HR-MS/MS in positive mode. In-source fragmentation was applied, followed by fragmentation of the tail ion (*m/z* = 452.2643) with HCD at 20 eV. The identified fragments are shown in red.

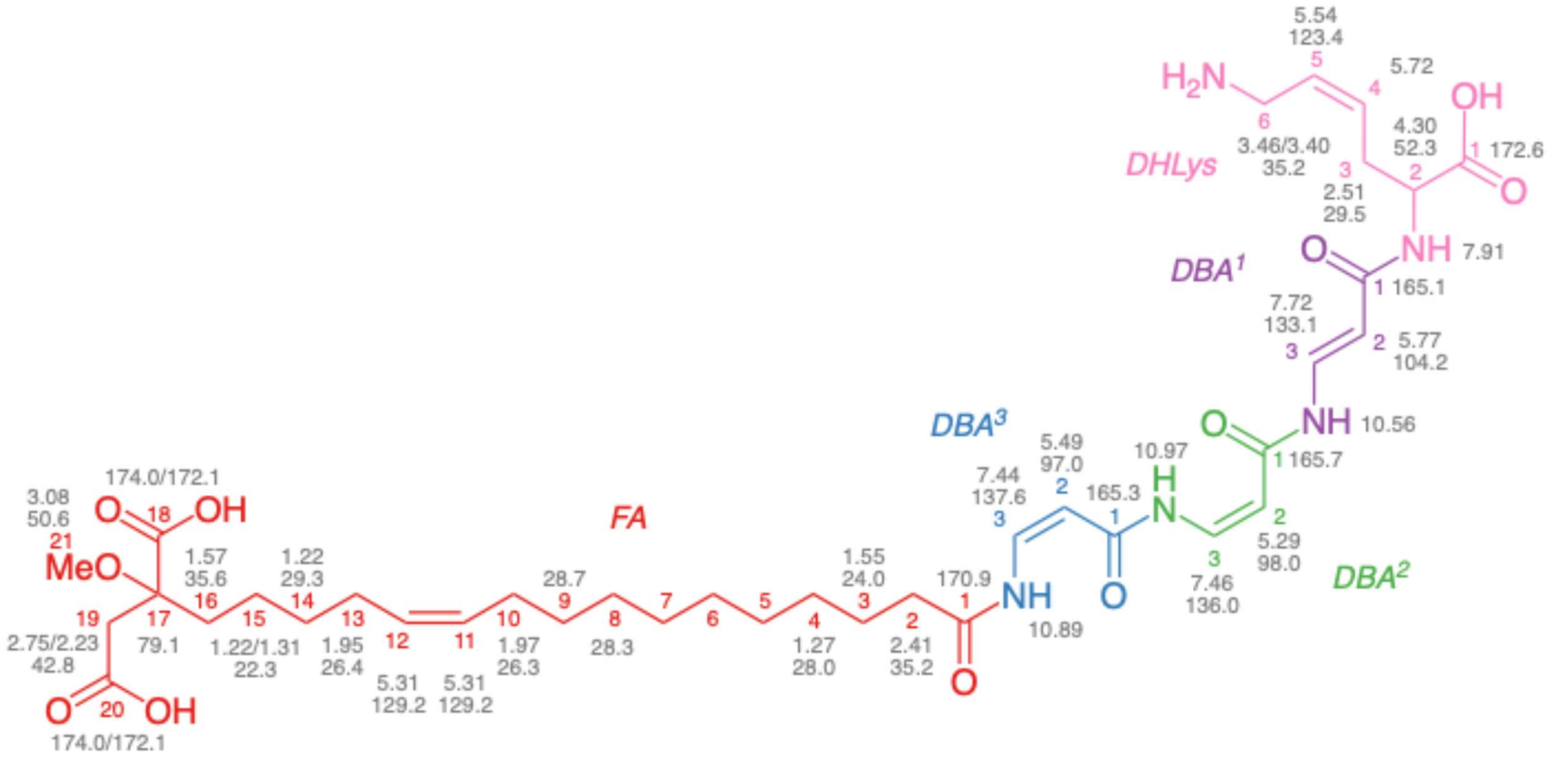


**Figure S6.** Summary of the $^{1}H$ and $^{13}C$-NMR chemical shift values for AFC-A (**1,** DMSO-$d_6$, 600 and 1200 MHz).

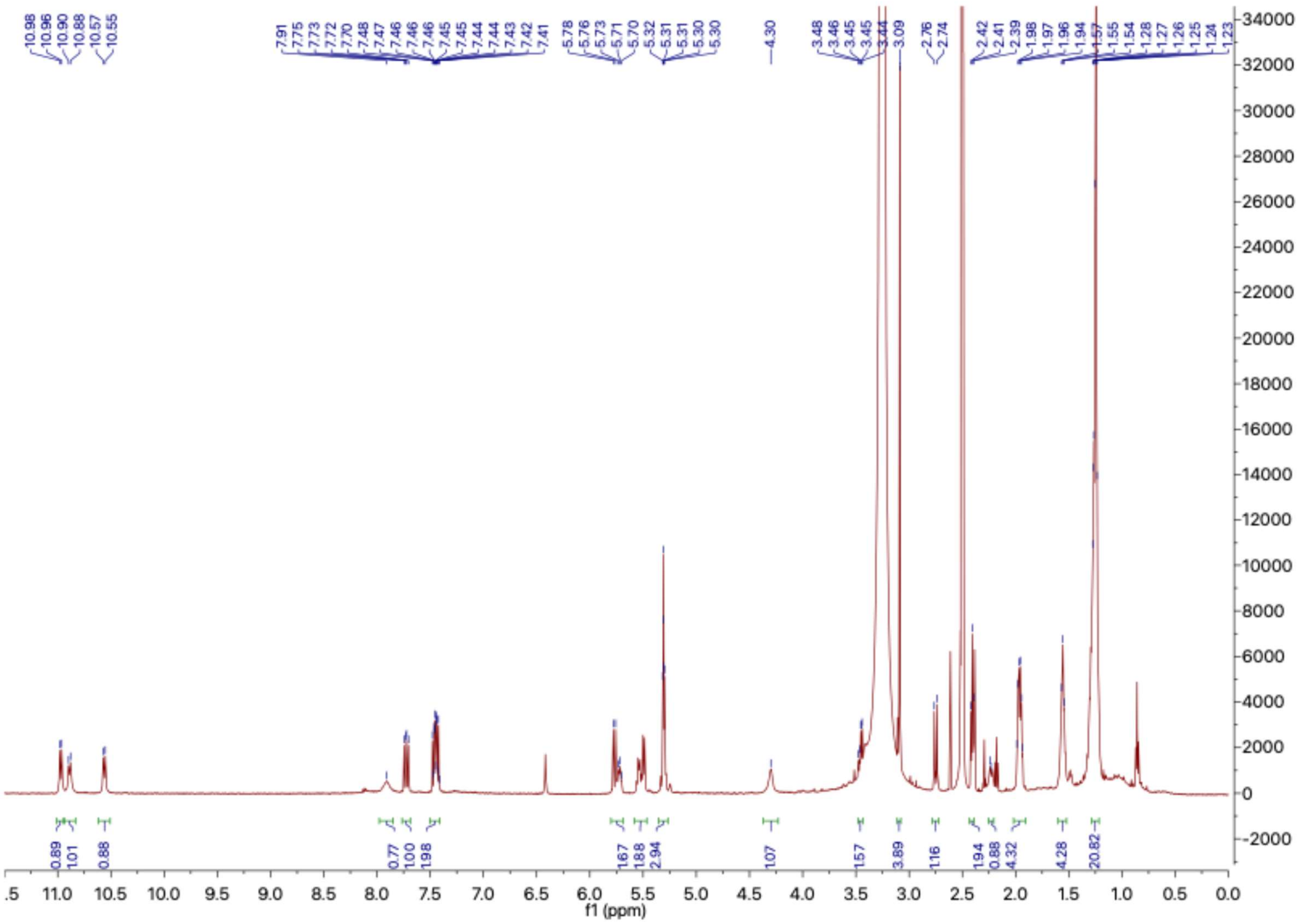

**Figure S7.** $^{1}$H-NMR Spectrum of AFC-A (**1**), (DMSO-$d_6$, 600 MHz, 313K)

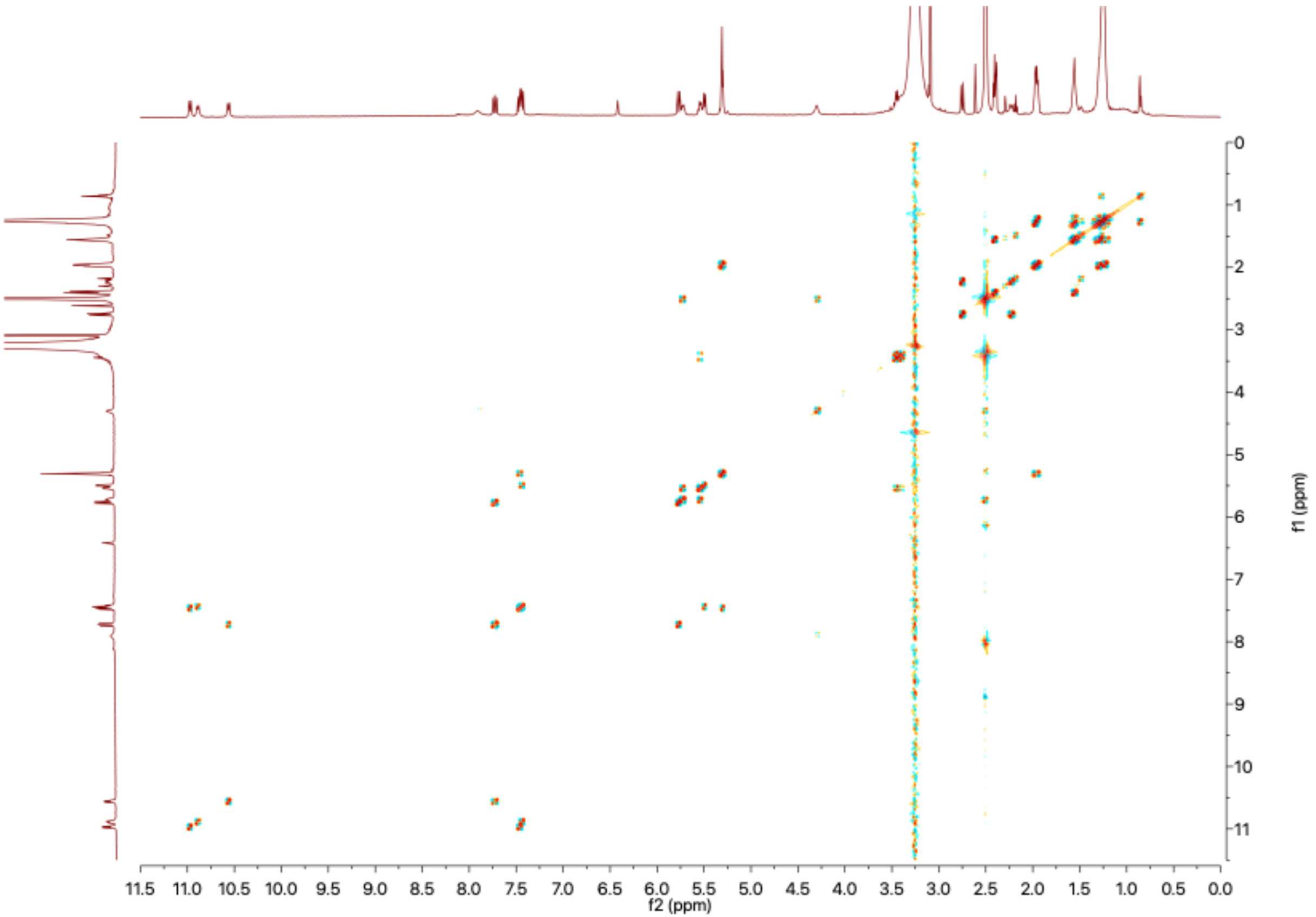


**Figure S8.** $^{1}H$-$^{1}H$-COSY Spectrum of AFC-A (**1**), (DMSO-$d_6$, 600 MHz, 313K)

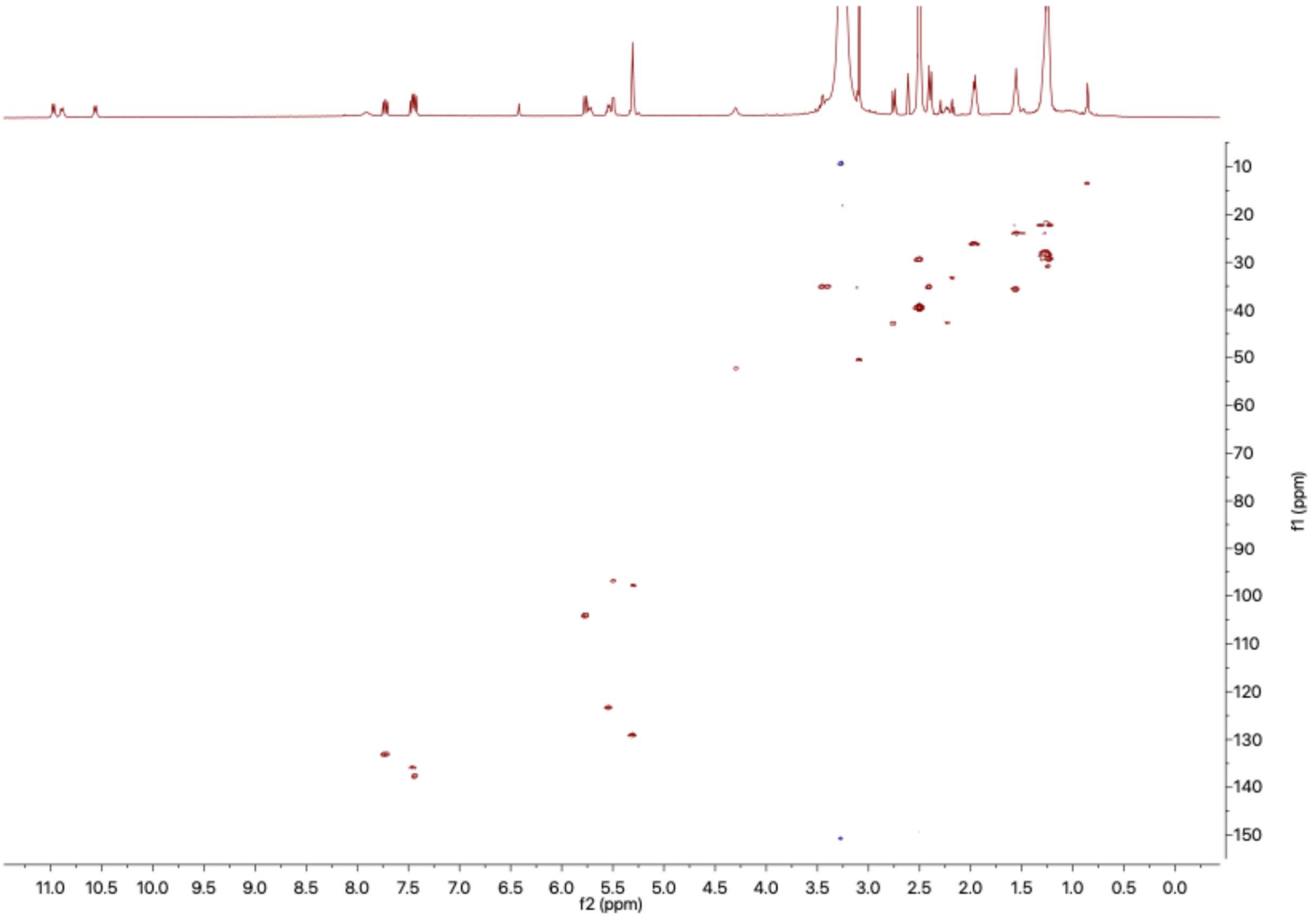


**Figure S9.** $^{1}H$-$^{13}C$-HSQC Spectrum of AFC-A (**1**), (DMSO-$d_6$, 600 MHz, 313K)

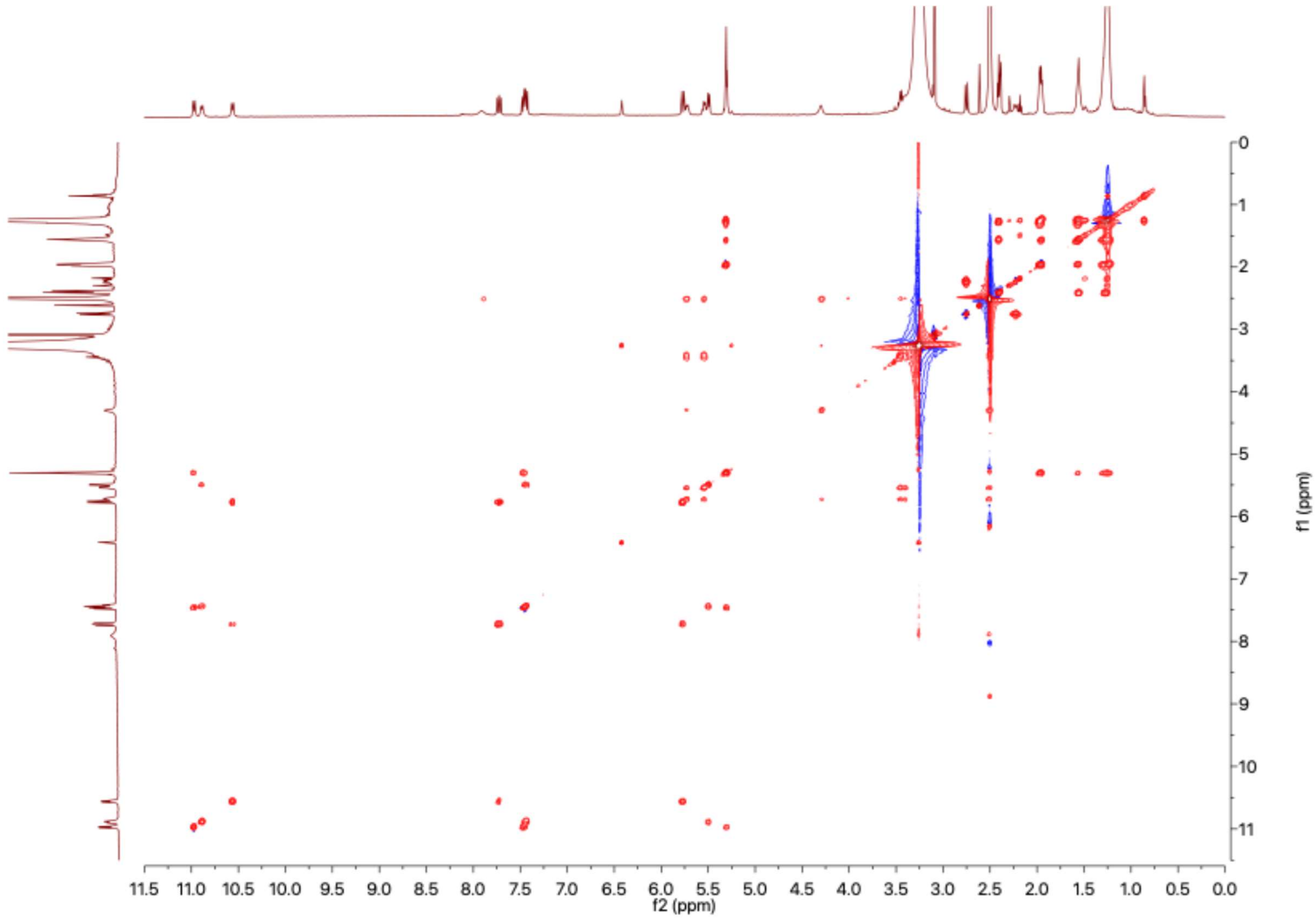


**Figure S10.** $^{1}H$-$^{1}H$-TOCSY Spectrum (100 ms) of AFC-A (**1**), (DMSO-$d_6$, 600 MHz, 313K)

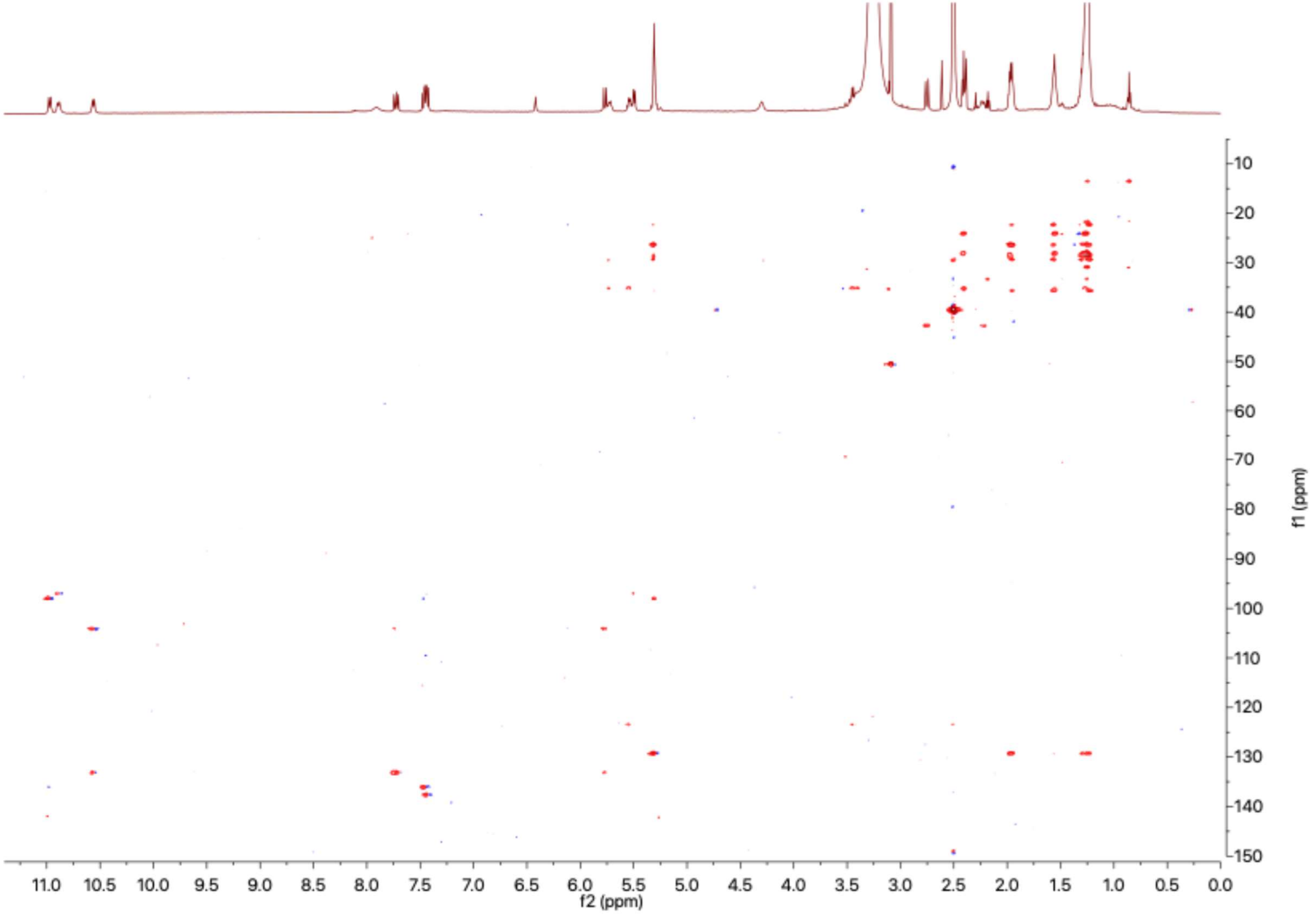


**Figure S11.** $^{1}$H-$^{13}$C-HSQC-TOCSY Spectrum (60 ms) of AFC-A (**1**), (DMSO-$d_6$, 600 MHz, 313K)

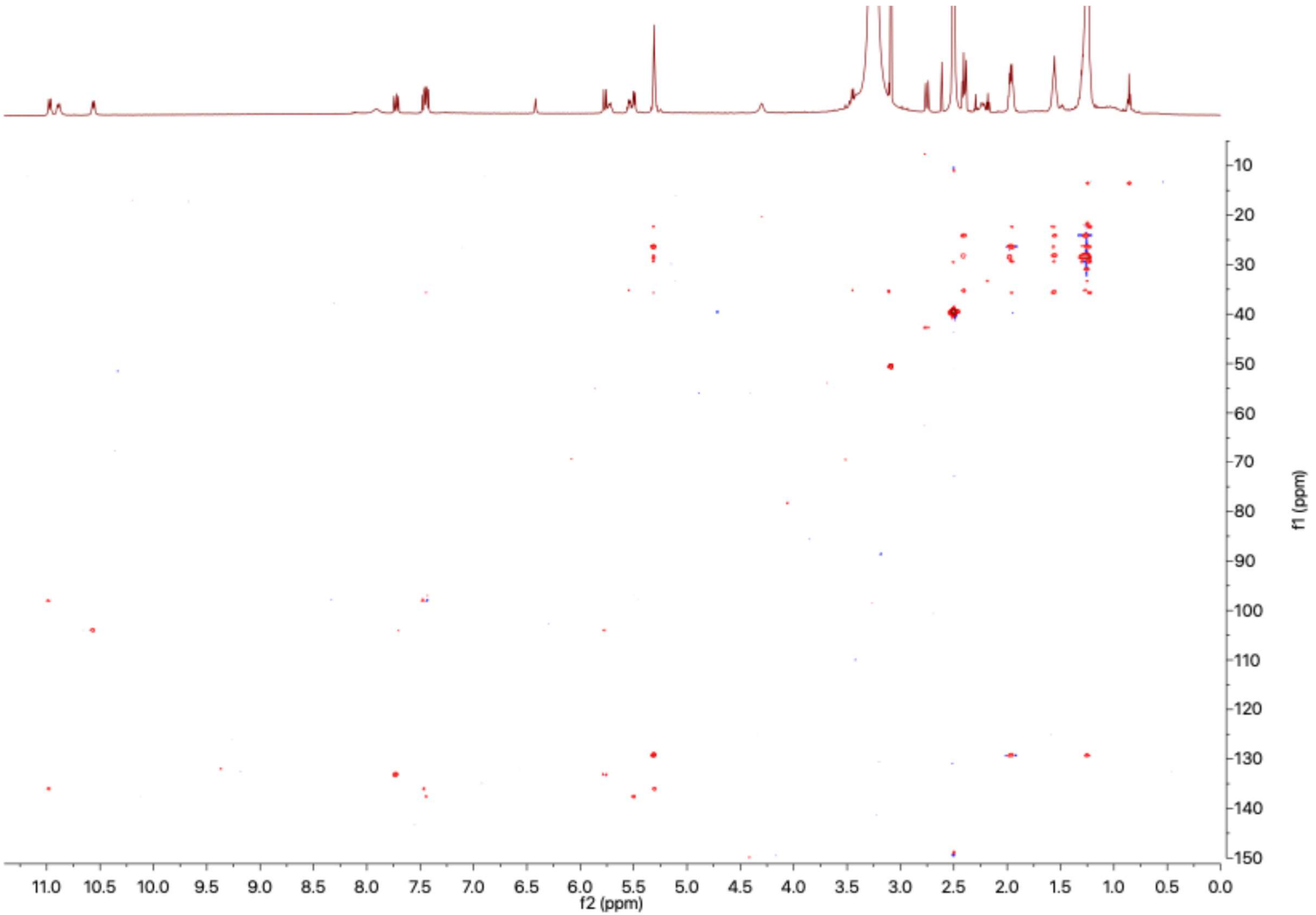


**Figure S12.** $^{1}H$-$^{13}C$-HSQC-TOCSY Spectrum (90 ms) of AFC-A (**1**), (DMSO-$d_6$, 600 MHz, 313K)

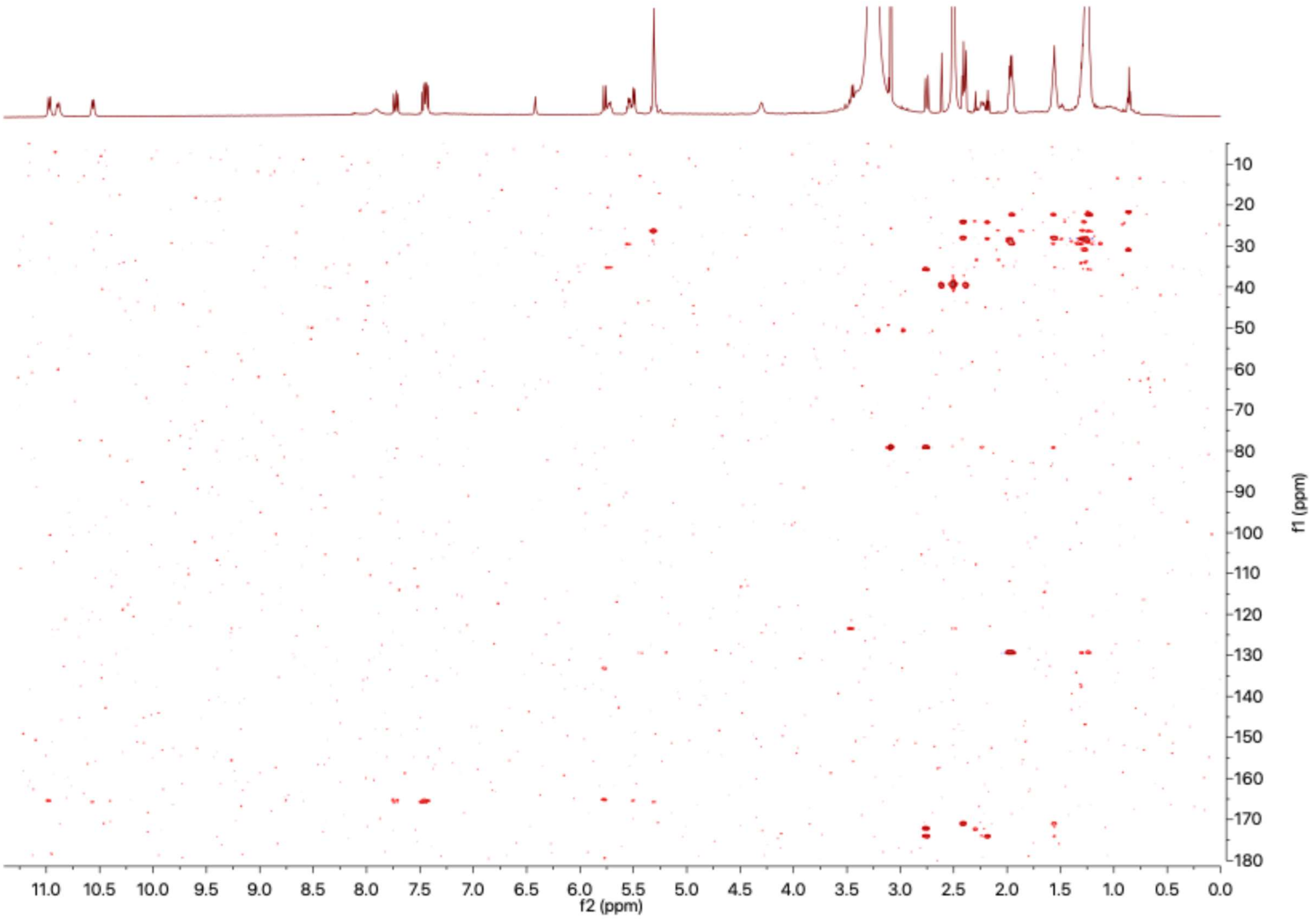


**Figure S13.** $^{1}H$-$^{13}C$-HMBC Spectrum of AFC-A (**1**), (DMSO-$d_6$, 600 MHz, 313K)

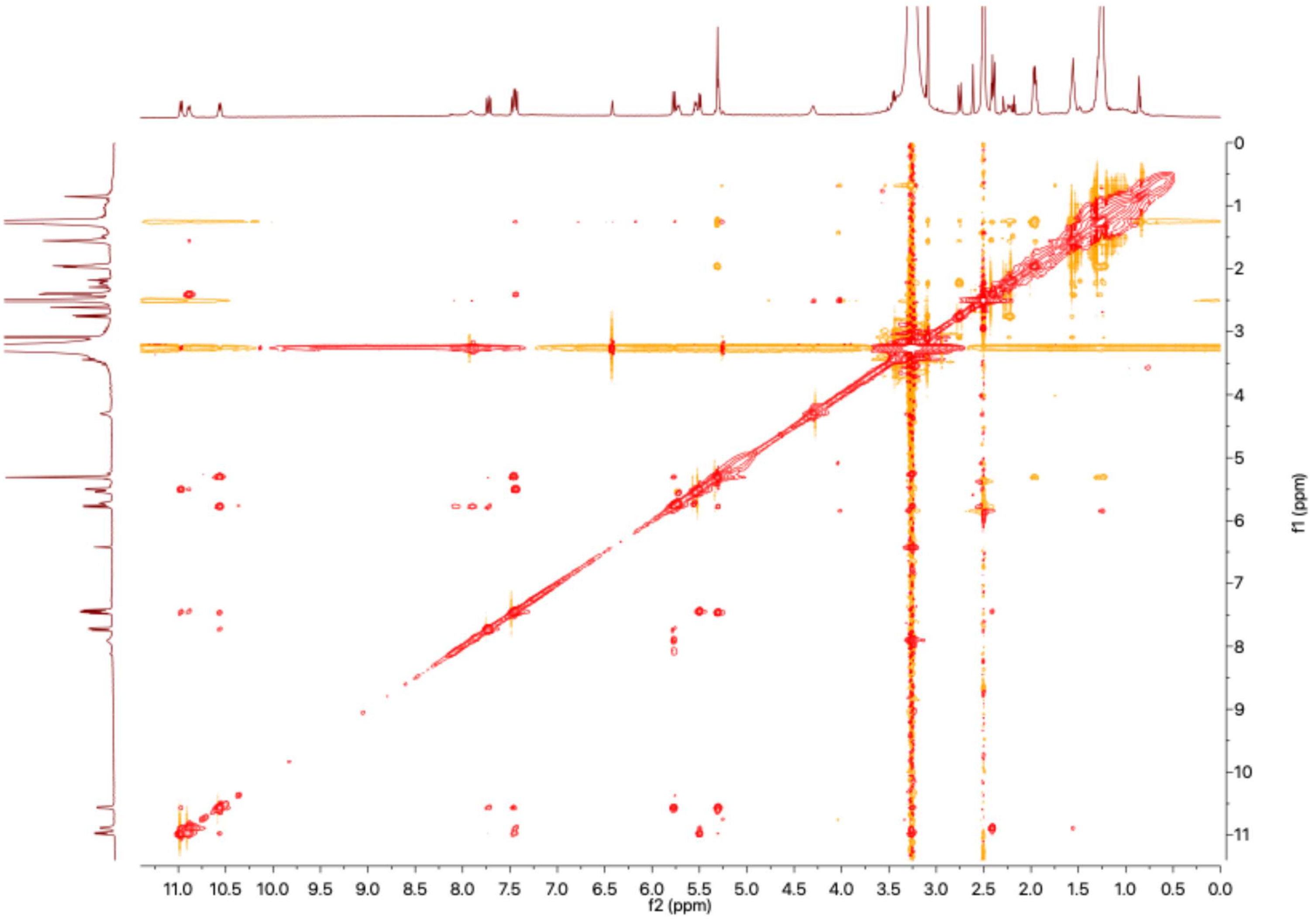


**Figure S14.** $^{1}$H-$^{1}$H-NOESY Spectrum of AFC-A (**1**), (DMSO-$d_6$, 600 MHz, 313K)

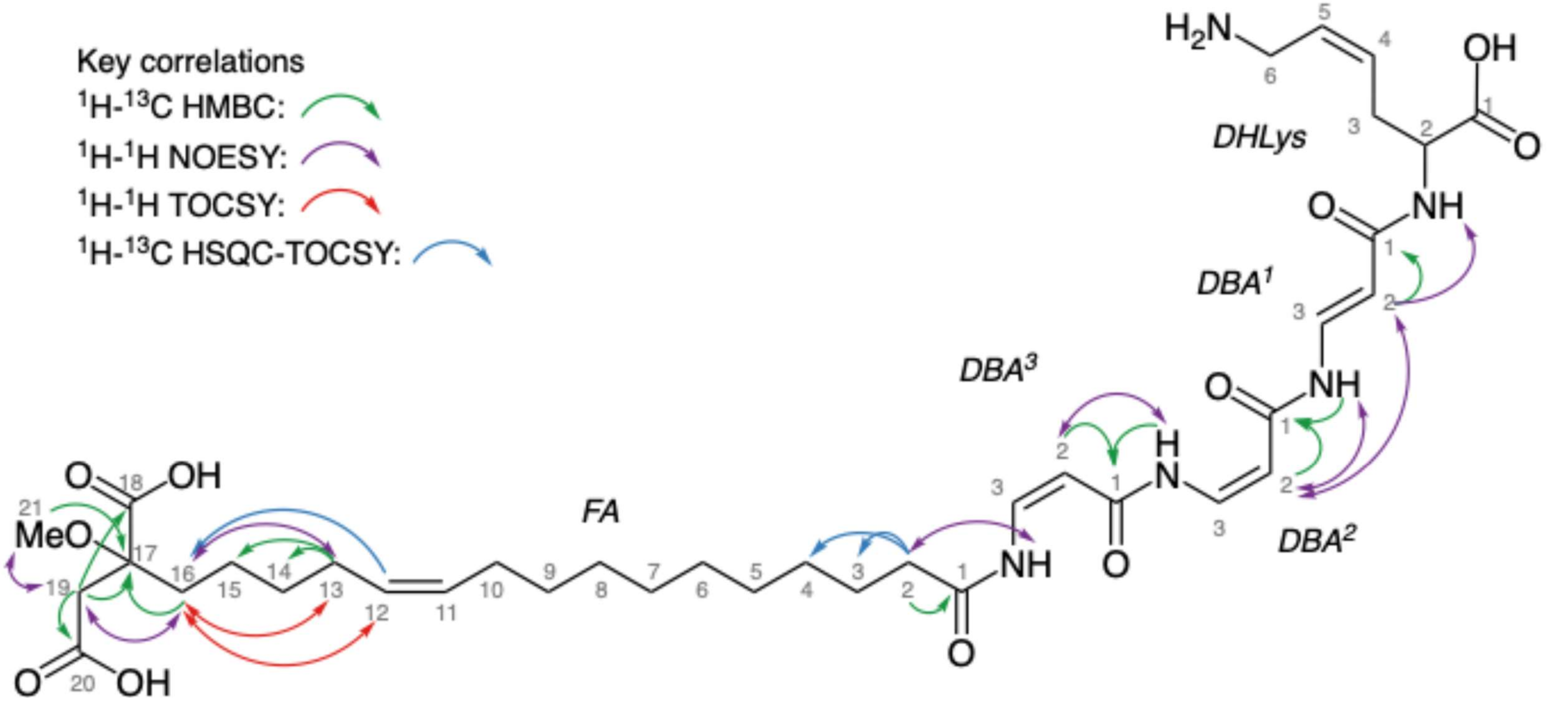


**Figure S15.** Key correlations for the structural elucidation of AFC-A (**1**).

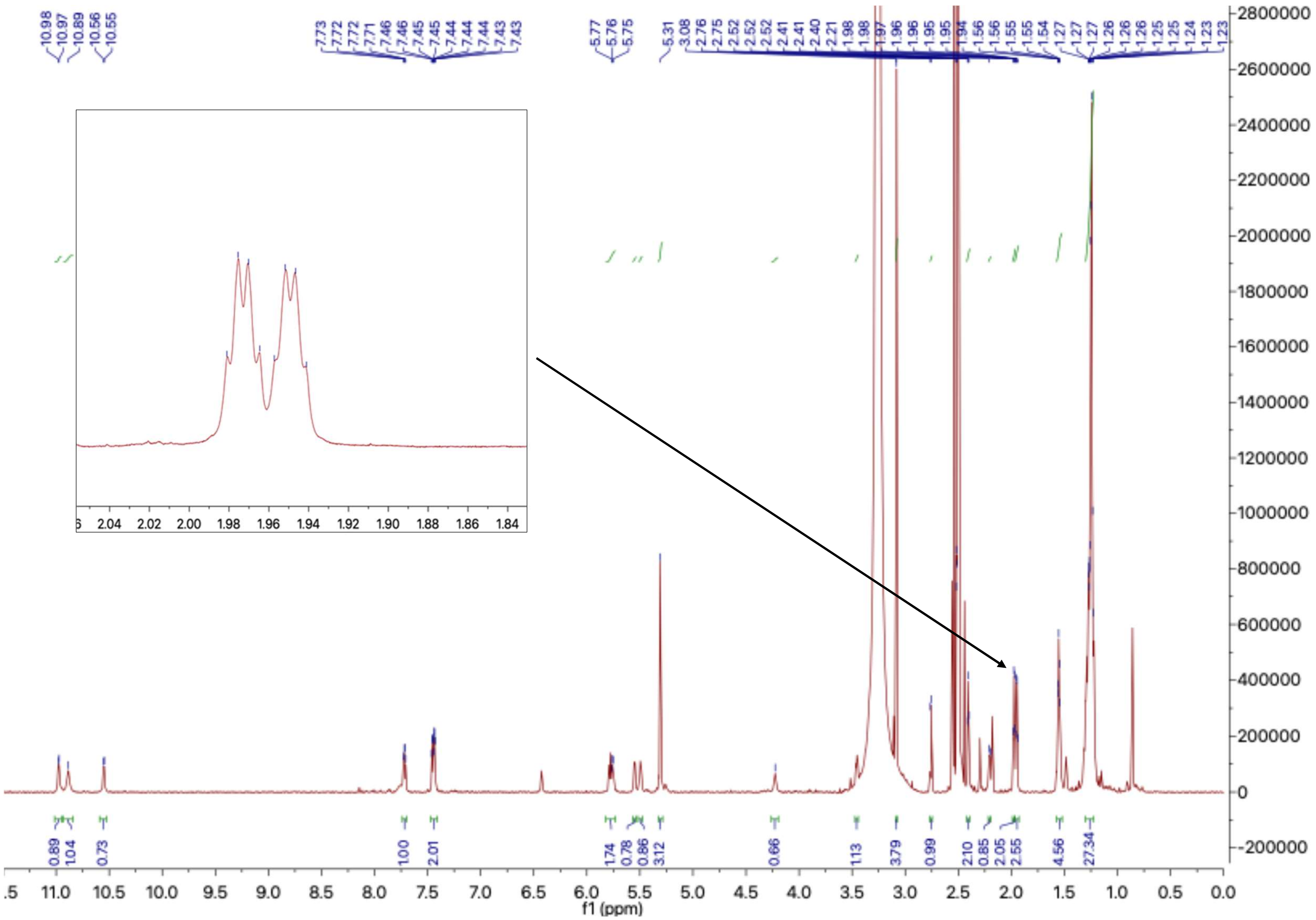

**Figure S16.** [1]H-NMR Spectrum of AFC-A (**1**), (DMSO-$d_6$, 1.2 GHz, 313K) with a highlighted region showing the resonances of protons H-10 and H-13.

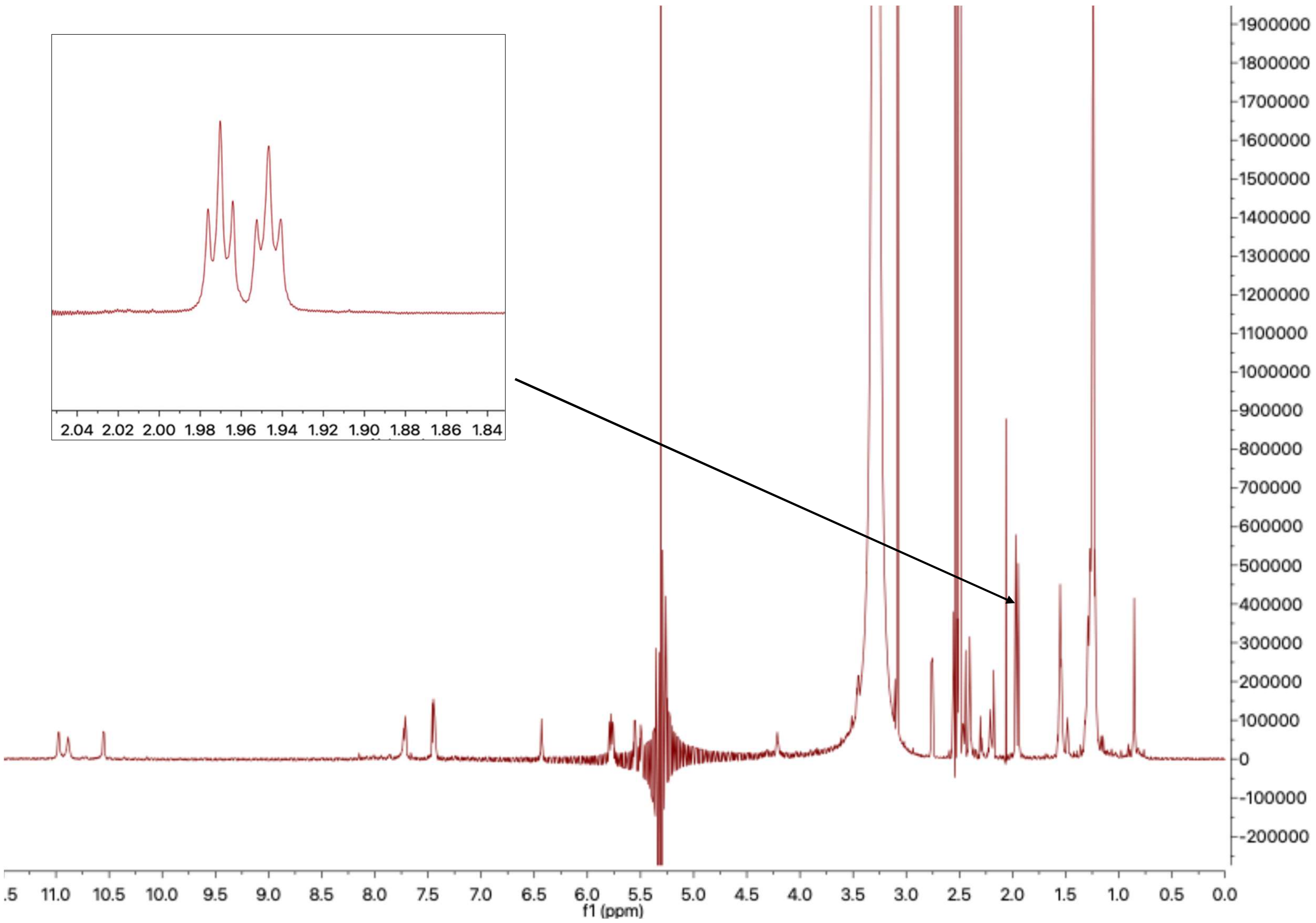


**Figure S17.** [1]H-NMR Spectrum recorded with homodecoupling at 5.3 ppm of AFC-A (**1**), (DMSO-$d_6$, 1.2 GHz, 313K) with a highlighted region showing the resonances of protons H-10 and H-13.

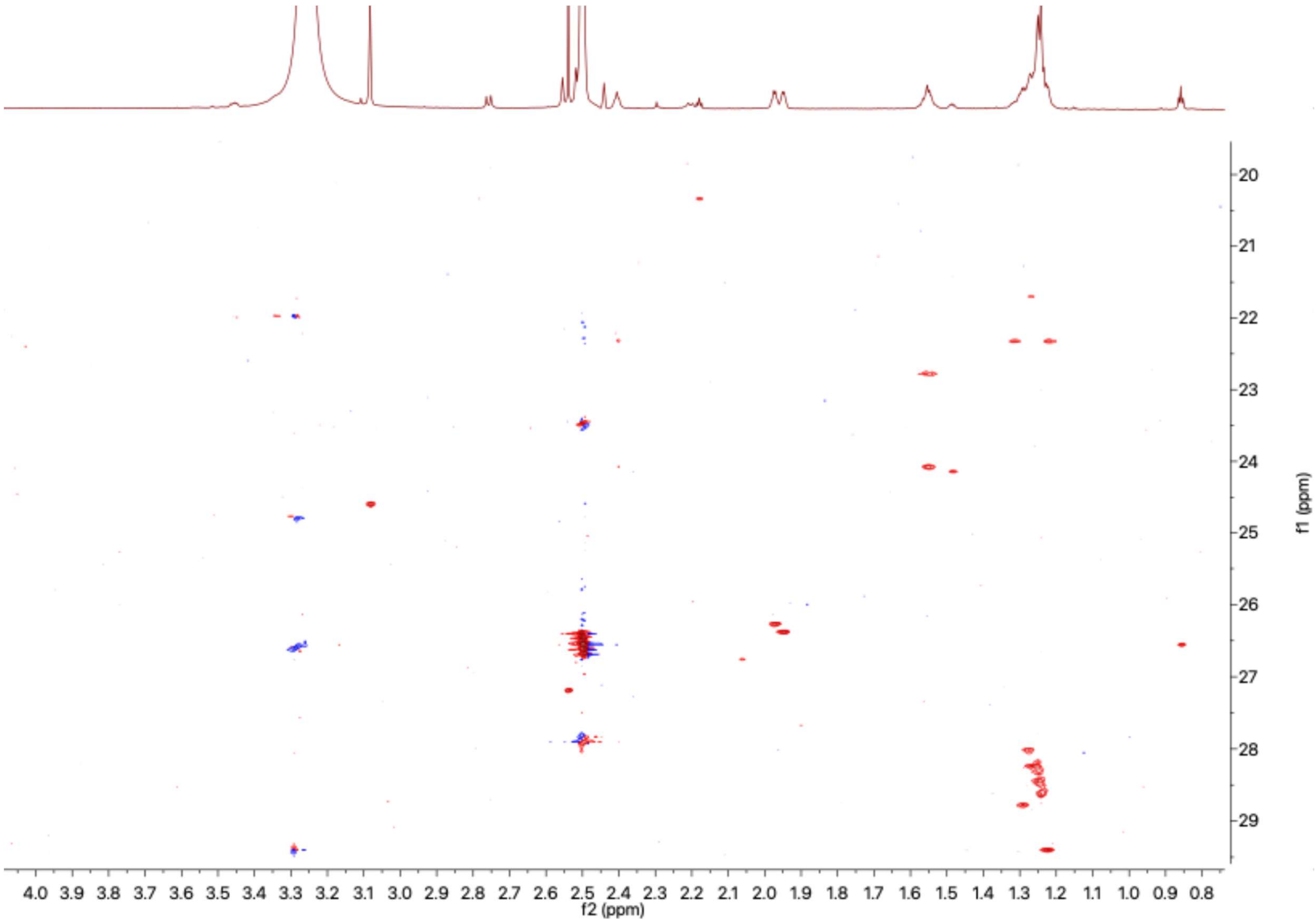


**Figure S18.** $^{1}H$-$^{13}C$-HSQC Spectrum of AFC-A (**1**) in high resolution, (DMSO-$d_6$, 1.2 GHz, 313K)

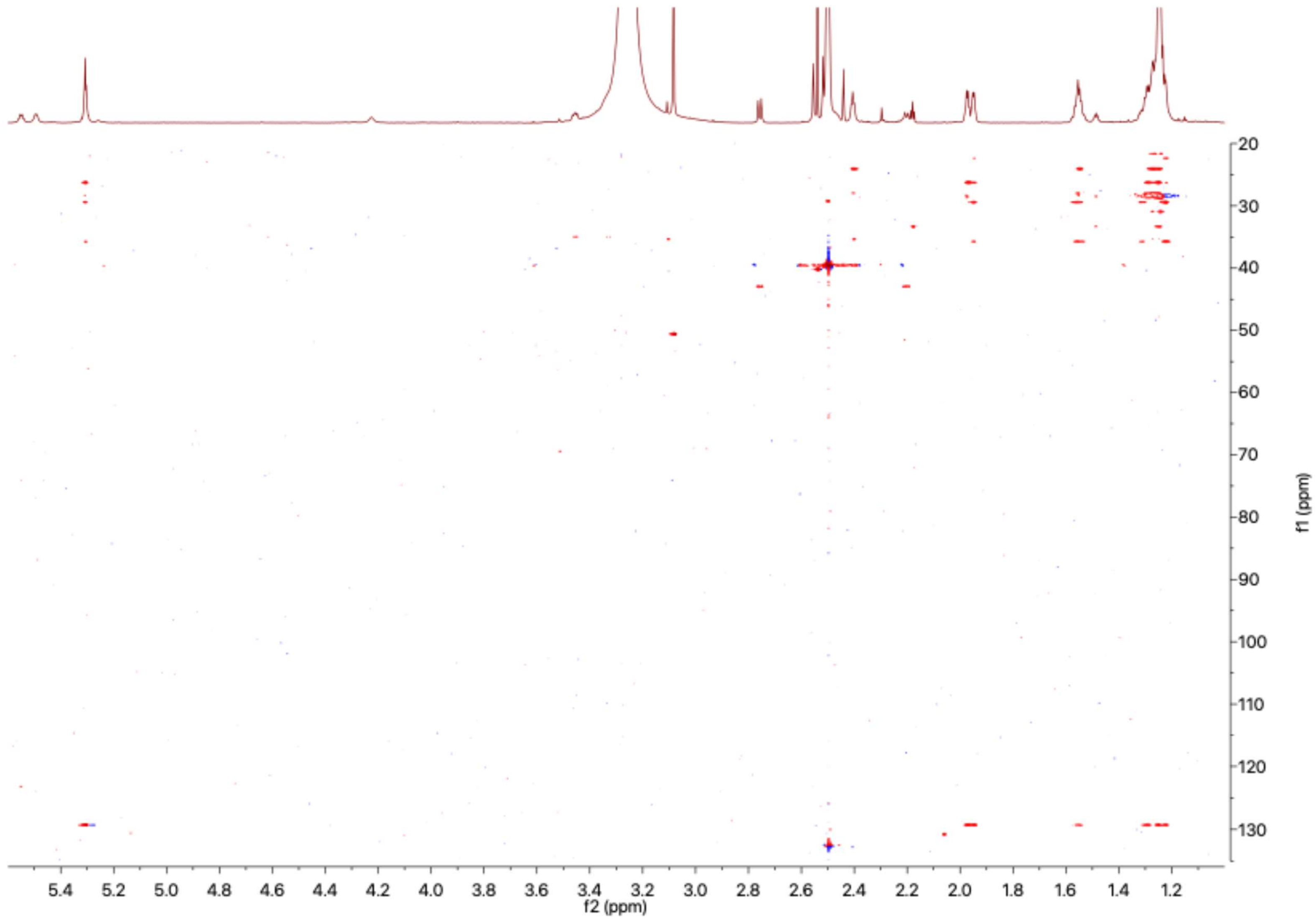


**Figure S19.** $^{1}H$-$^{13}C$-HSQC-TOCSY Spectrum (100 ms) of AFC-A (**1**) with decoupling at 5.3 ppm, (DMSO-$d_6$, 1.2 GHz, 313K) with the part of the spectrum showing the key correlations for the fatty acid moiety enlarged.

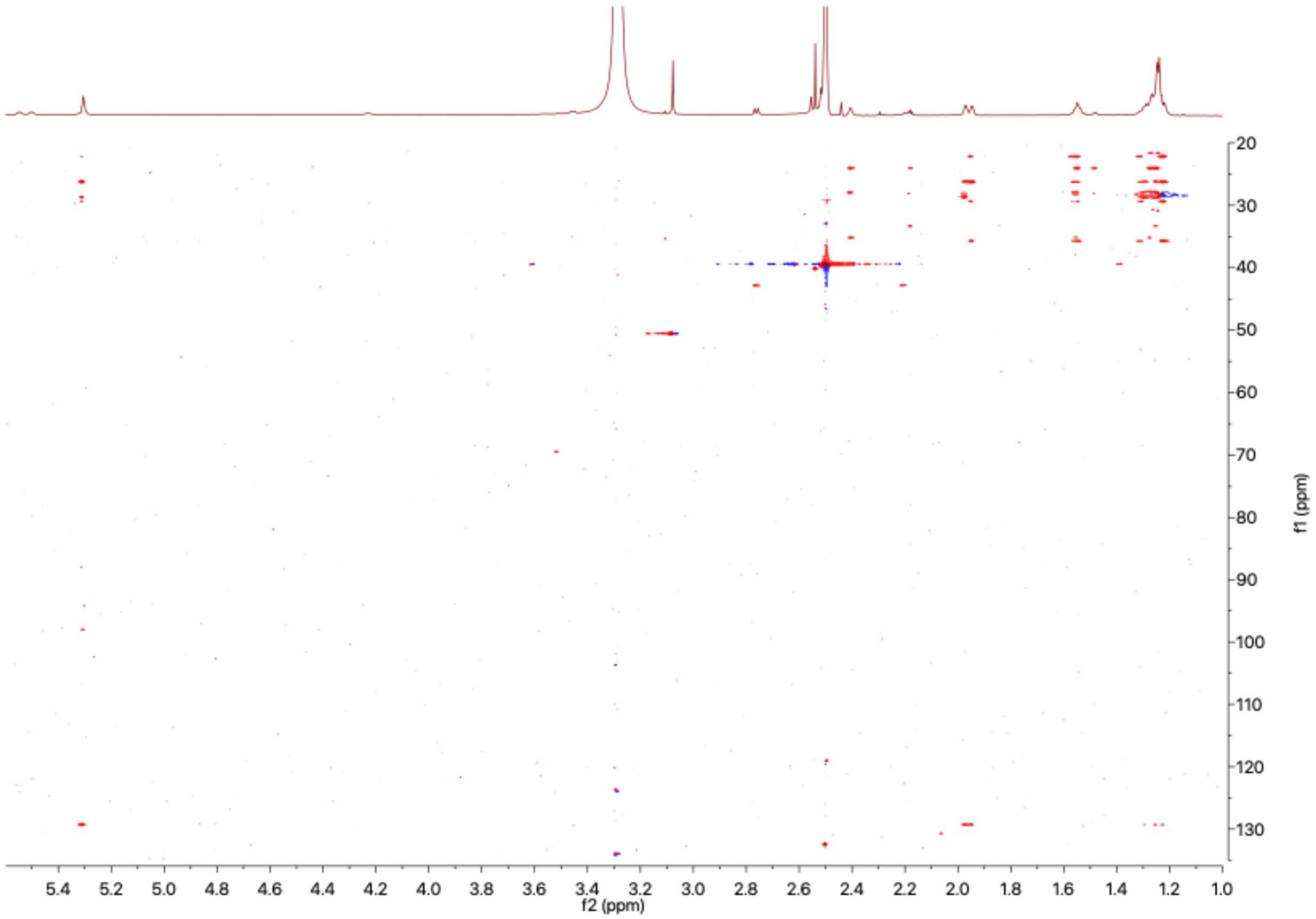


**Figure S20.** $^{1}H$-$^{13}C$-HSQC-TOCSY Spectrum (60 ms) of AFC-A (**1**) with decoupling at 5.3 ppm, (DMSO-$d_6$, 1.2 GHz, 313K) the part of the spectrum showing the key correlations for the fatty acid moiety has been enlarged.

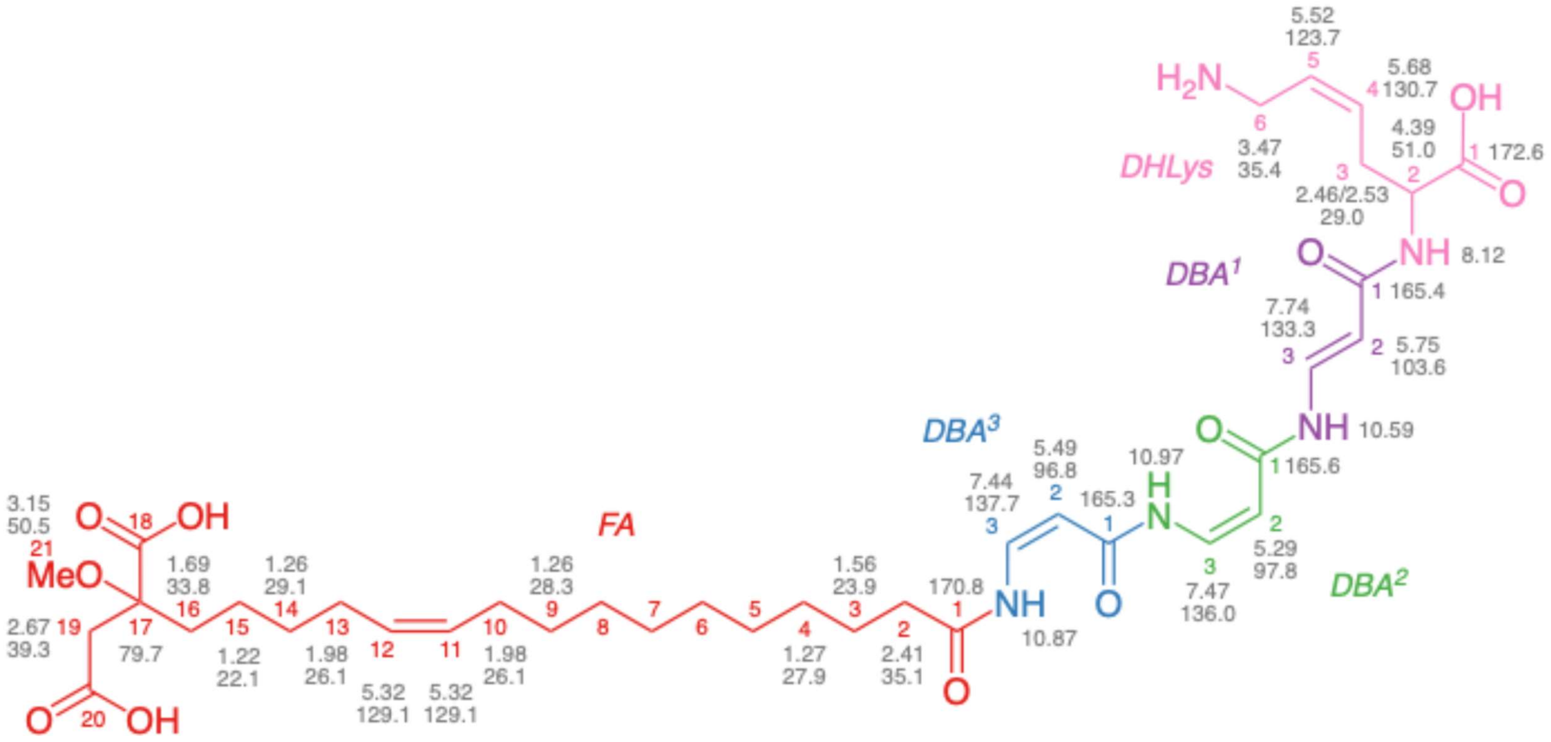


**Figure S21.** Summary of chemical shifts for AFC-A (**1**) using the sample obtained from LB medium.

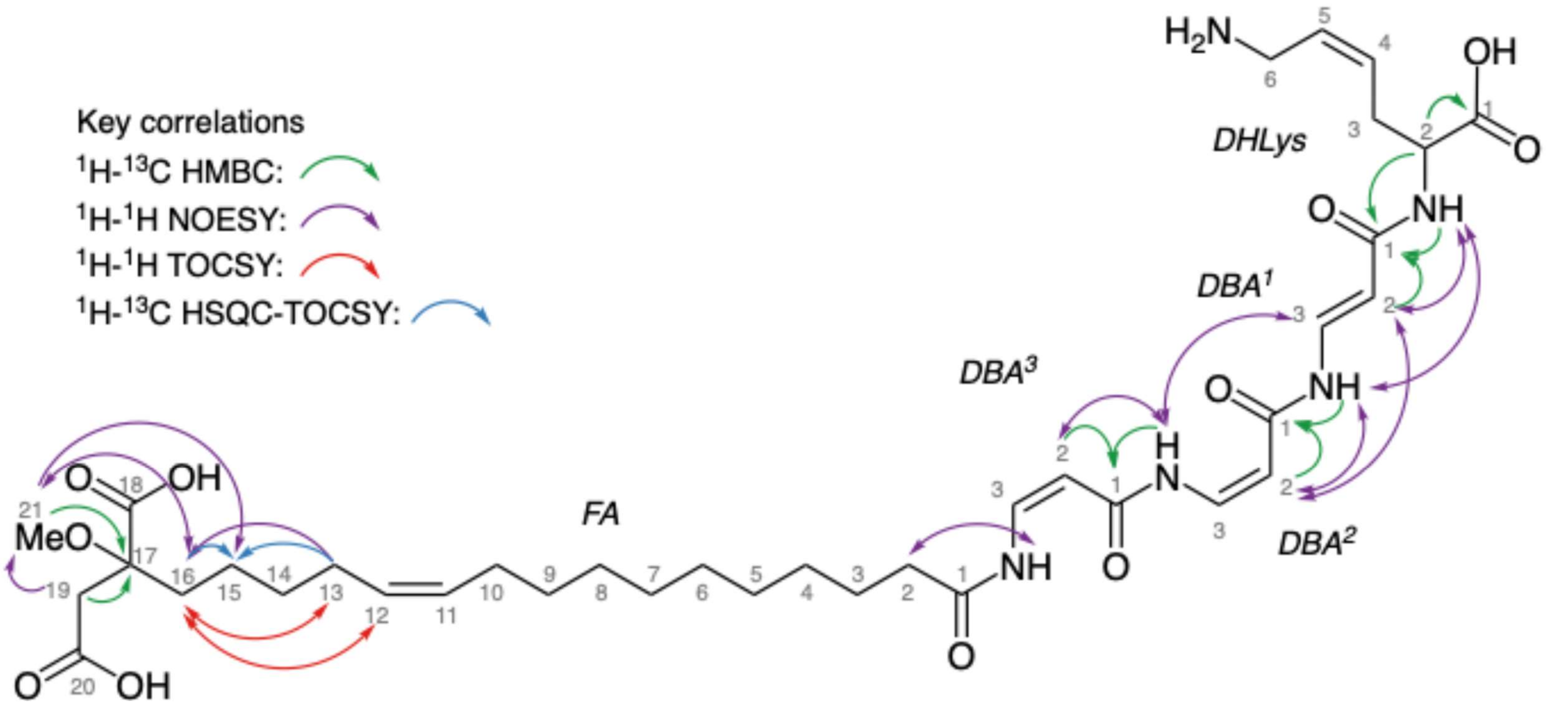


**Figure S22.** Key correlations used for the structure elucidation of AFC-A (**1**) using the sample obtained from LB medium.

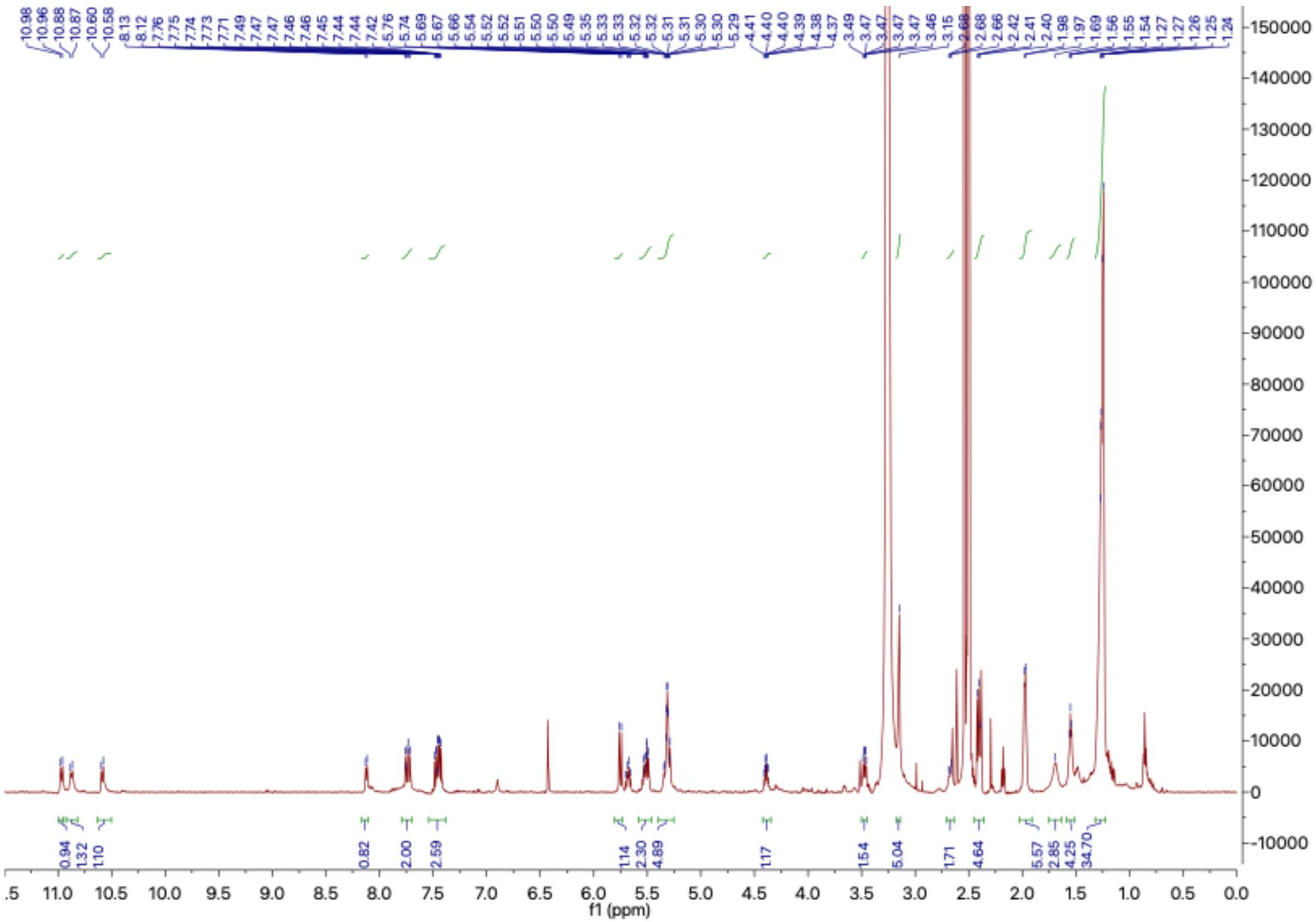

**Figure S23.** [1]H-NMR Spectrum of AFC-A (**1**) obtained from LB medium, (DMSO-$d_6$, 600 MHz)

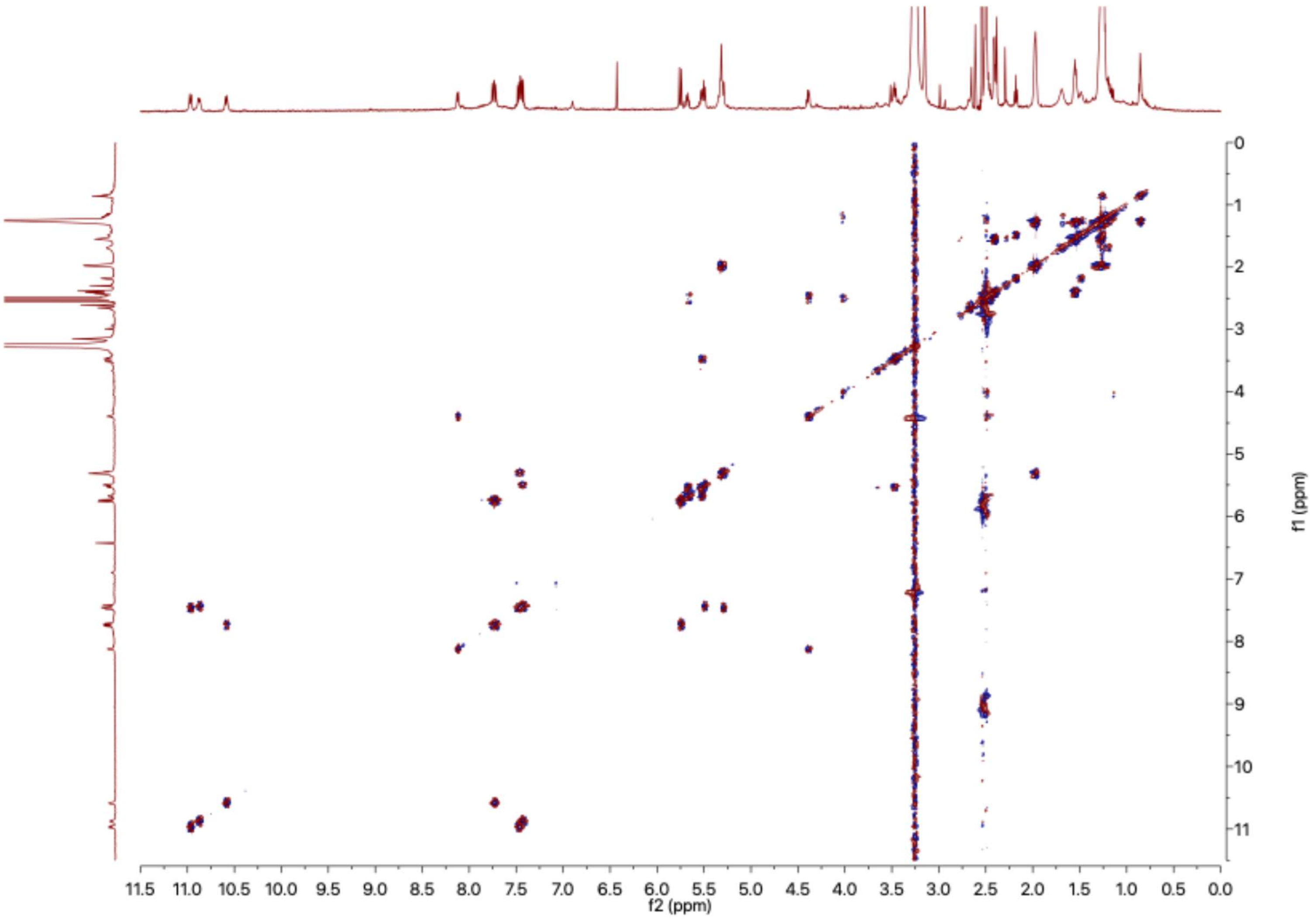


**Figure S24.** $^{1}H$-$^{1}H$-COSY Spectrum of AFC-A (**1**) obtained from LB medium, (DMSO-$d_6$, 600 MHz, 313K)

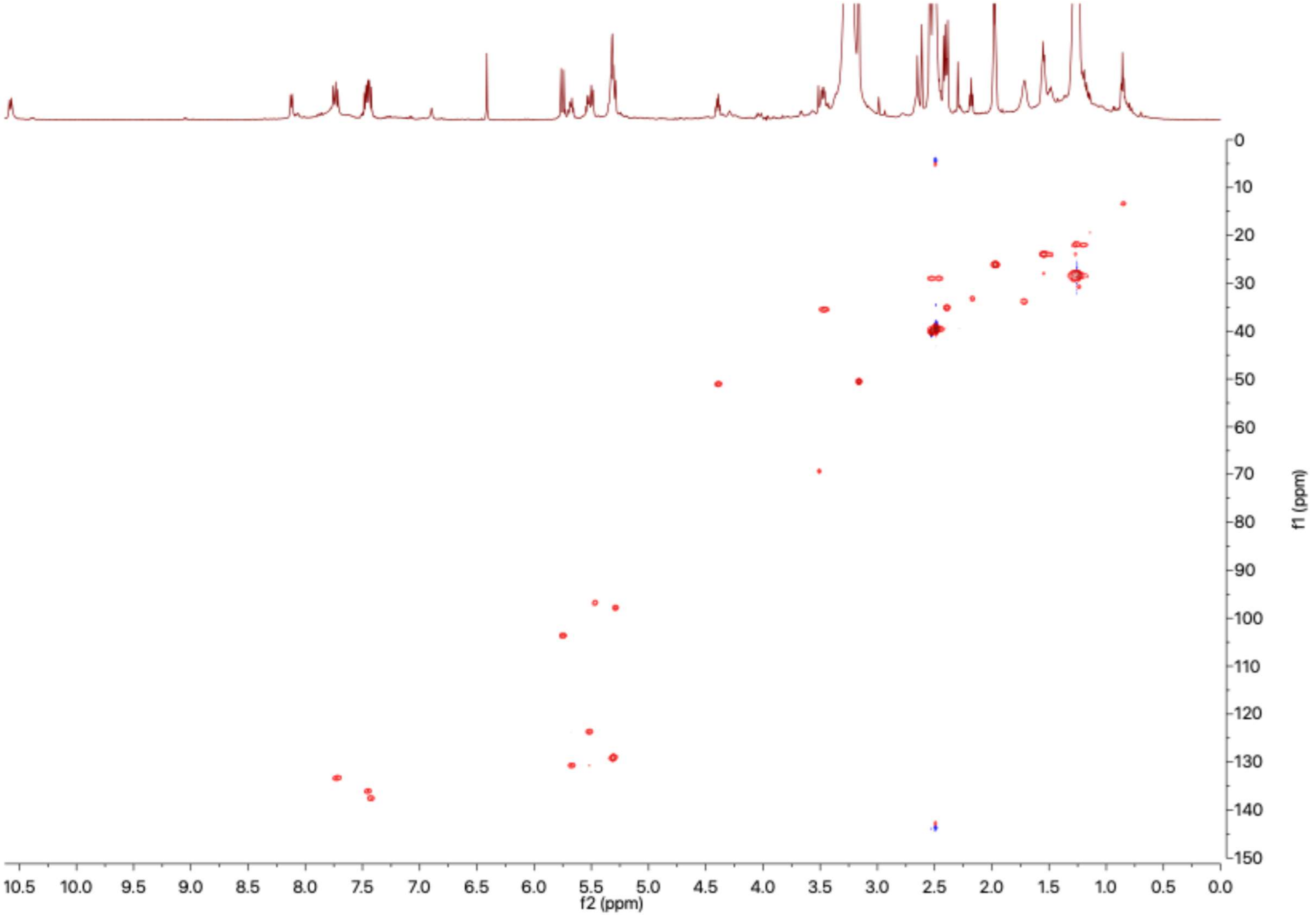


**Figure S25.** $^{1}H$-$^{13}C$-HSQC Spectrum of AFC-A (**1**) obtained from LB medium, (DMSO-$d_6$, 600 MHz, 313K)

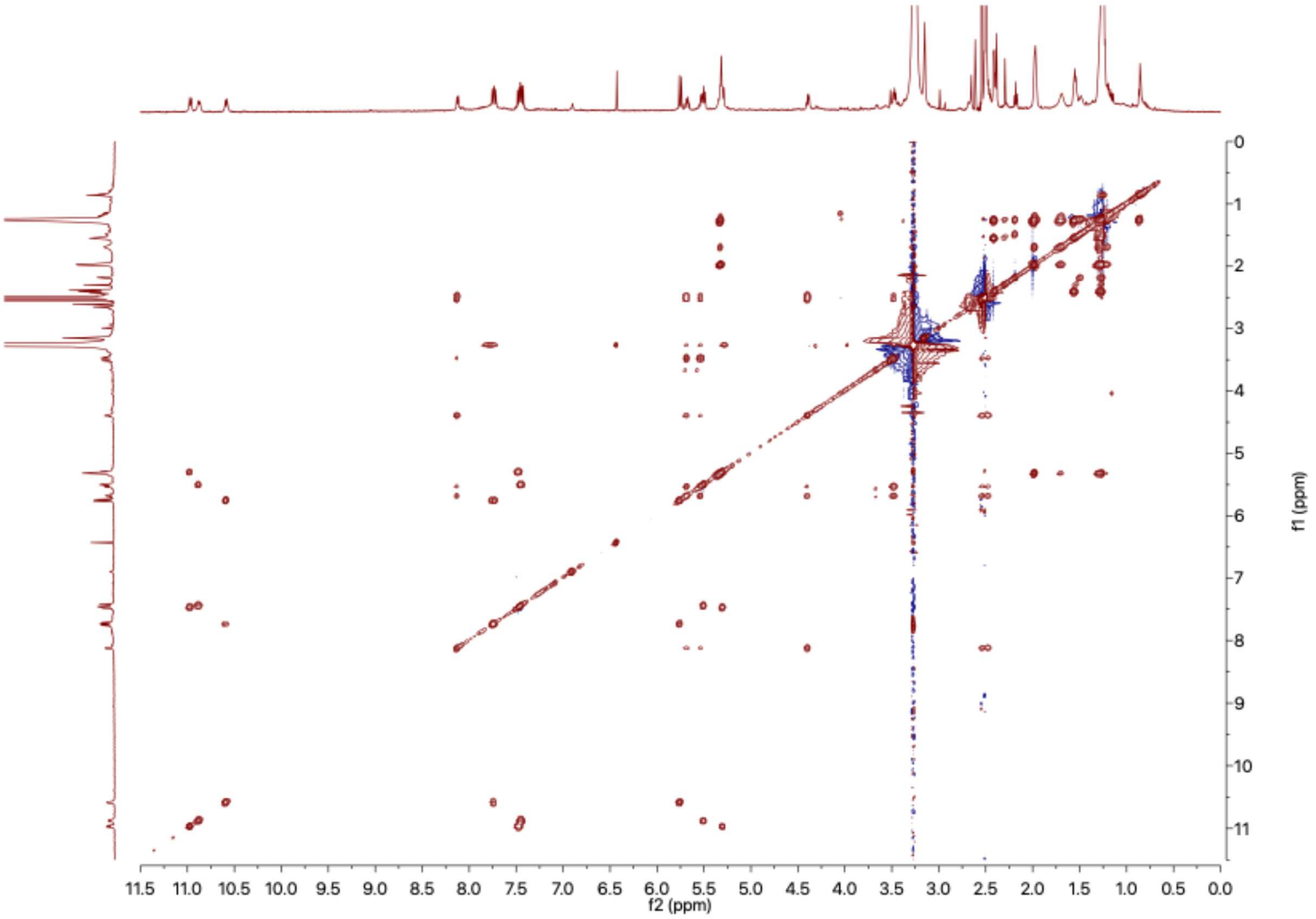


**Figure S26.** $^{1}H$-$^{1}H$-TOCSY Spectrum of AFC-A (**1**) obtained from LB medium, (DMSO-$d_6$, 600 MHz, 313K)

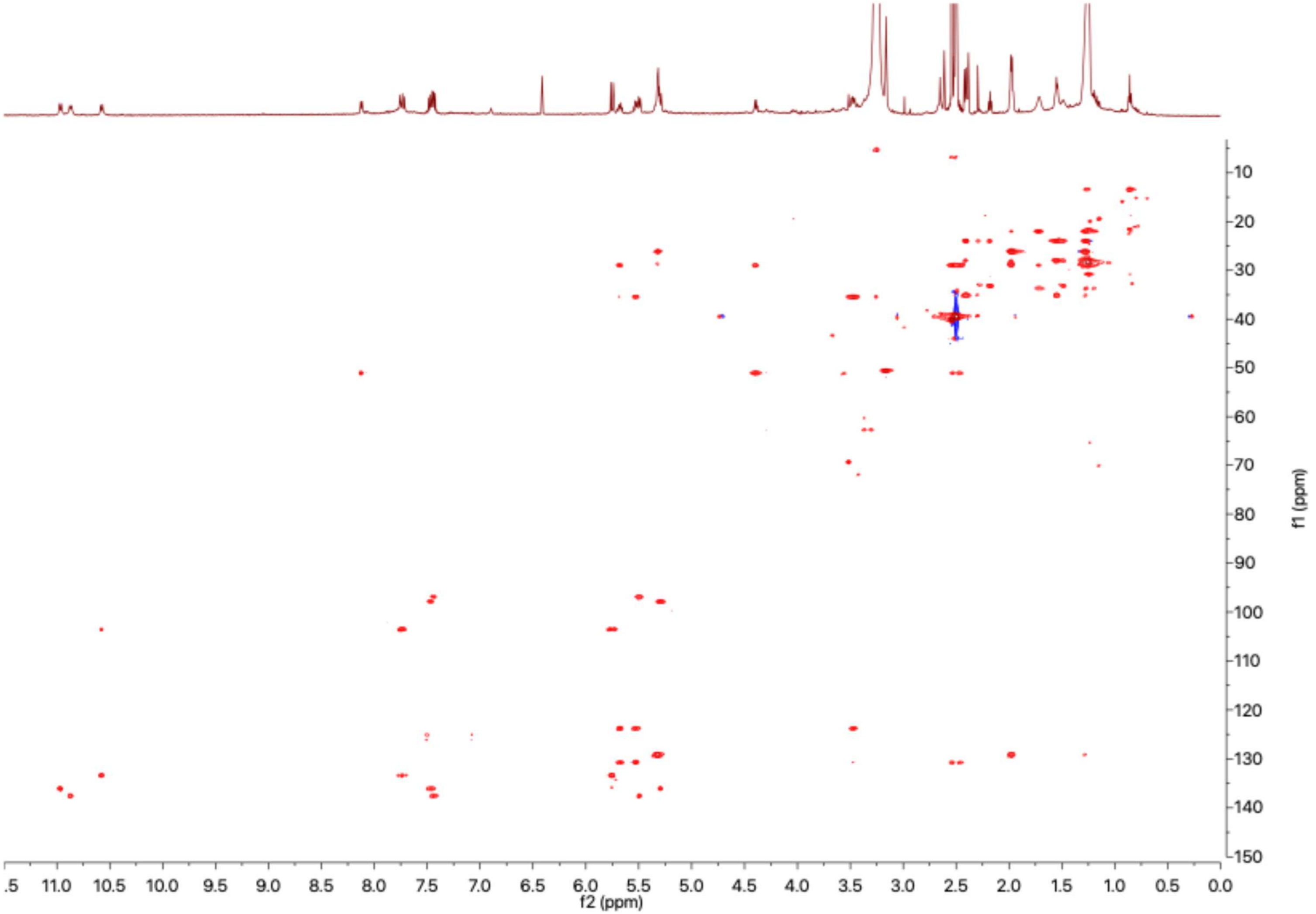


**Figure S27.** $^{1}$H-$^{13}$C-HSQC-TOCSY Spectrum (60 ms) of AFC-A (**1**) obtained from LB medium, (DMSO-$d_6$, 600 MHz, 313K)

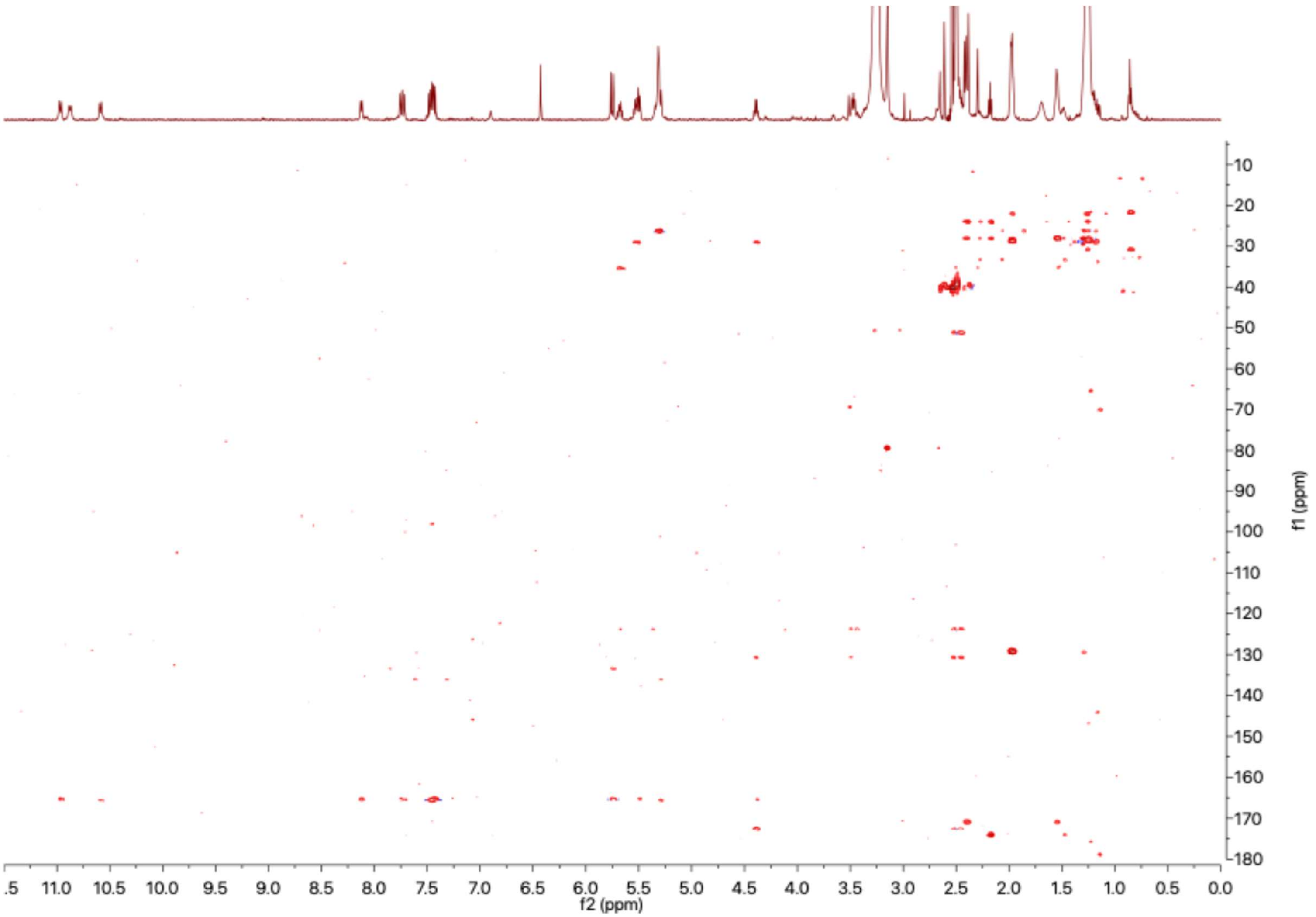


**Figure S28.** $^{1}H$-$^{13}C$-HMBC Spectrum of AFC-A (**1**) obtained from LB medium, (DMSO-$d_6$, 600 MHz, 313K)

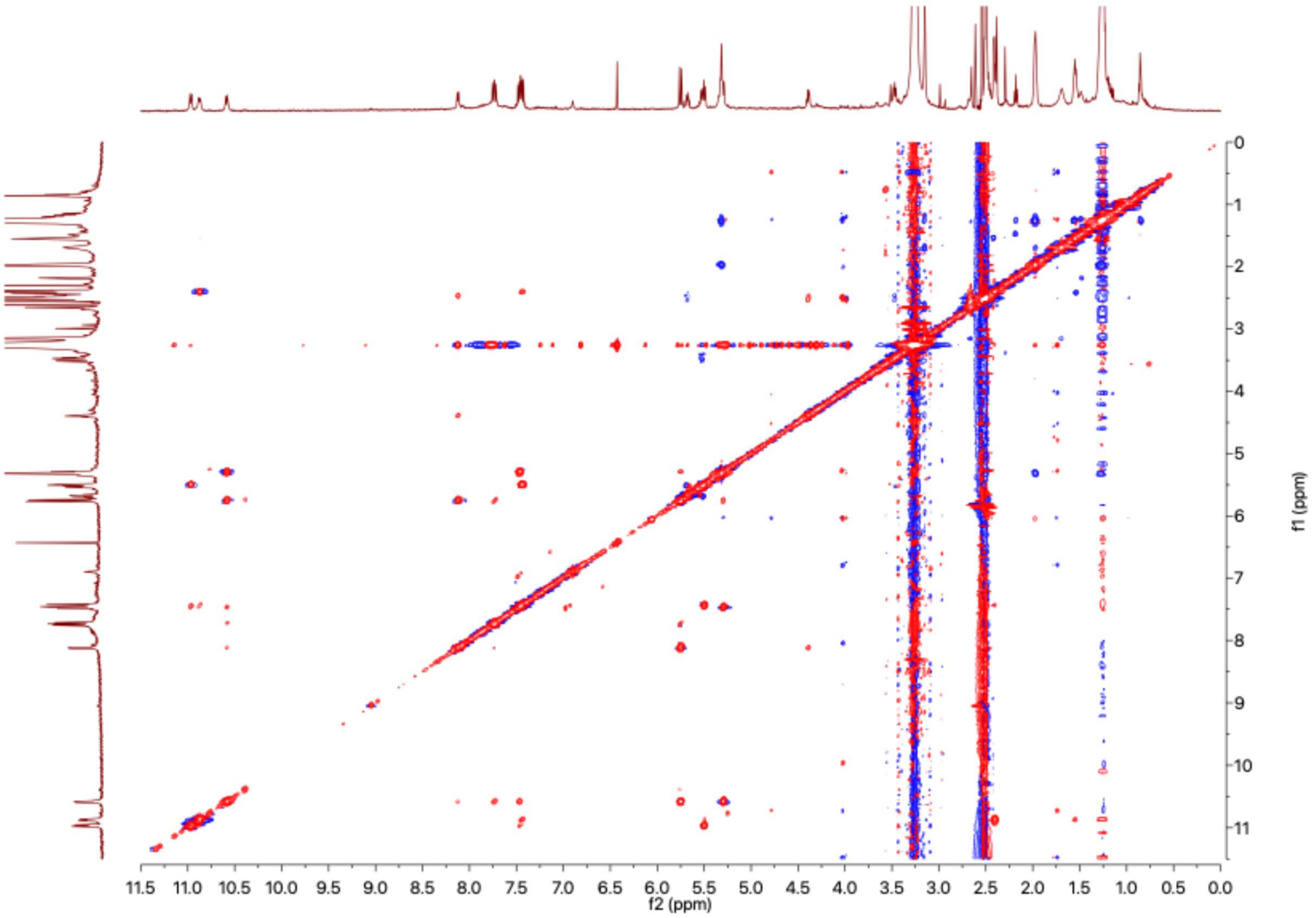


**Figure S29.** $^{1}$H-$^{1}$H-NOESY Spectrum of AFC-A (**1**) obtained from LB medium, (DMSO-$d_6$, 600 MHz, 313K)

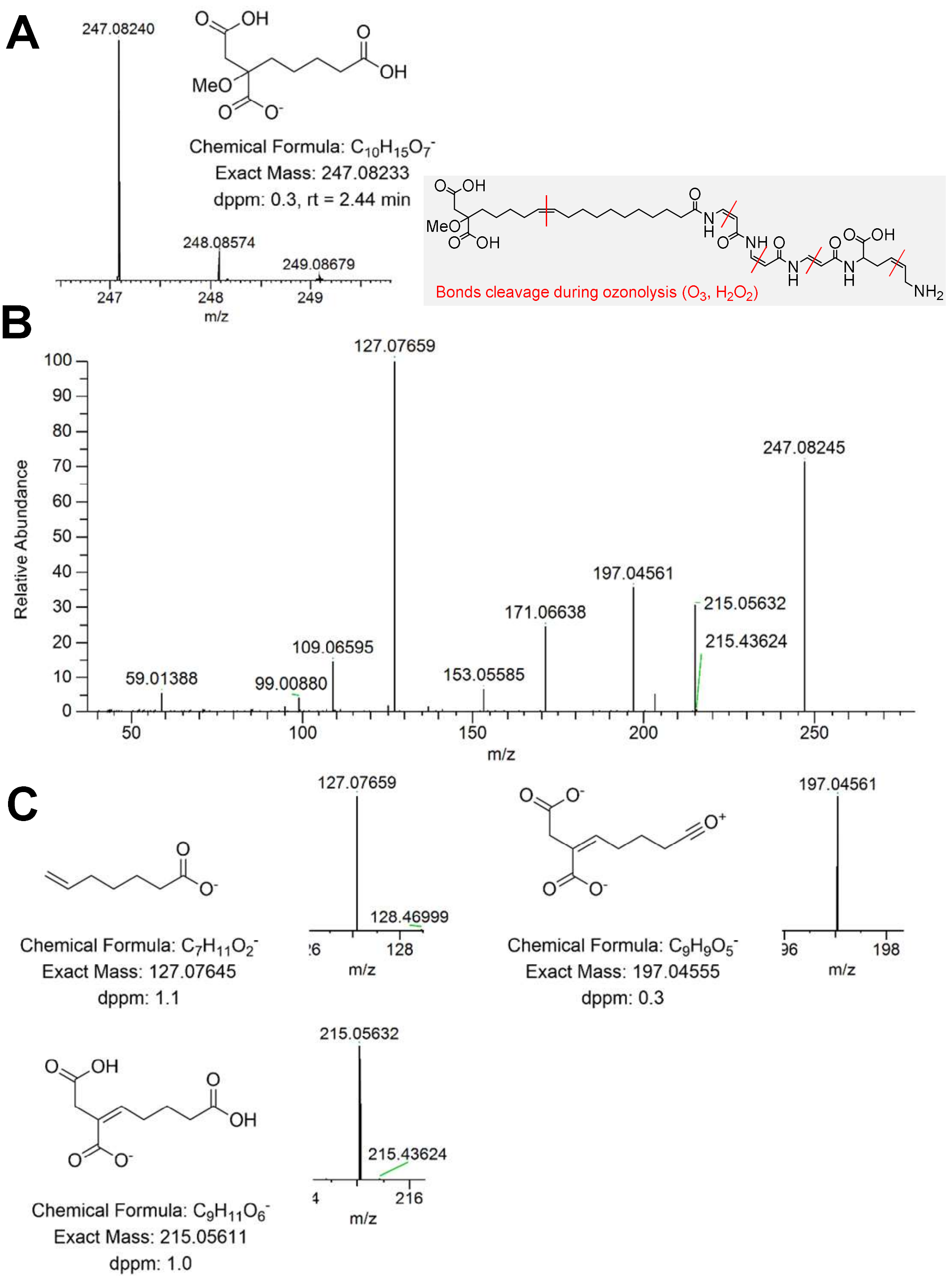


**Figure S30.** AFC-A (**1**) degradation product at 2.44 min after oxidation with ozone and oxidative work up. (**A**) HR-MS spectrum, (**B**) HR-MS/MS spectrum, and (**C**) proposed fragment structures with *m/z* extracted from HR-MS/MS spectrum. The results were obtained using Orbitrap Exploris 240 in negative mode.

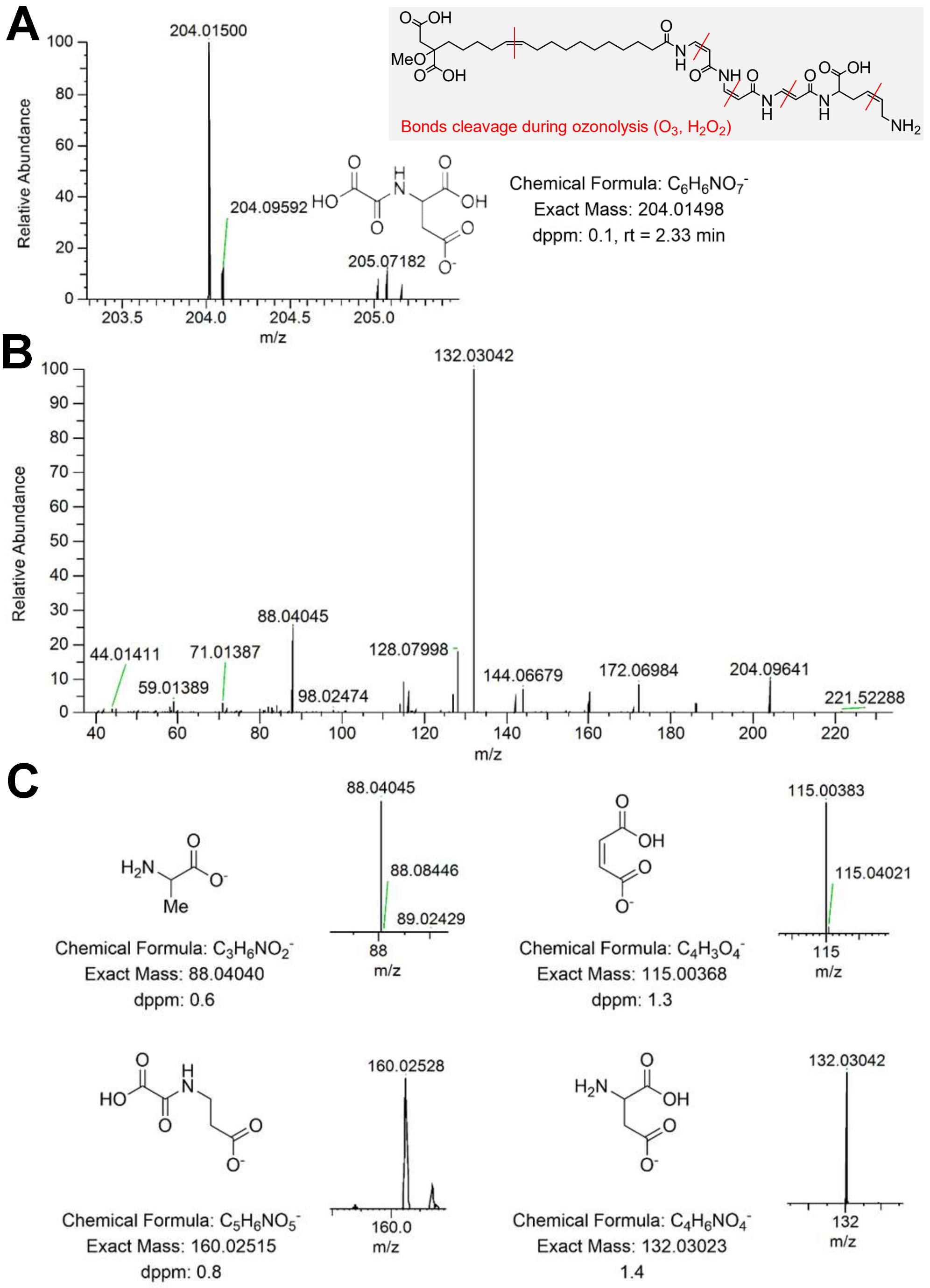


**Figure S31.** AFC-A (**1**) degradation product at 2.33 min after oxidation with ozone. (**A**) HR-MS spectrum, (**B**) HR-MS/MS spectrum, and (**C**) proposed fragment structures with *m*/*z* extracted from HR-MS/MS spectrum. The results were obtained using Orbitrap Exploris 240 in negative mode.

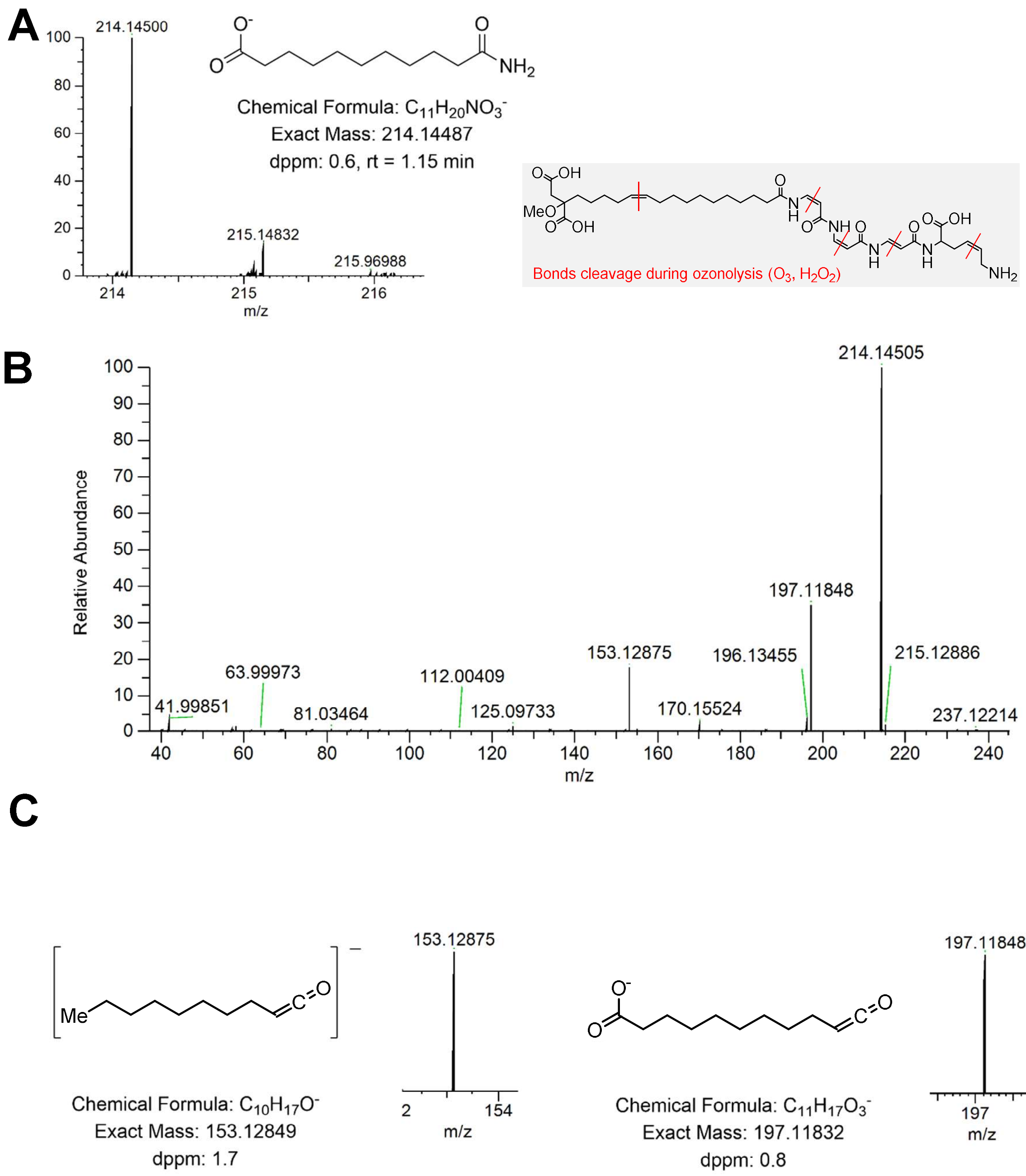


**Figure S32.** AFC-A (**1**) degradation product at 1.15 min after oxidation with ozone. (**A**) HR-MS spectrum, (**B**) HR-MS/MS spectrum, and (**C**) proposed fragment structures with *m/z* extracted from HR-MS/MS spectrum. The results were obtained using Orbitrap Exploris 240 in negative mode.

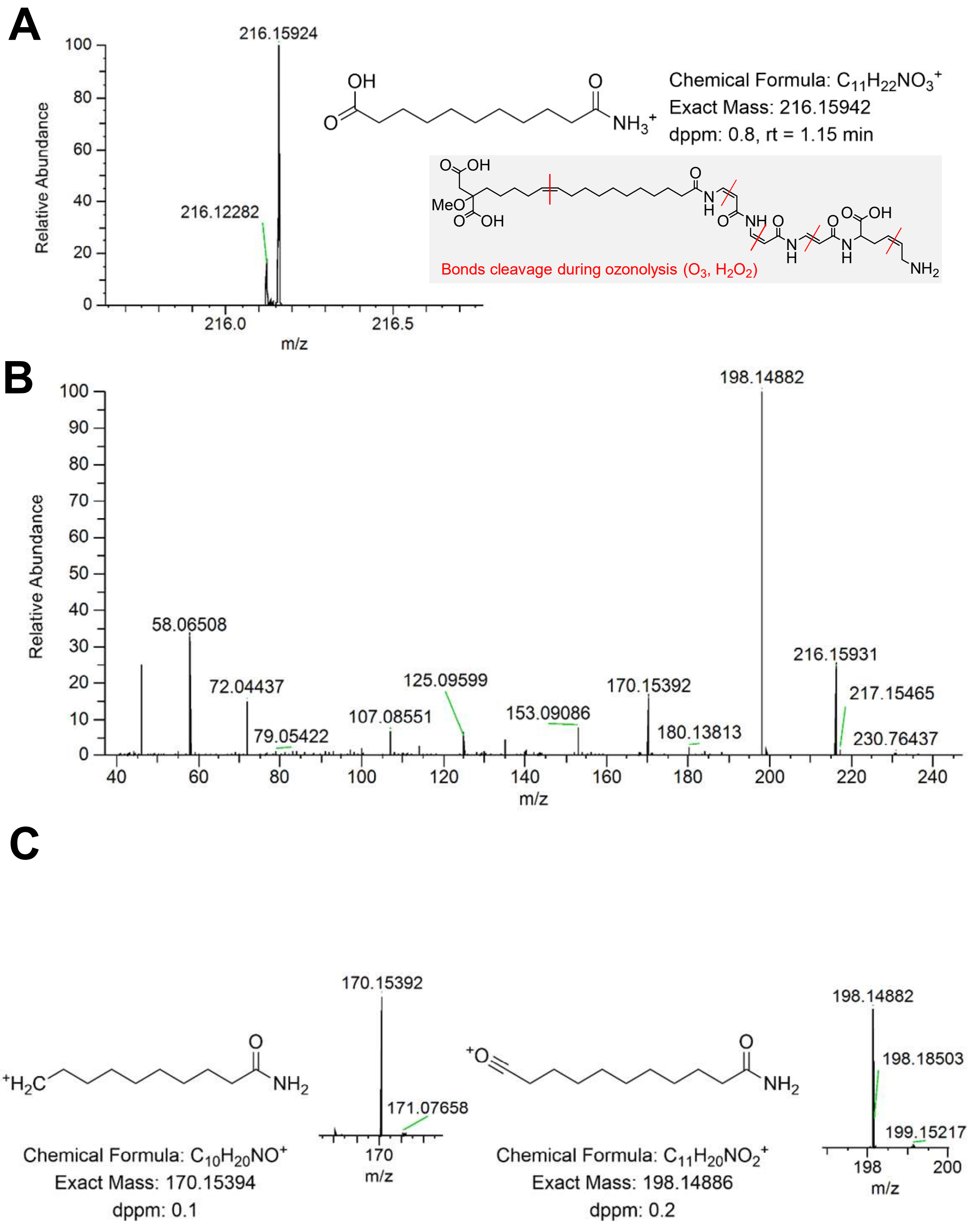


**Figure S33.** AFC-A degradation product at 1.15 min after oxidation with ozone. (**A**) HR-MS spectrum, (**B**) HR-MS/MS spectrum, and (**C**) proposed fragment structures with *m/z* extracted from HR-MS/MS spectrum. The results were obtained using Orbitrap Exploris 240 in positive mode.

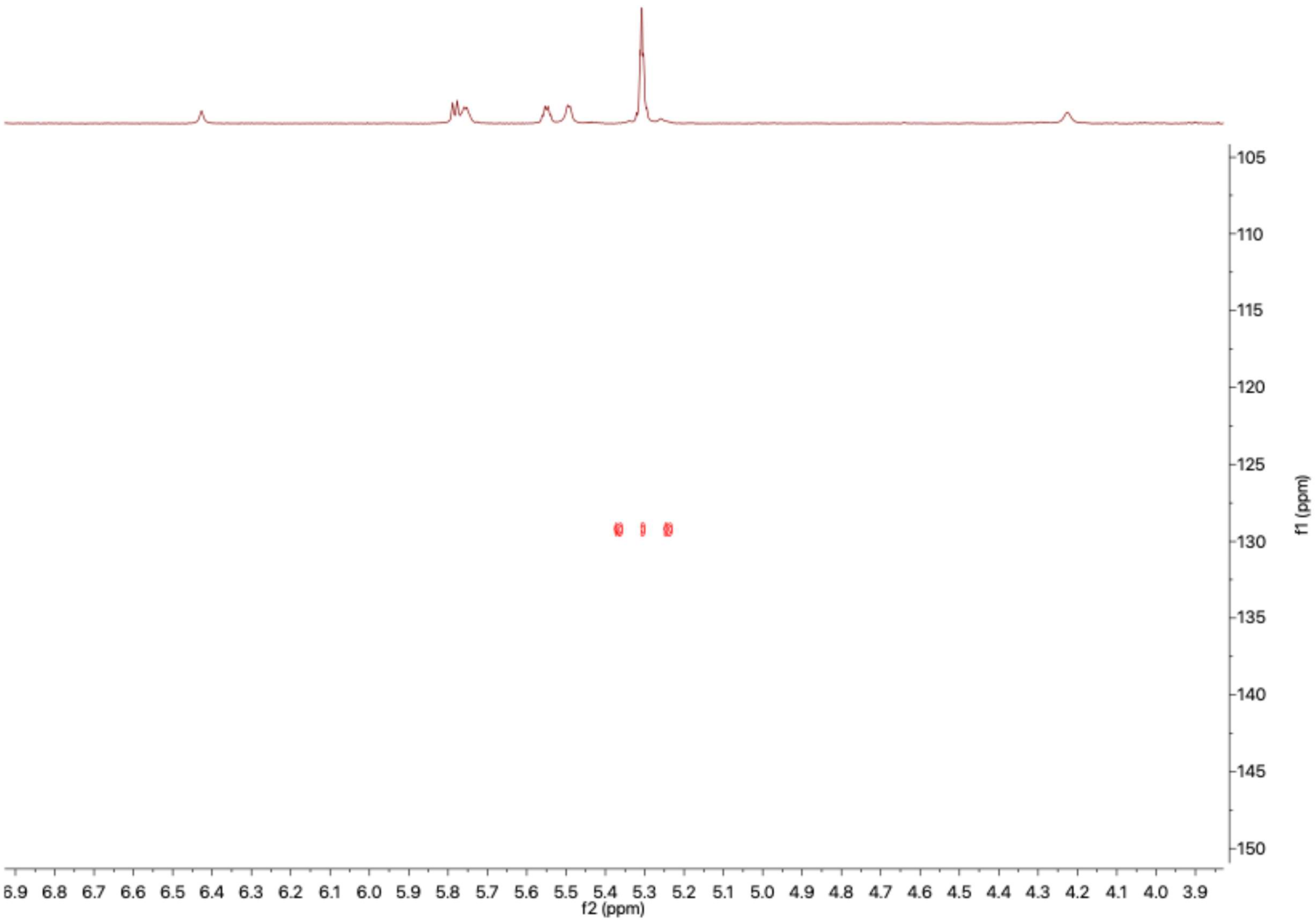


**Figure S34.** $^{1}H$-$^{13}C$-HSQC Spectrum of AFC-A (**1**) acquired without $^{13}C$ decoupling and with selective $^{1}H$ decoupling at 1.96 ppm during acquisition, (DMSO-$d_6$, 1.2 GHz, 313K).

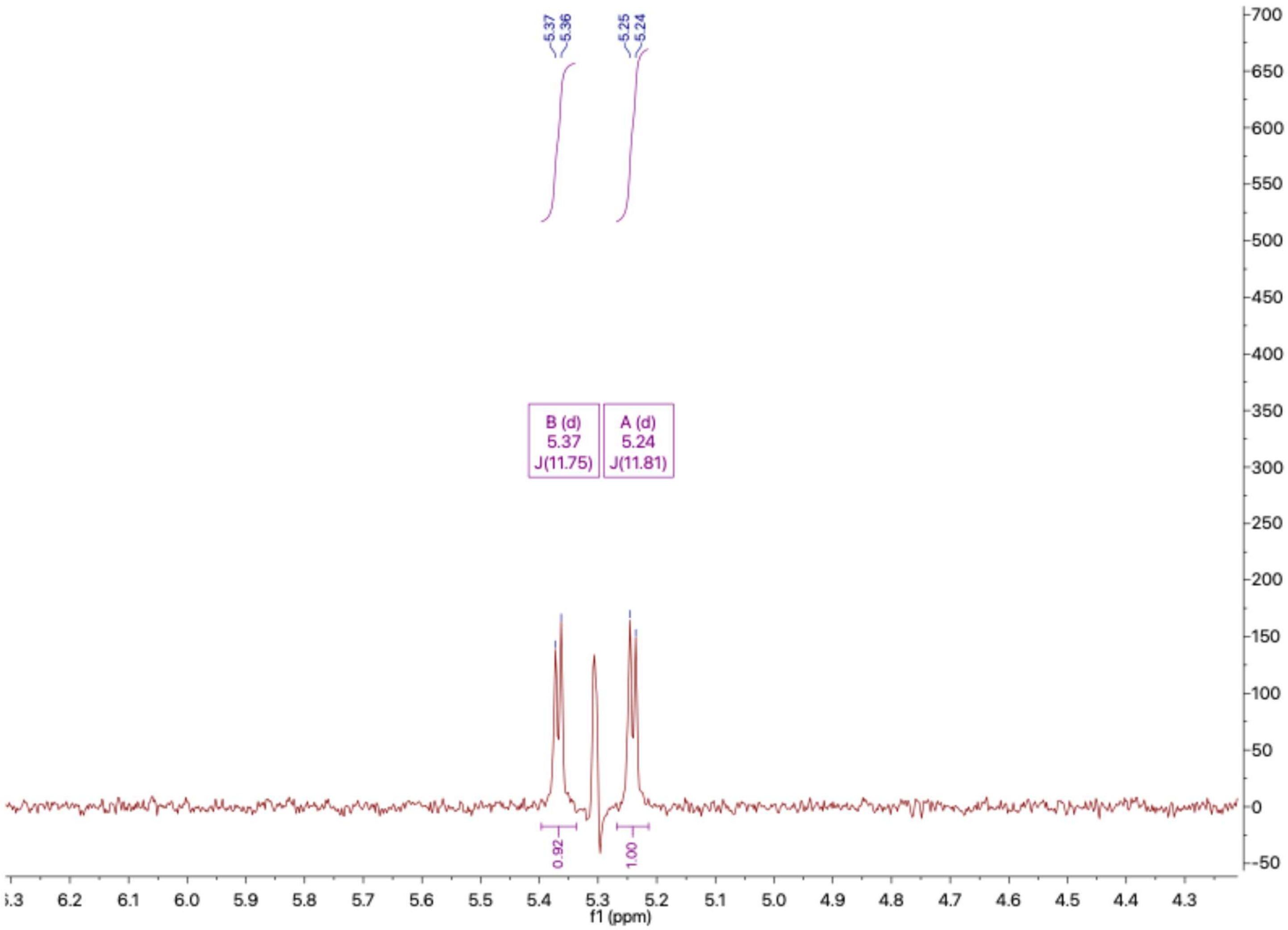


**Figure S35.** Extracted trace at 129.2 ppm from the $^1$H-$^{13}$C-HSQC spectrum of AFC-A (**1**) acquired without $^{13}$C decoupling and with selective $^1$H decoupling at 1.96 ppm during acquisition, (DMSO-$d_6$, 1.2 GHz, 313K).

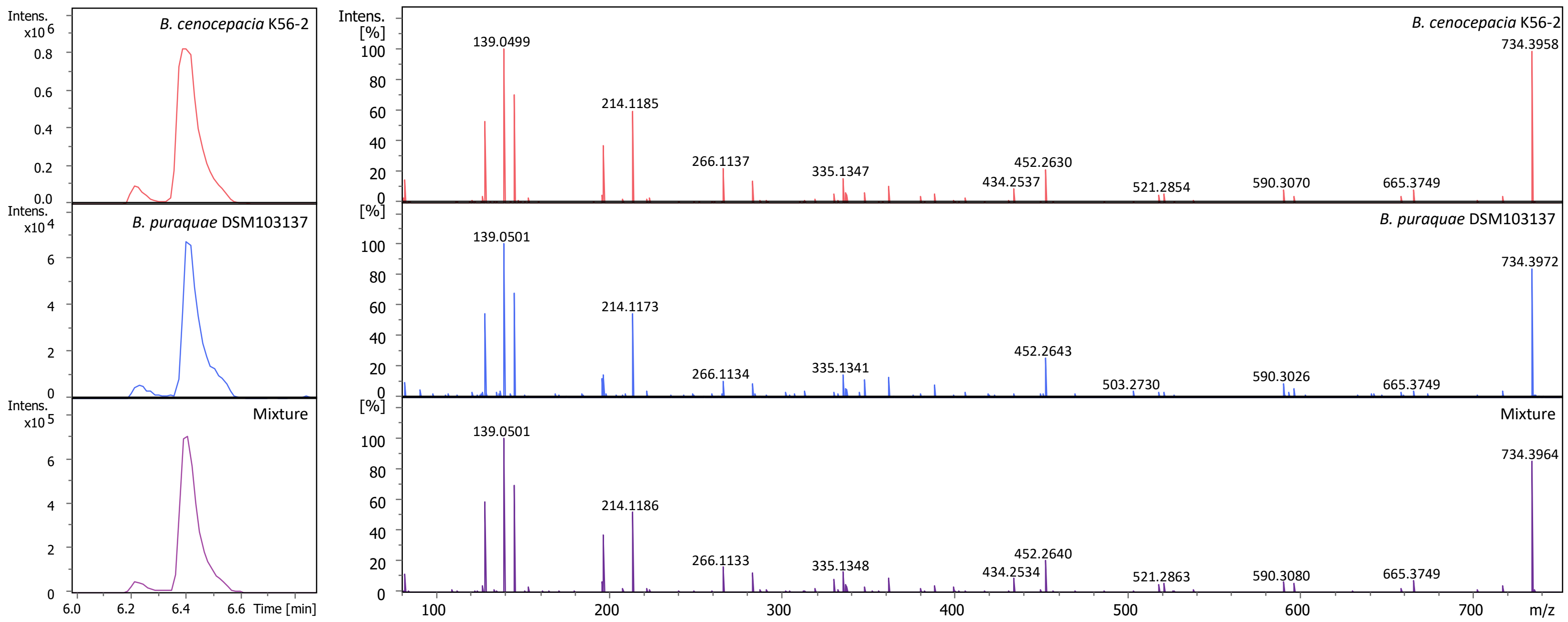


**Figure S36.** UHPLC-HR-MS/MS comparison of AFC-A from *B. cenocepacia* K56-2 and *B. puraquae* DSM103137 grown in PDB.

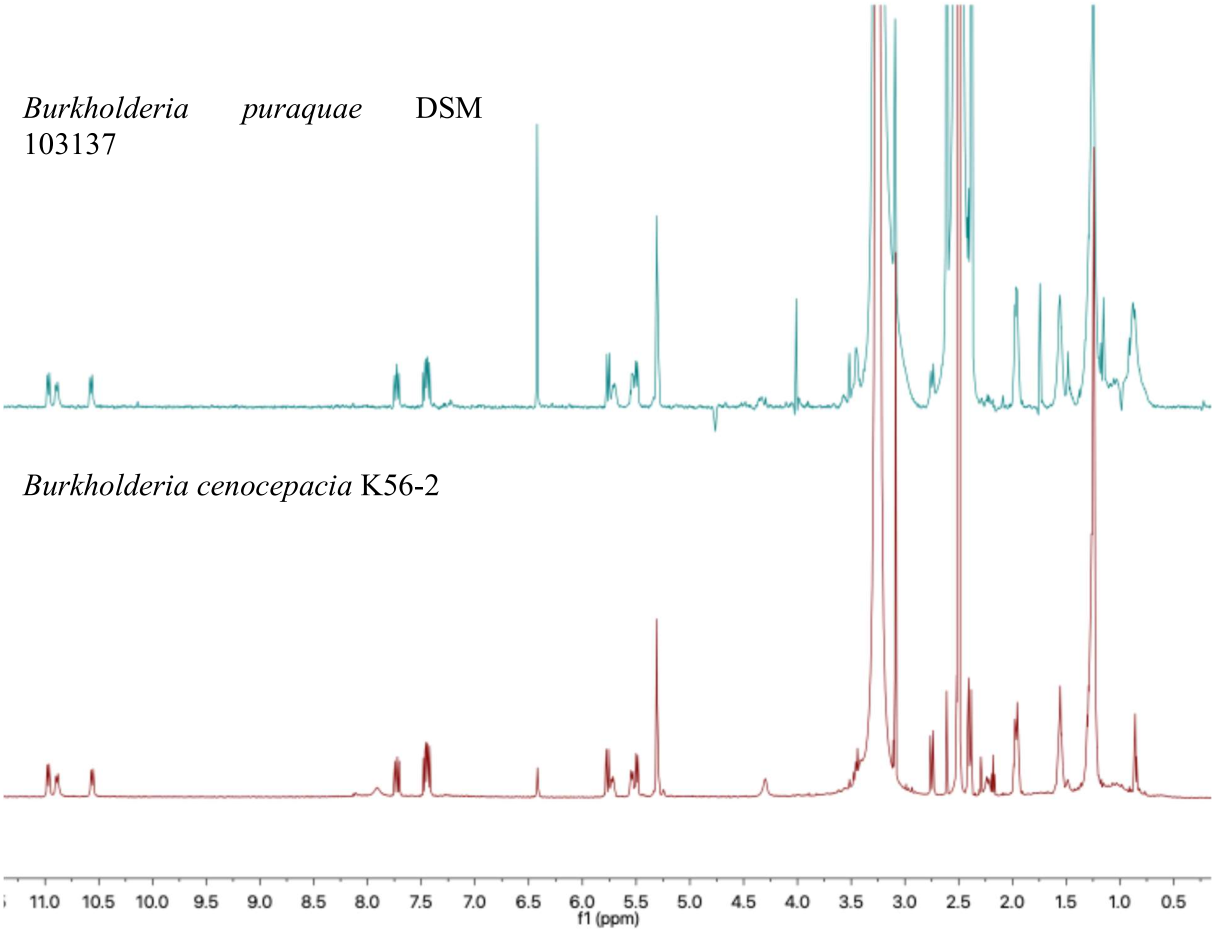


**Figure S37.** Comparison of the [1]H-NMR spectra of AFC-A extracted from *Burkholderia puraquae* DSM 103137 (top) and *Burkholderia cenocepacia* K56-2 (bottom). Spectra were acquired in DMSO-$d_6$ on a 600 MHz spectrometer at 313K.

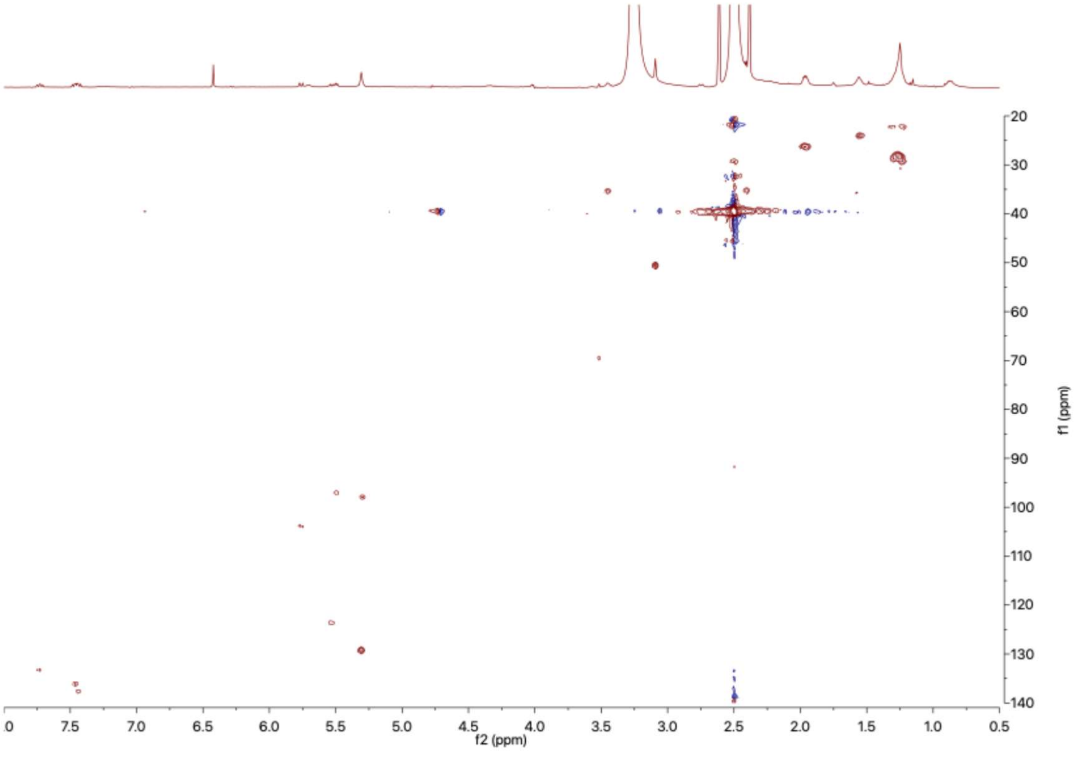


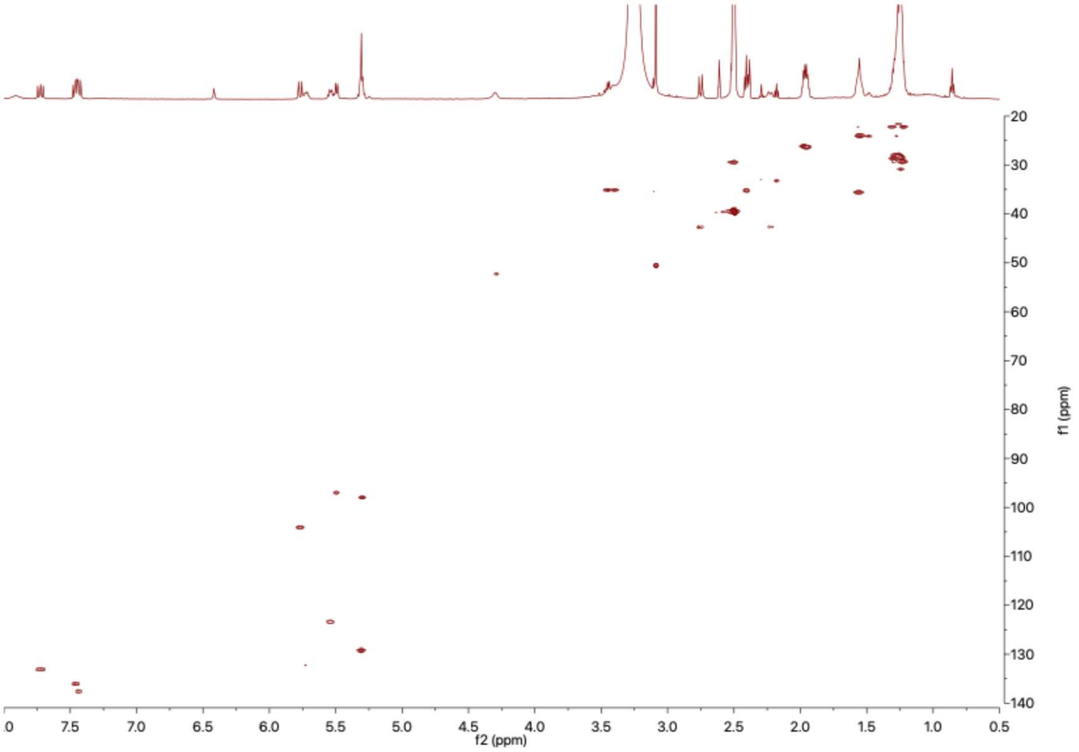


**Figure S38.** Comparison of the $^{1}H$-$^{13}C$-HSQC spectra of AFC-A extracted from *Burkholderia puraquae* DSM 103137 (top) and *Burkholderia cenocepacia* K56-2 (bottom). Spectra were acquired in DMSO-$d_6$ on a 600 MHz spectrometer at 313K.

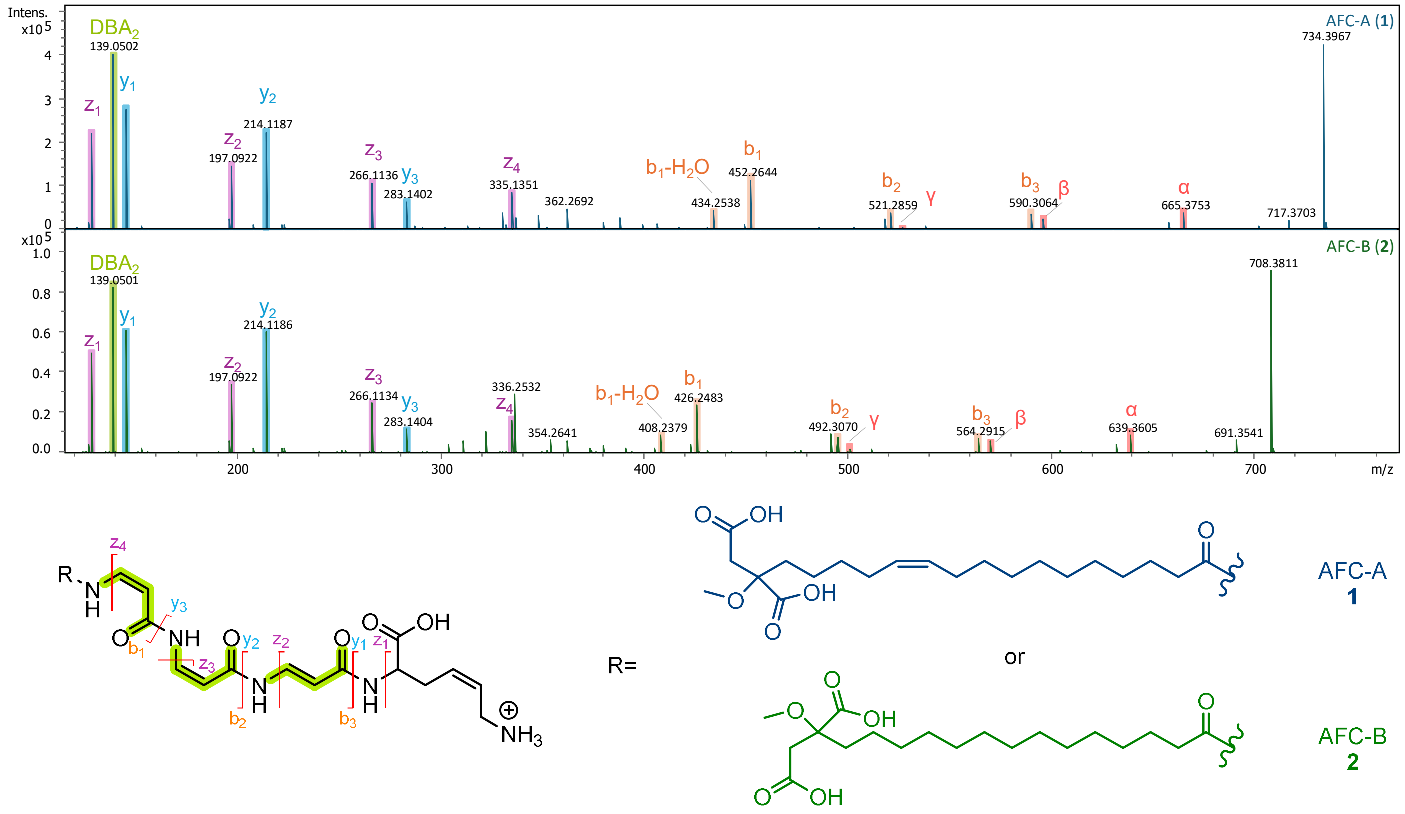


**Figure S39.** HR-MS/MS fragmentation pattern of MMFA-AFCs AFC-A (**1**) and AFC-B (**2**).

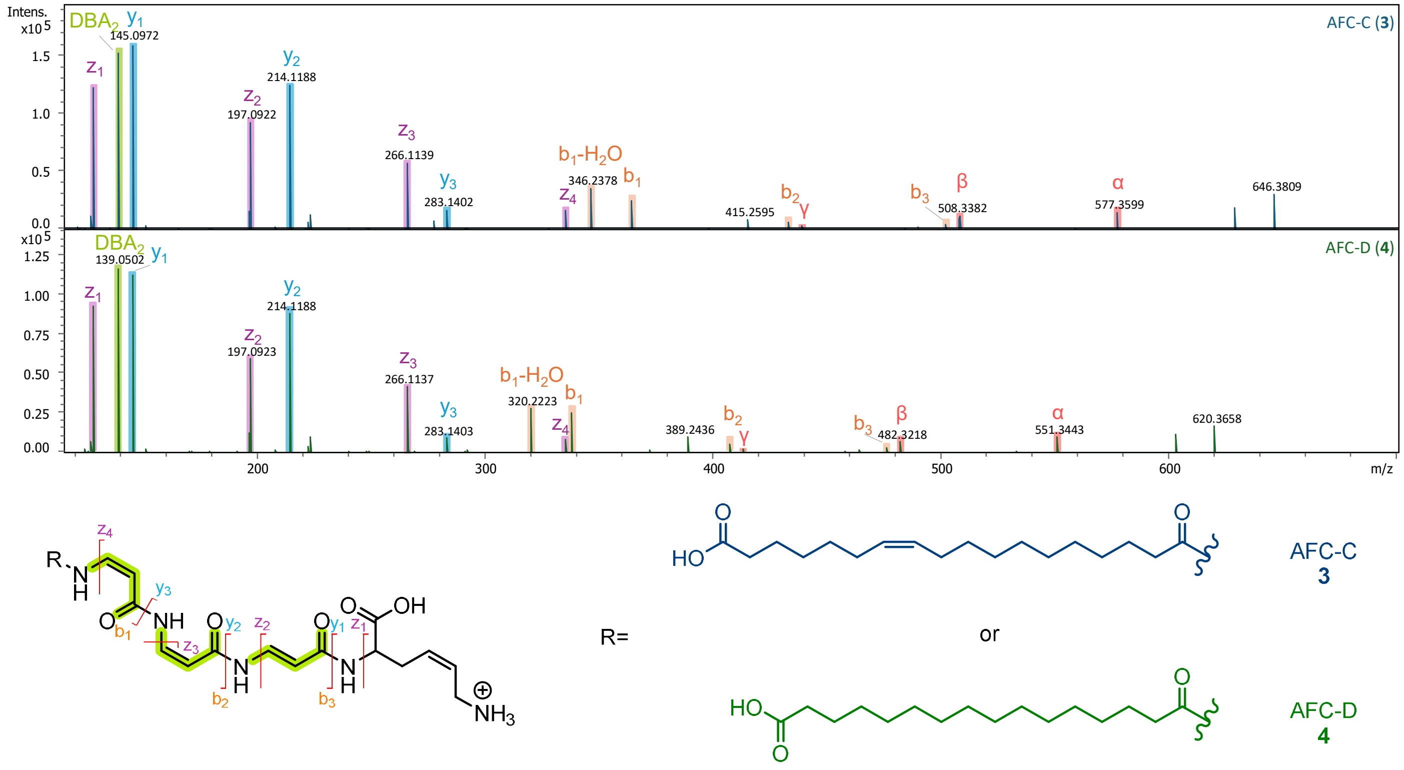


**Figure S40.** HR-MS/MS fragmentation pattern of ω-CFA-AFCs AFC-C (**3**) and AFC-D (**4**).

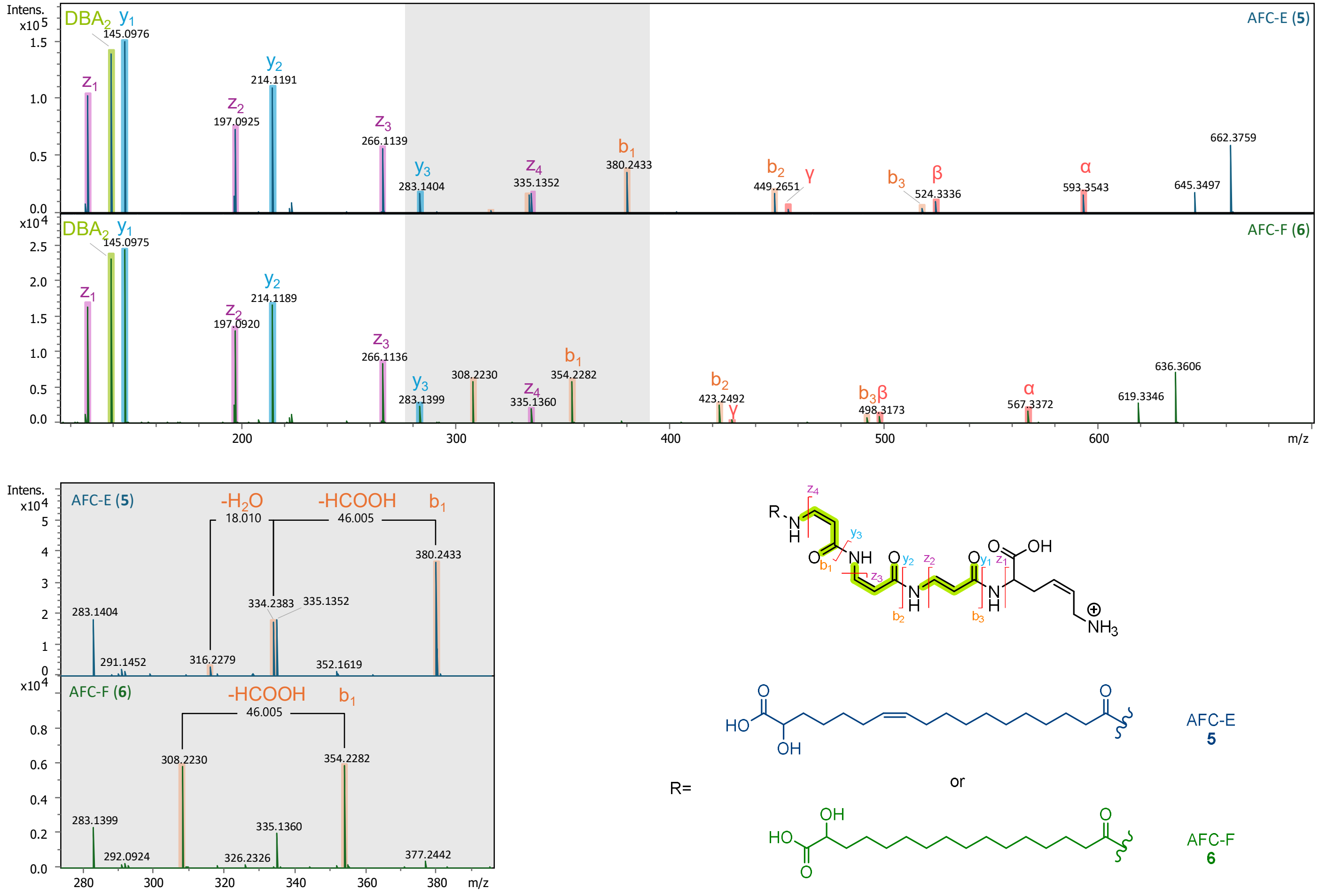

**Figure S41.** HR-MS/MS fragmentation pattern of 2-hydroxy-CFA-AFCs AFC-E (**5**) and AFC-F (**6**). The enlarged spectrum highlights the sequential loss of terminal carboxy and hydroxy groups from the $b_1$ ion (shaded in grey).

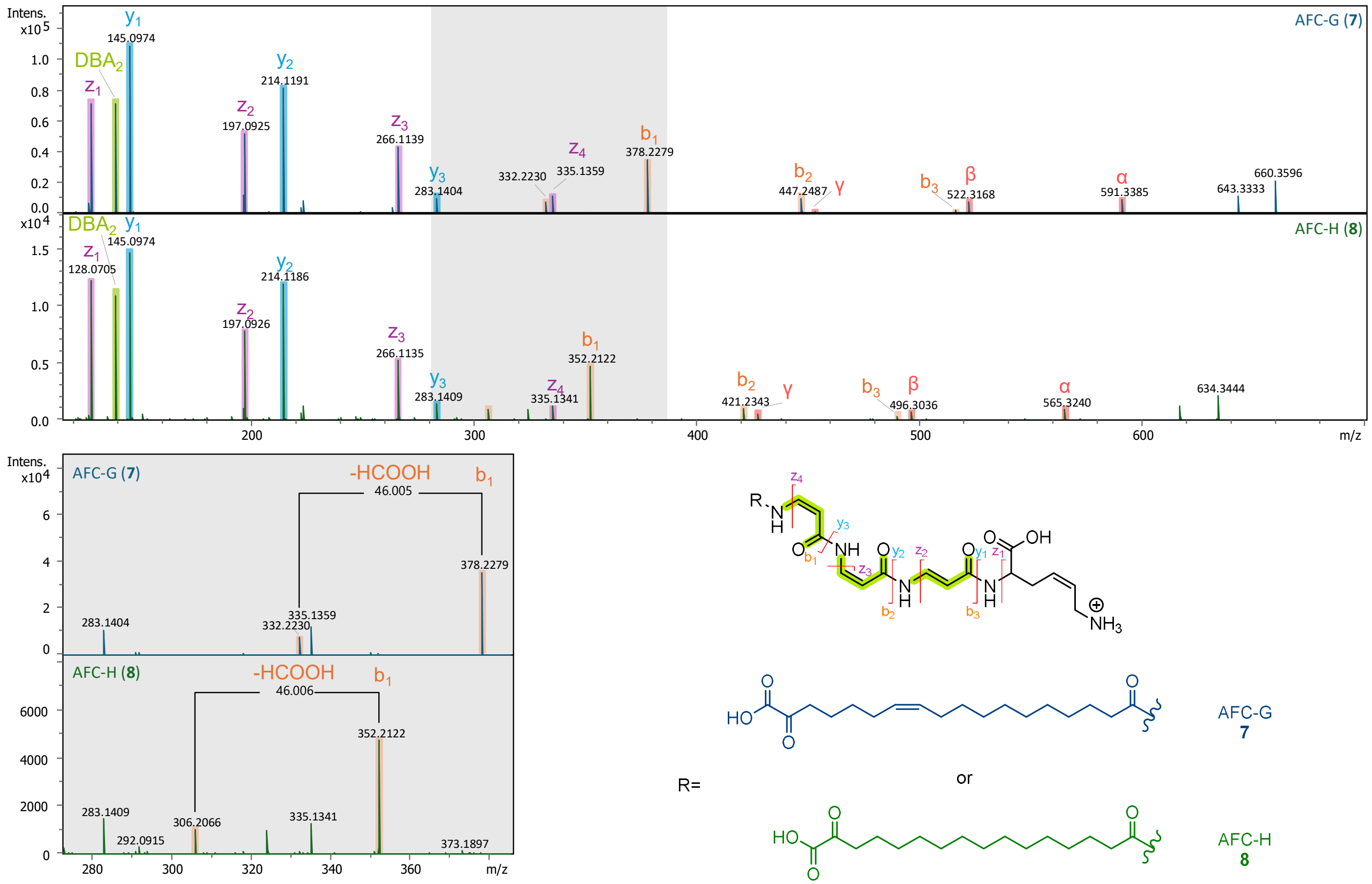


**Figure S42.** HR-MS/MS fragmentation pattern of 2-oxo-CFA-AFCs AFC-G (**7**) and AFC-H (**8**). The enlarged spectrum highlights the loss of terminal carboxy group from the $b_1$ ion (shaded in grey).

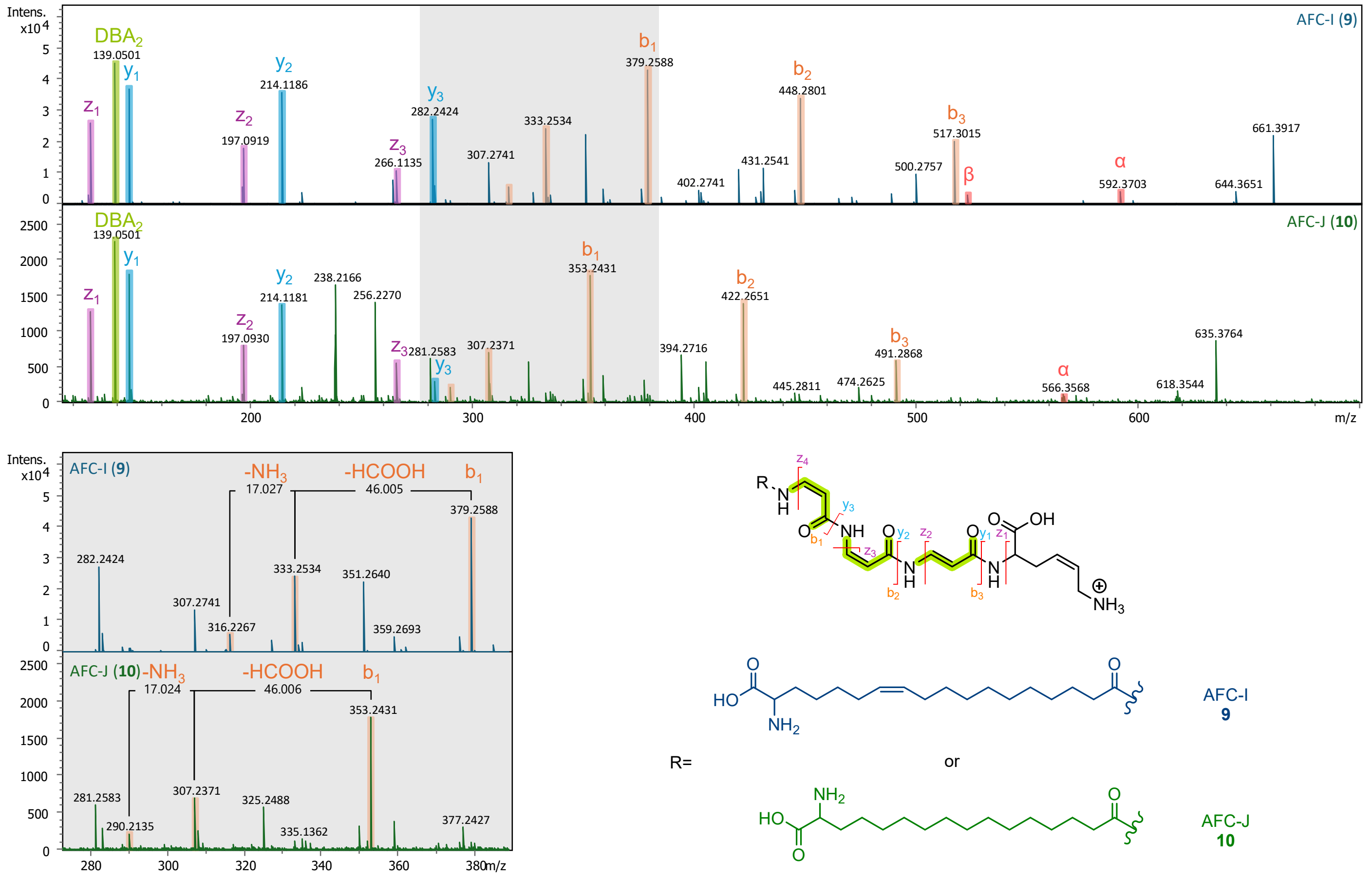

**Figure S43.** HR-MS/MS fragmentation pattern of 2-amino-CFA-AFCs AFC-I (**9**) and AFC-J (**10**). The enlarged spectrum highlights the sequential loss of terminal carboxy and amino groups from the $b_1$ ion.

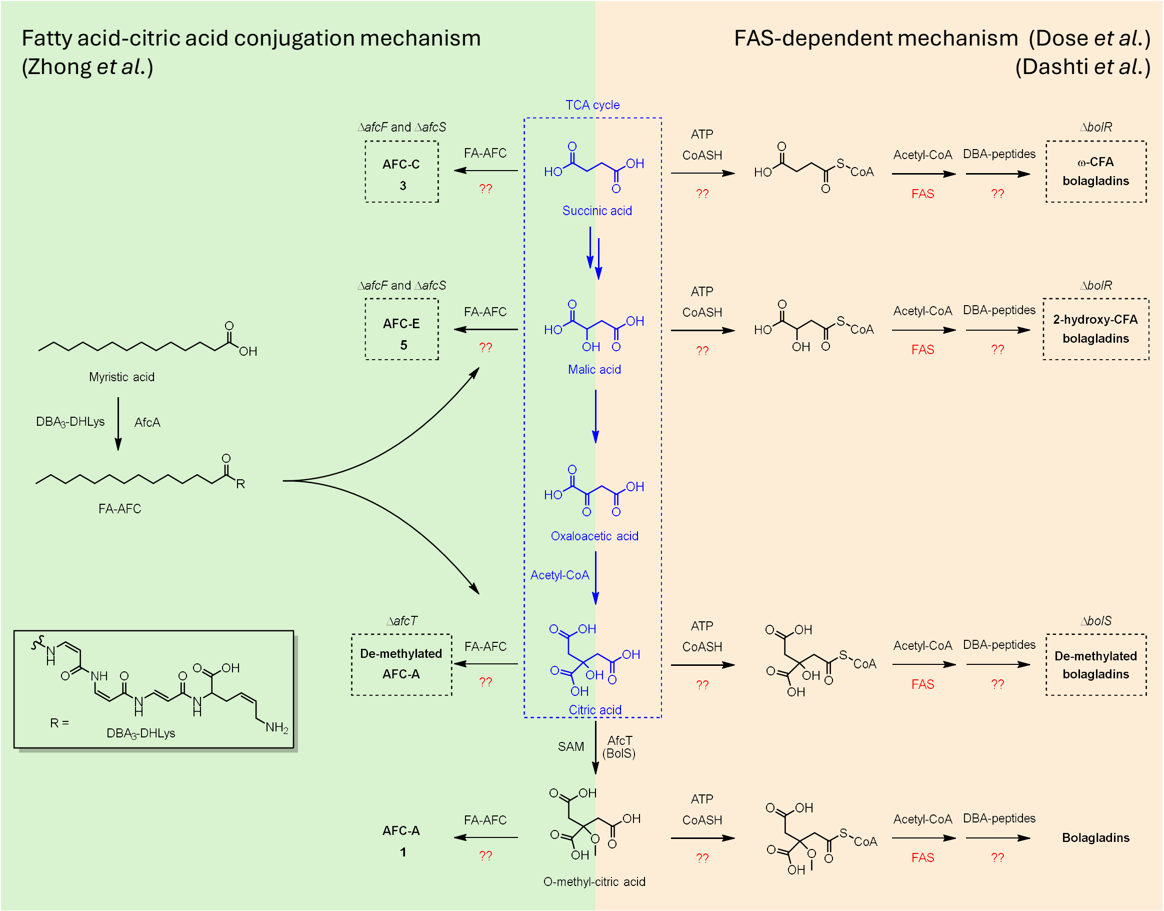


**Figure S44.** Previously proposed biosynthetic pathways of AFC and bolagladins. Based on earlier studies, AFCs biosynthesis may proceed either via fatty acid-citric acid conjugation[3] (left, green) or through a FAS-dependent pathway (right, orange) as proposed for bolagladin biosynthesis[26,28]. Reactions shared with the TCA cycle are shown in blue, and proposed reactions are highlighted in red.

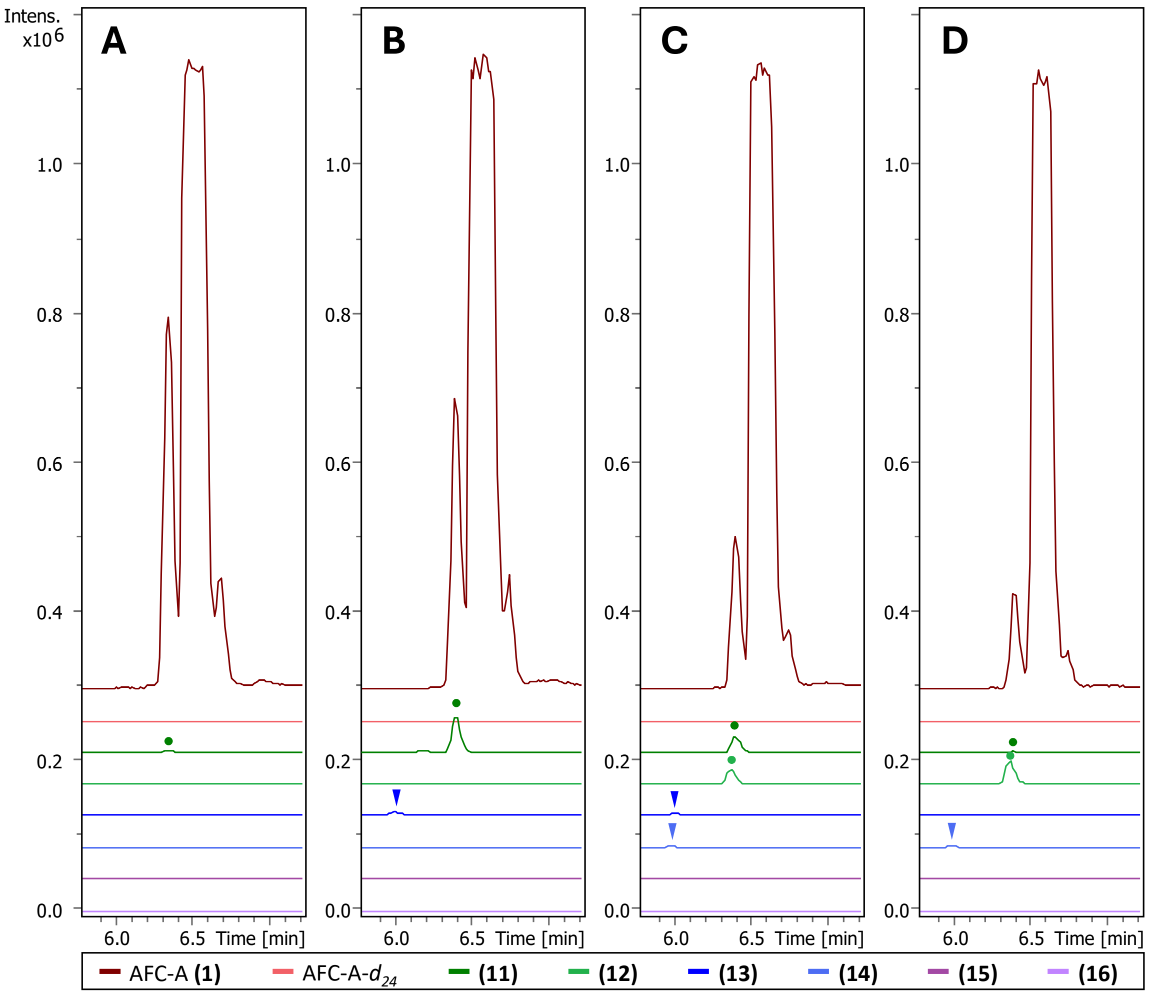


**Figure S45.** Extracted ion chromatograph of AFC production following supplementation with myristic acids. PDB cultures of wild-type K56-2 were treated with (**A**) DMSO, (**B**) myristic acid, (**C**) myristic acid:myristic acid-$d_{27}$=1:1, and (**D**) myristic acid-$d_{27}$. Peaks marked with dots indicate accumulated ω-carboxylic myristic acid-AFC intermediates (**11** and **12**; green lines), while arrows denote minor unfavored final MMFA products (**13** and **14**; blue lines). No 2-hydroxy carboxylic myristic acid-AFCs were detected (**15** and **16**; purple lines).

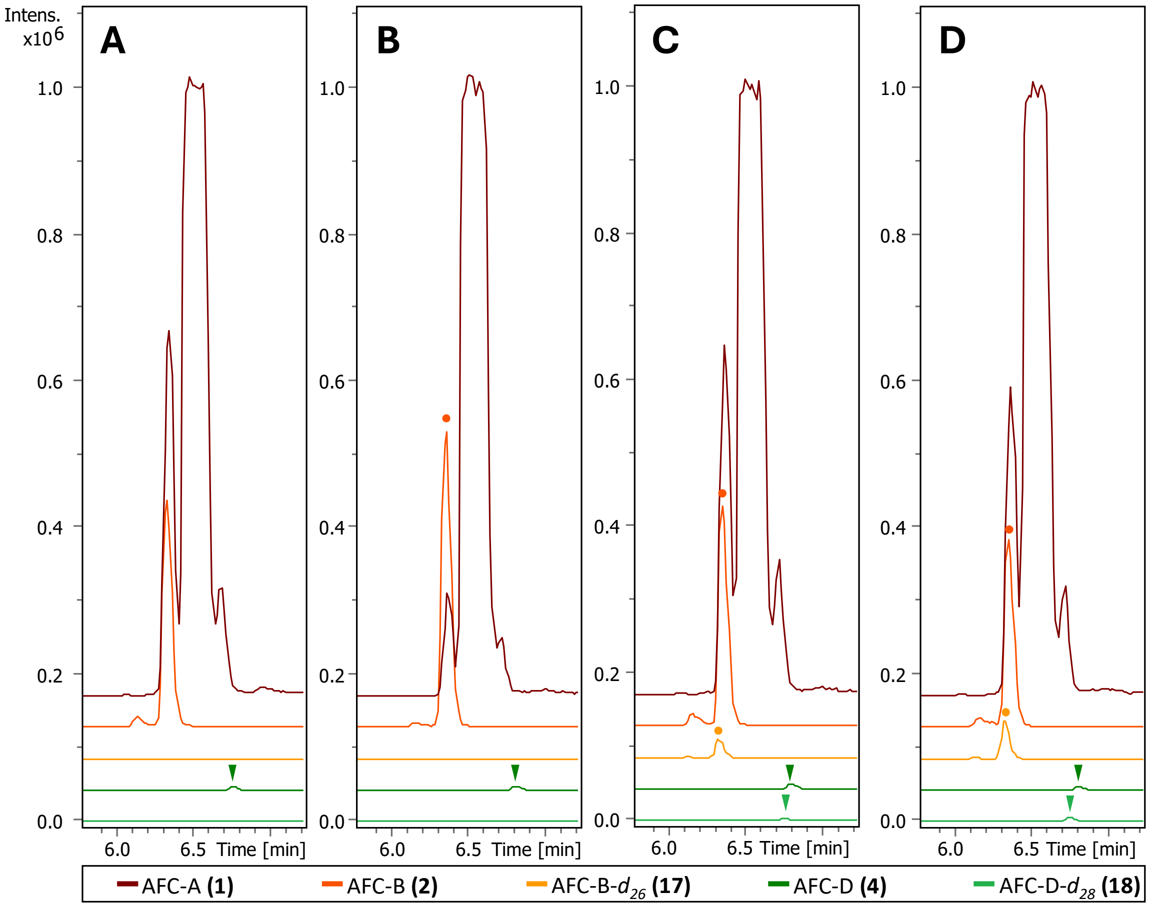


**Figure S46.** Extracted ion chromatograph of AFC production following supplementation with palmitic acids. PDB cultures of wild-type K56-2 were treated with (**A**) DMSO, (**B**) palmitic acid, (**C**) palmitic acid:palmitic acid-$d_{31}$=1:1, and (**D**) palmitic acid-$d_{31}$. Peaks marked with dots indicate *O*-methylated malic acid-palmitic acid-AFCs (**2** and **17**; orange lines), while arrows denote minor ω-carboxylic palmitic acid-AFC intermediates (**4** and **18**; green lines).

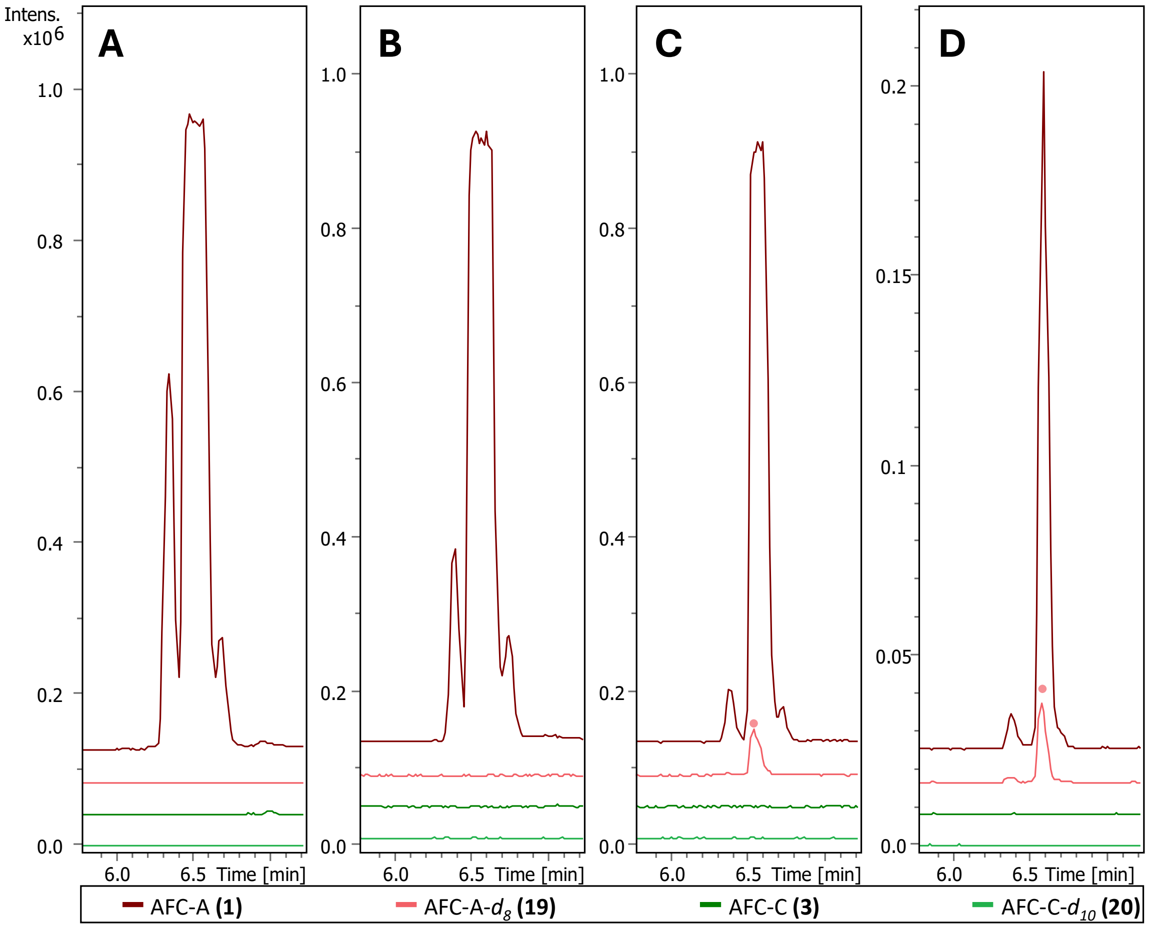


**Figure S47.** Extracted ion chromatograph of AFC production following supplementation with *cis*-vaccenic acids. PDB cultures of wild-type K56-2 were treated with **(A)** DMSO, **(B)** *cis*-vaccenic acid, **(C)** *cis*-vaccenic acid: *cis*-vaccenic acid-$d_{13}$=1:1, and **(D)** *cis*-vaccenic acid-$d_{13}$. Peaks denoted with dots indicate deuterium-labeled AFC-A-$d_8$ (**19**; pink), no deuterium-labeled AFC intermediate AFC-C-$d_{10}$ (**20**; light green) was detected.

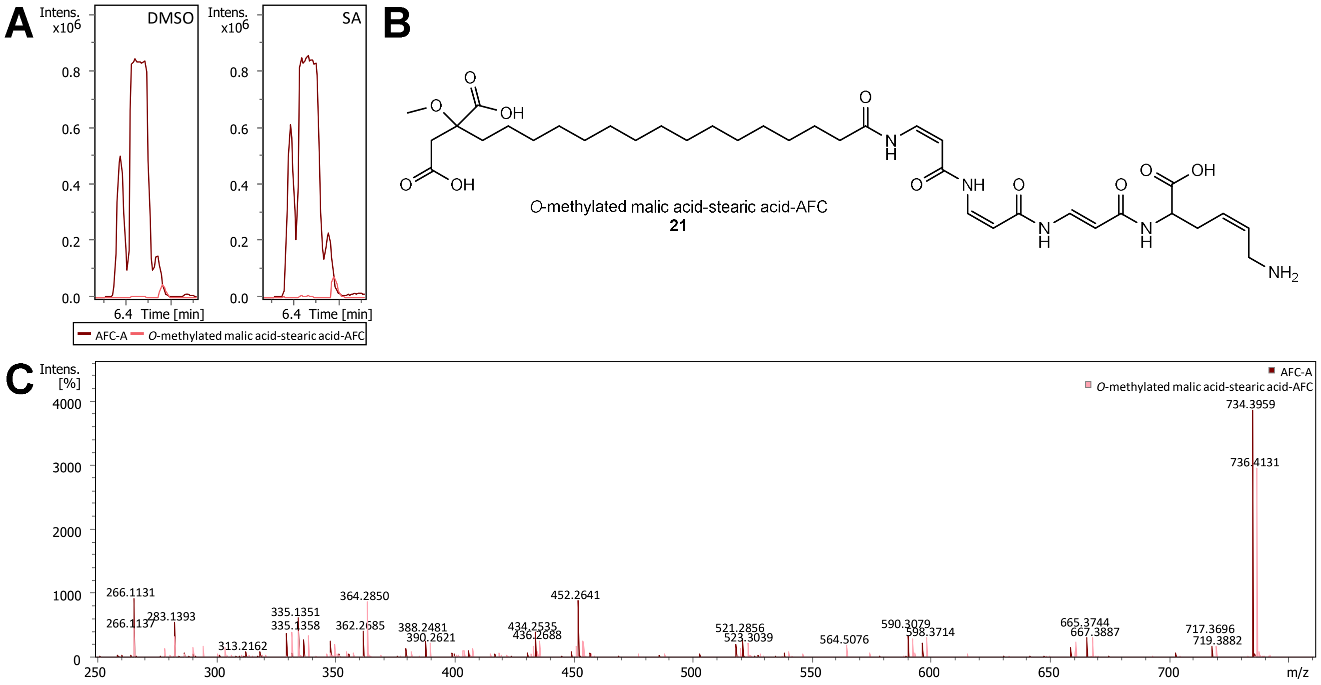


**Figure S48.** AFC production following supplementation with stearic acid in PDB. (**A**) Extracted ion chromatograph of *O*-methylated malic acid-stearic acid-AFC (**21**) in DMSO control and stearic acid-supplemented cultures. (**B**) Structure of the *O*-methylated malic acid-stearic acid-AFC (**21**). (**C**) HR-MS/MS fragmentation comparison between AFC-A and *O*-methylated malic acid-stearic acid-AFC (**21**), showing a +2H shift in the acyl side-chain fragments.

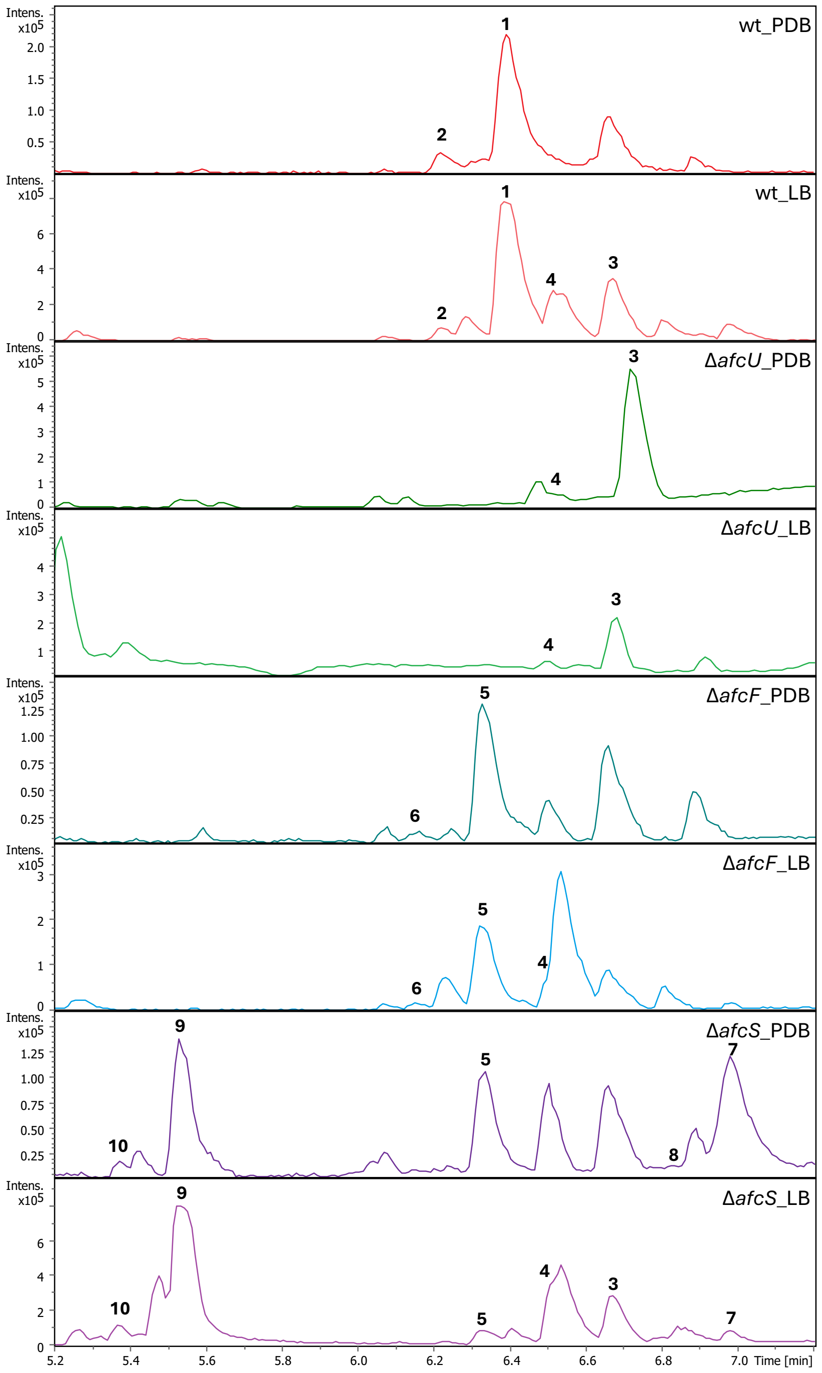


**Figure S49.** Base peak chromatograms of AFCs produced by wild-type, Δ*afcU*, Δ*afcF* and Δ*afcS* strains grown in PDB and LB medium. AFC-A to J is indicated by compound numbers **1** to **10**.

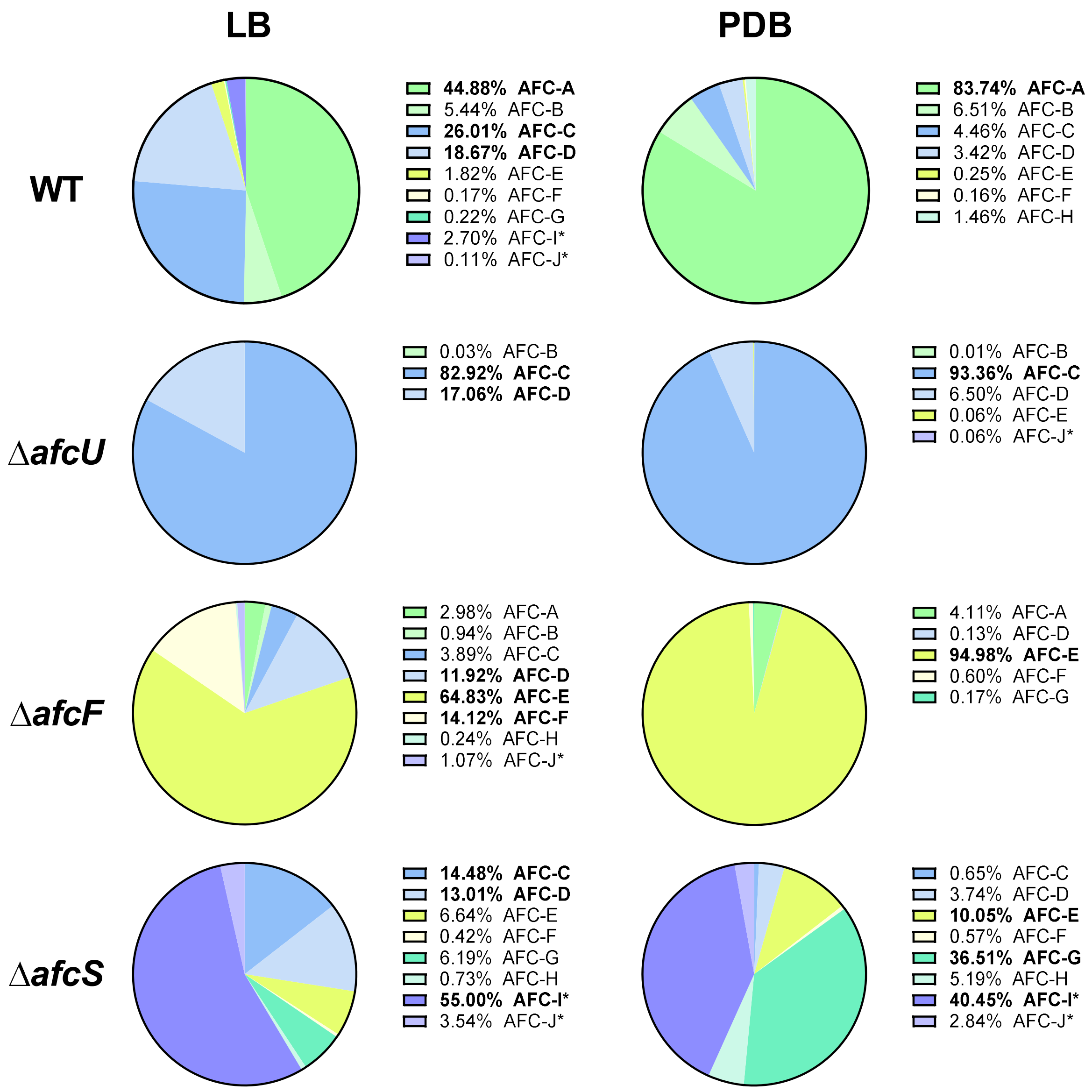


**Figure S50.** Ion abundances of AFCs produced by wildtype, Δ*afcU*, Δ*afcF* and Δ*afcS* strains grown in LB and PDB media. Peak areas of AFC ions (**1** to **10**) with $1^+$ and $2^+$ charges were integrated. Compounds with relative amounts greater than 10% are shown in bold. *Note: AFC-I and AFC-J predominantly form $2^+$ ions, so their absolute quantities may differ from those of the other AFCs.

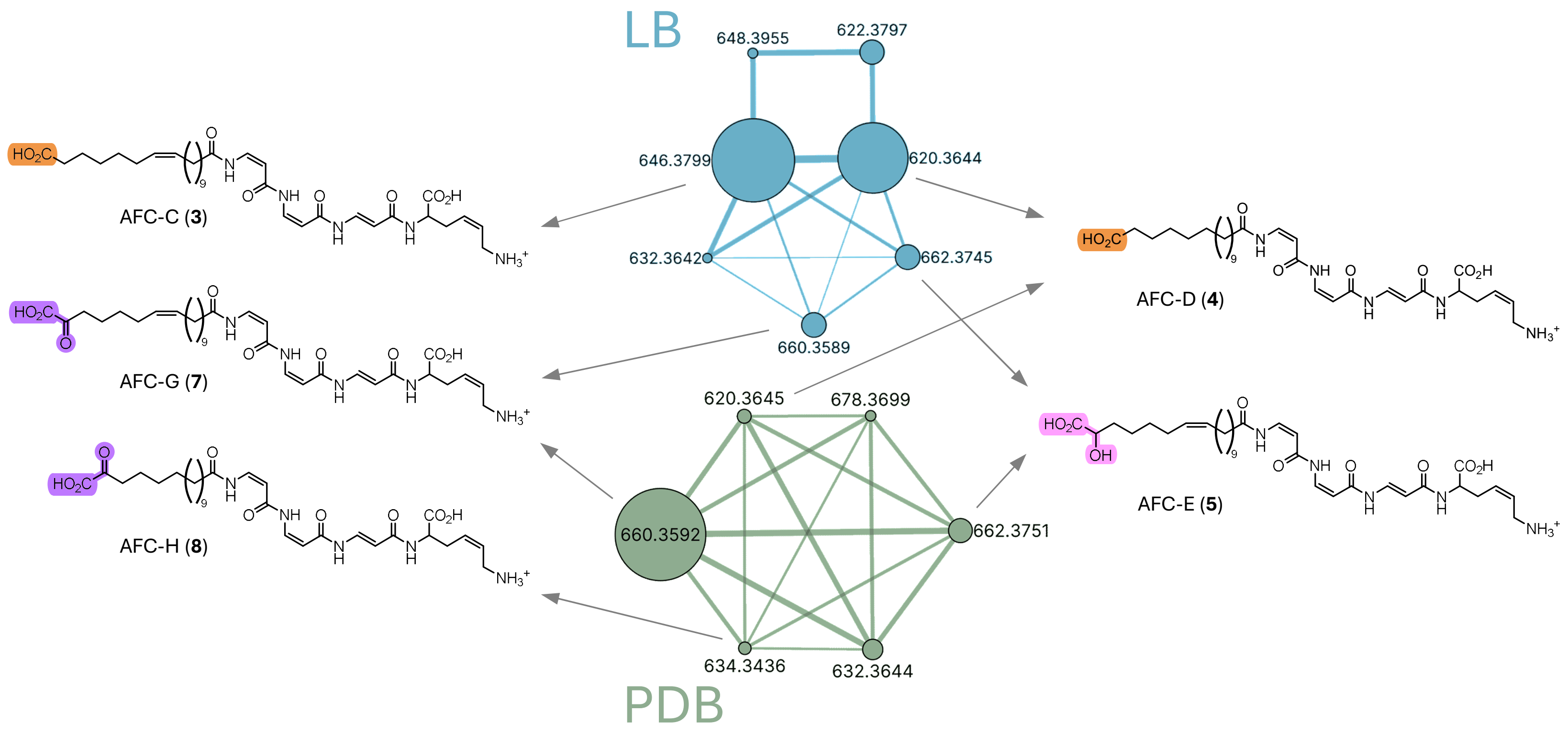


**Figure S51.** Molecular networking analysis of the Δ*afcF* mutant cultured in LB and PDB media, highlighting the network of AFC and its derivatives. Node size corresponds to the peak area of each compound. Proposed structures of main derivatives are illustrated, with the functional groups on the aliphatic side chain (carboxylic acid, 2-hydroxy carboxylic acid and 2-oxo carboxylic acid) highlighted in orange, pink and purple, respectively.

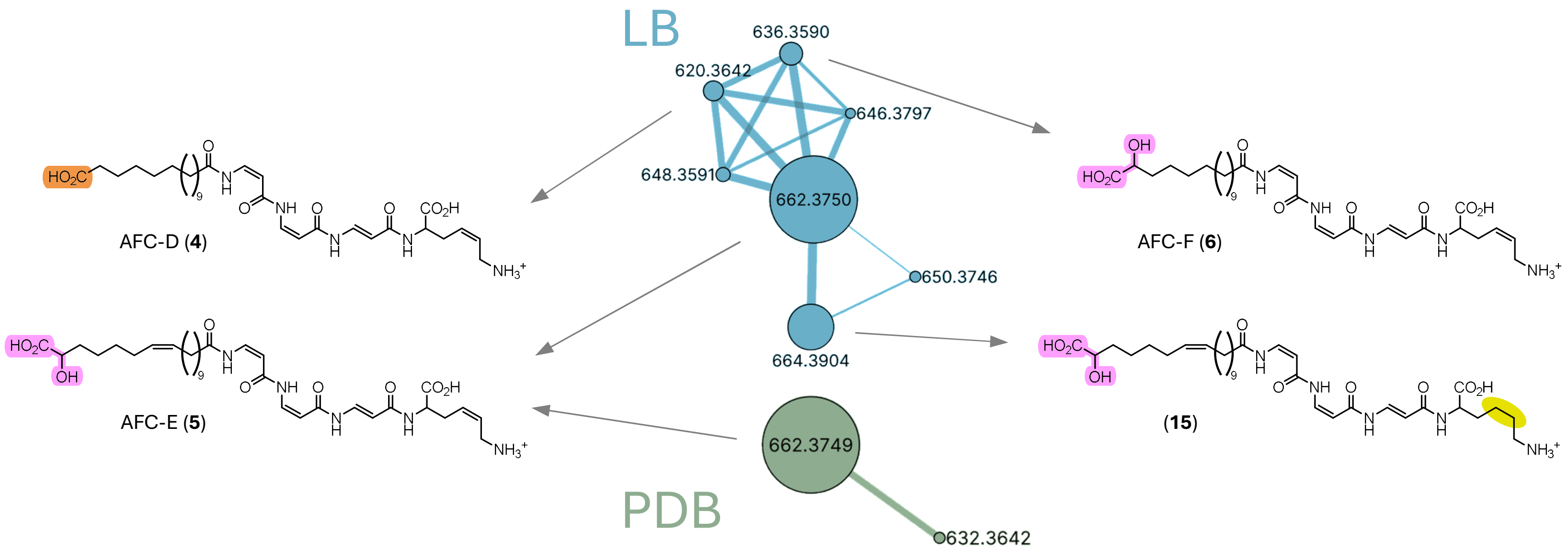


**Figure S52.** Molecular networking analysis results of the Δ*afcS* mutant cultured in LB and PDB medium, highlighting the network of AFC and its derivatives. Node size corresponds to the peak area of each compound. Proposed structures of the main derivatives are illustrated. Functional groups on the aliphatic side chain (carboxylic acid and 2-hydroxy-carboxylic acid) are highlighted in orange and pink, respectively. The non-reduced bond on lysine at the C-terminus is highlighted in yellow.

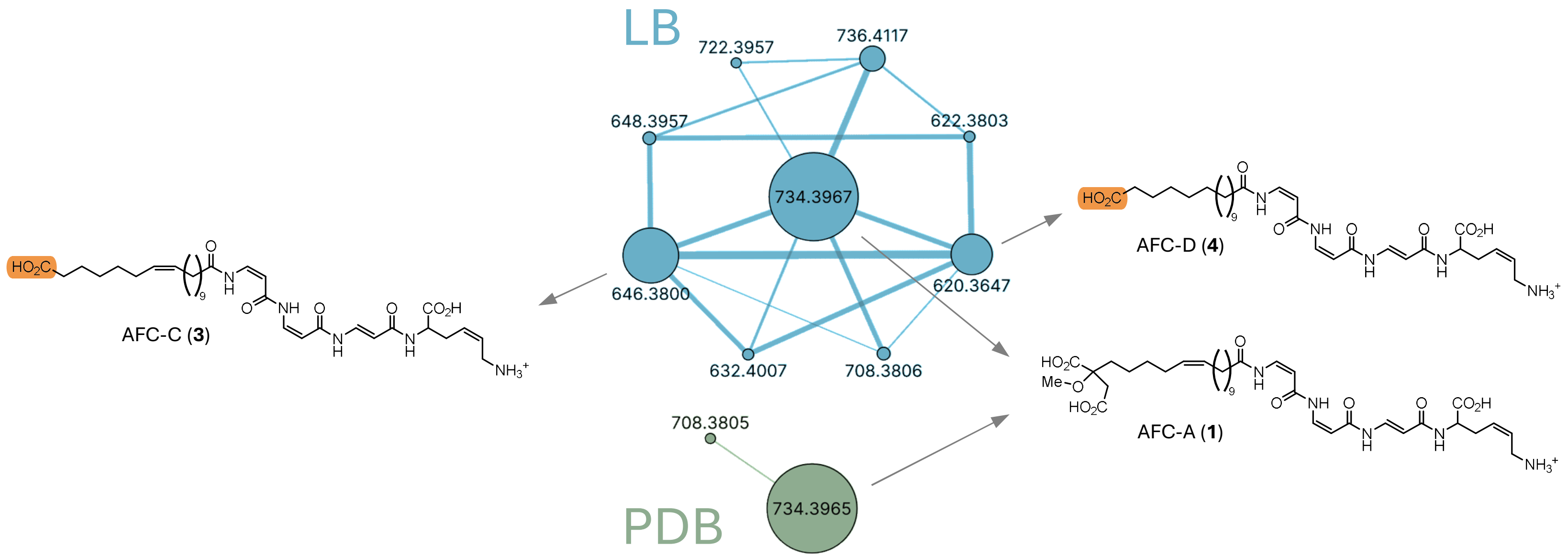


**Figure S53.** Molecular networking analysis results of the wildtype sample cultured in LB and PDB medium, highlighting the network of AFC and its derivatives. Node size corresponds to the peak area of each compound. Proposed structures of the main derivatives are illustrated. The carboxylic acid functional group on the aliphatic side chain is highlighted in orange.

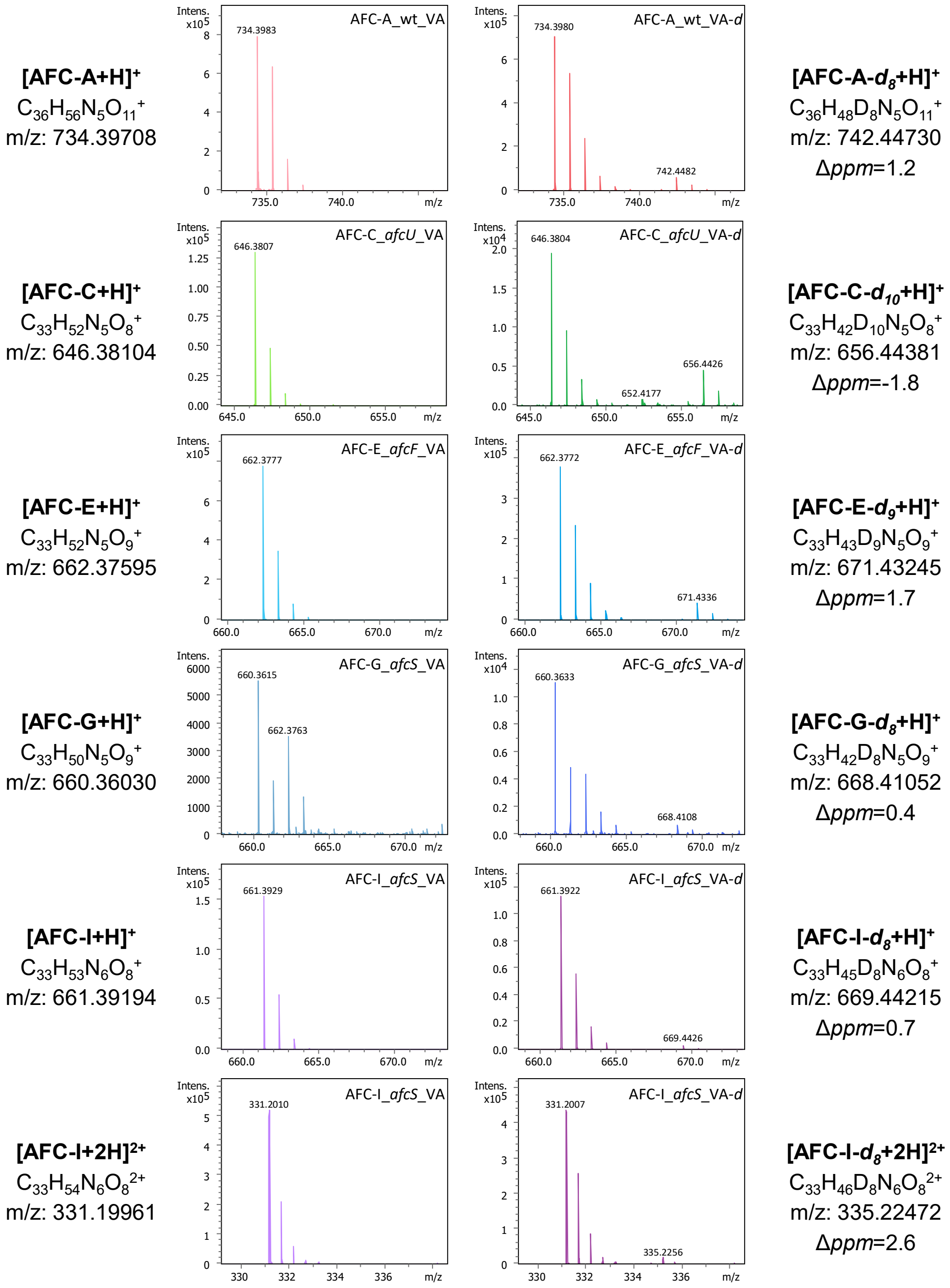


**Figure S54.** MS identification of deuterium-labeled AFCs following supplementation with *cis*-vaccenic acid-$d_{13}$. AFC compounds were detected in cultures of wild-type, Δ*afcU*, Δ*afcF* and Δ*afcS* strains grown in PDB medium with either *cis*-vaccenic acid (VA) or *cis*-vaccenic acid-$d_{13}$ (VA-*d*).

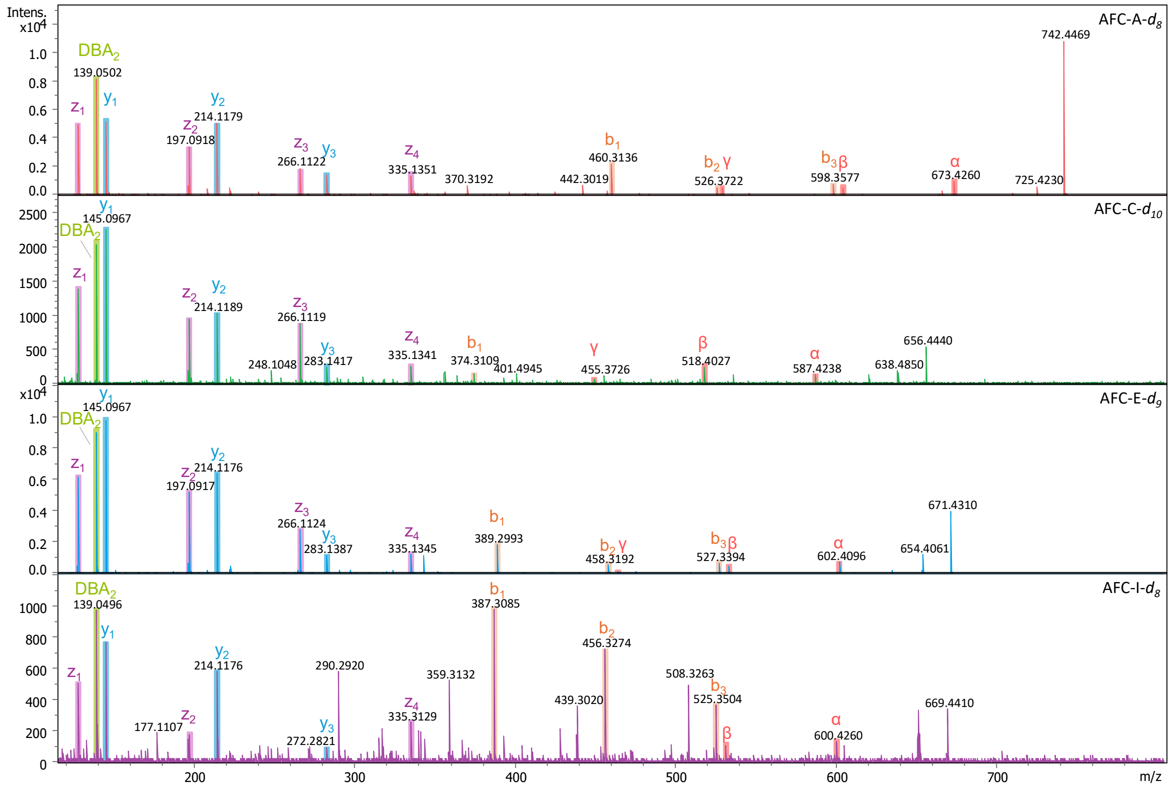


**Figure S55.** HR-MS/MS fragmentation of deuterium-labeled AFCs. Deuterium labeled AFC compounds were identified based on HR-MS/MS fragmentation patterns. Fragmentation data for AFC-G-$d_8$ are not shown due to low signal intensity.

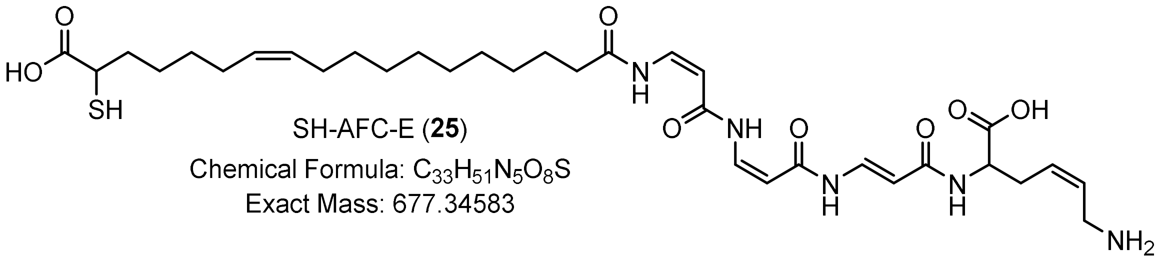

SH-AFC-E (**25**)
Chemical Formula: $C_{33}H_{51}N_5O_8S$
Exact Mass: 677.34583

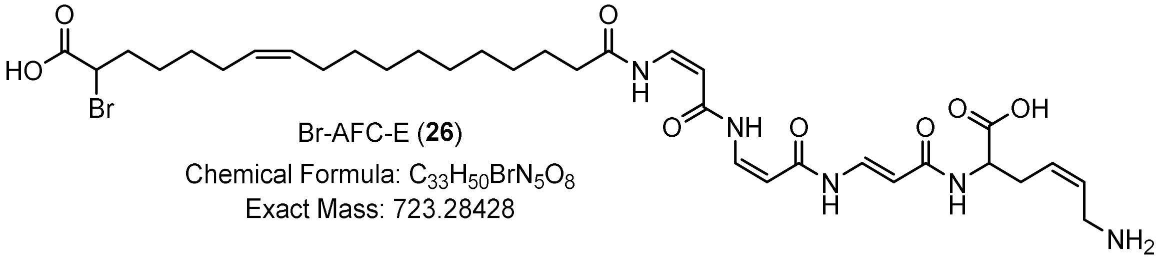

Br-AFC-E (**26**)
Chemical Formula: $C_{33}H_{50}BrN_5O_8$
Exact Mass: 723.28428

**Figure S56.** Proposed SH- and Br-substituted AFC products formed upon supplementation with mercaptosuccinic acid and bromosuccinic acid.

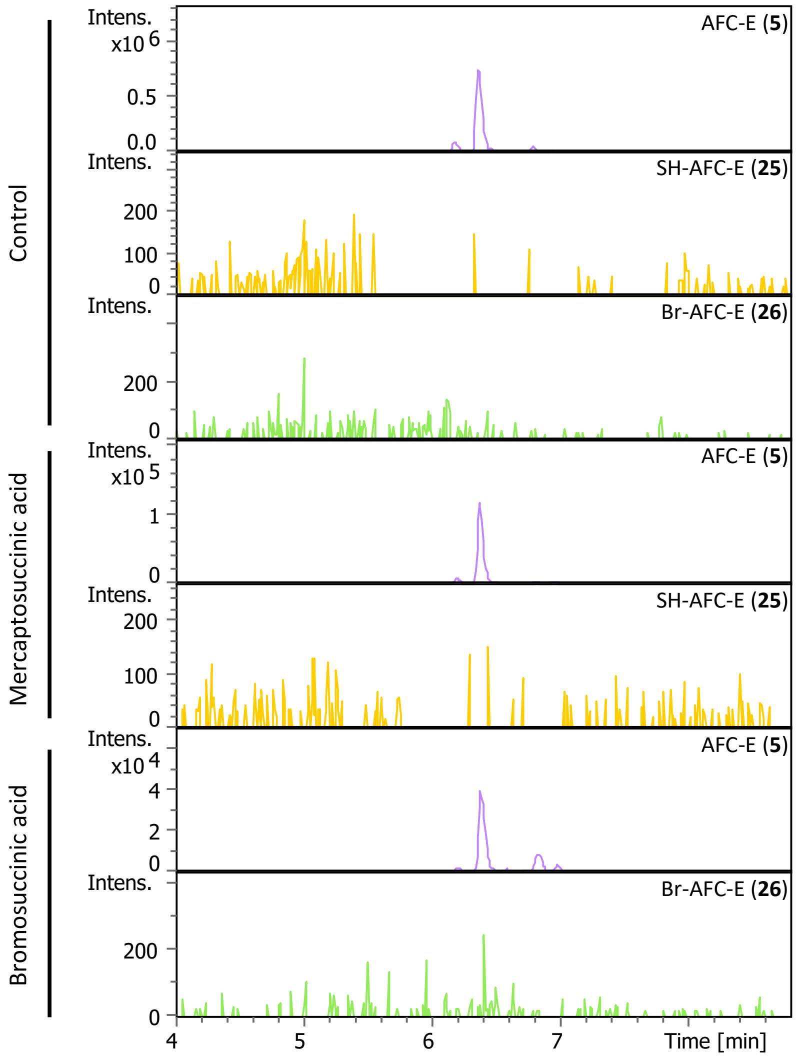


**Figure S57.** Extracted ion chromatograms from UHPLC-HR-MS showing altered AFC-E production upon supplementation with malic acid analogs. The Δ*afcS* strain culture medium was supplemented with the malic acid mimics mercaptosuccinic acid and bromosuccinic acid. No SH- or Br-substituted AFC-E derivatives were detected.

(11) Takahashi, H.; Iwasaki, A.; Ebihara, A.; Taguchi, R.; Jeelani, G.; Nozaki, T.; Suenaga, K. Isolation of hennaminal and isolation and total synthesis of hennamide, pyrrolinone

compounds from the marine *Cyanobacterium rivularia* sp. *Org. Lett.* **2023**, *25* (14), 2400–2404.
(12) Miller, V. L.; Mekalanos, J. J. A novel suicide vector and its use in construction of insertion mutations: osmoregulation of outer membrane proteins and virulence determinants in *Vibrio cholerae* requires toxR. *Journal of Bacteriology* **1988**, *170* (6), 2575–2583.
(13) Hanahan, D. Studies on transformation of *Escherichia coli* with plasmids. *Journal of Molecular Biology* **1983**, *166* (4), 557–580.
(14) Casadaban, M. J.; Cohen, S. N. Analysis of gene control signals by DNA fusion and cloning in *Escherichia coli*. *Journal of Molecular Biology* **1980**, *138* (2), 179–207.
(15) Vanlaere, E.; Baldwin, A.; Gevers, D.; Henry, D.; De Brandt, E.; LiPuma, J. J.; Mahenthiralingam, E.; Speert, D. P.; Dowson, C.; Vandamme, P. Taxon K, a complex within the *Burkholderia cepacia* complex, comprises at least two novel species, *Burkholderia contaminans* sp. nov. and *Burkholderia lata* sp. nov. *Int J Syst Evol Microbiol* **2009**, *59* (Pt 1), 102–111.
(16) Van, V. T.; Berge, O.; Balandreau, J.; Ké, S. N.; Heulin, T. Isolement et activité nitrogénasique de *Burkholderia vietnamiensis*, bactérie fixatrice d'azote associée au riz (Oryza sativa L) cultivé sur un sol sulfaté du Viêt-nam. *Agronomie* **1996**, *16* (8), 479–491.
(17) Wallner, A.; Moulin, L.; Busset, N.; Rimbault, I.; Béna, G. Genetic diversity of type 3 secretion system in *Burkholderia* s.l. and links with plant host adaptation. *Front. Microbiol.* **2021**, *12*.
(18) Brett, P. J.; DeShazer, D.; Woods, D. E. *Burkholderia thailandensis* sp. nov., a *Burkholderia pseudomallei*-like species. *Int J Syst Bacteriol* **1998**, *48 Pt 1*, 317–320.
(19) Martina, P.; Leguizamon, M.; Prieto, C. I.; Sousa, S. A.; Montanaro, P.; Draghi, W. O.; Stämmler, M.; Bettiol, M.; de Carvalho, C. C. C. R.; Palau, J.; Figoli, C.; Alvarez, F.; Benetti, S.; Lejona, S.; Vescina, C.; Ferreras, J.; Lasch, P.; Lagares, A.; Zorreguieta, A.; Leitão, J. H.; Yantorno, O. M.; Bosch, A. *Burkholderia puraquae* sp. nov., a novel species of the *Burkholderia cepacia* complex isolated from hospital settings and agricultural soils. *International Journal of Systematic and Evolutionary Microbiology* **2018**, *68* (1), 14–20.
(20) Gotschlich, A.; Huber, B.; Geisenberger, O.; Tögl, A.; Steidle, A.; Riedel, K.; Hill, P.; Tümmler, B.; Vandamme, P.; Middleton, B.; Camara, M.; Williams, P.; Hardman, A.; Eberl, L. Synthesis of multiple *N*-acylhomoserine lactones is wide-spread among the members of the *Burkholderia cepacia* complex. *Systematic and Applied Microbiology* **2001**, *24* (1), 1–14.
(21) Subramoni, S.; Agnoli, K.; Eberl, L.; Lewenza, S.; Sokol, P. A. Role of *Burkholderia cenocepacia afcE* and *afcF* genes in determining lipid-metabolism-associated phenotypes. *Microbiology* **2013**, *159* (Pt_3), 603–614.
(22) Toyofuku, M.; Cárcamo-Oyarce, G.; Yamamoto, T.; Eisenstein, F.; Hsiao, C.-C.; Kurosawa, M.; Gademann, K.; Pilhofer, M.; Nomura, N.; Eberl, L. Prophage-triggered membrane vesicle formation through peptidoglycan damage in *Bacillus subtilis*. *Nat Commun* **2017**, *8* (1), 481.
(23) Figurski, D. H.; Helinski, D. R. Replication of an origin-containing derivative of plasmid RK2 dependent on a plasmid function provided in trans. *Proceedings of the National Academy of Sciences* **1979**, *76* (4), 1648–1652.
(24) Kovach, M. E.; Elzer, P. H.; Steven Hill, D.; Robertson, G. T.; Farris, M. A.; Roop, R. M.; Peterson, K. M. Four new derivatives of the broad-host-range cloning vector pBBR1MCS, carrying different antibiotic-resistance cassettes. *Gene* **1995**, *166* (1), 175–176.
